# Anatomical, ontogenetic, and genomic homologies guide reconstructions of the teeth-to-baleen transition in mysticete whales

**DOI:** 10.1101/2022.03.10.483660

**Authors:** John Gatesy, Eric G. Ekdale, Thomas A. Deméré, Agnese Lanzetti, Jason Randall, Annalisa Berta, Joseph J. El Adli, Mark S. Springer, Michael R. McGowen

## Abstract

The transition in Mysticeti (Cetacea) from capture of individual prey using teeth to bulk filtering batches of small prey using baleen ranks among the most dramatic evolutionary transformations in mammalian history. We review phylogenetic work on the homology of mysticete feeding structures from anatomical, ontogenetic, and genomic perspectives. Six characters with key functional significance for filter-feeding behavior are mapped to cladograms based on 11 morphological datasets to reconstruct evolutionary change across the teeth-to-baleen transition. This comparative summary within a common parsimony framework reveals extensive conflicts among independent systematic efforts but also broad support for the newly named clade Kinetomenta (Aetiocetidae + Chaeomysticeti). Complementary anatomical studies using CTscans and ontogenetic series hint at commonalities between the developmental programs for teeth and baleen, lending further support for a ‘transitional chimaeric feeder’ scenario that best explains current knowledge on the transition to filter feeding. For some extant mysticetes, the ontogenetic sequence in fetal specimens recapitulates the inferred evolutionary transformation: from teeth, to teeth and baleen, to just baleen. Phylogenetic mapping of inactivating mutations reveals mutational decay of ‘dental genes’ related to enamel formation before the emergence of crown Mysticeti, while ‘baleen genes’ that were repurposed or newly derived during the evolutionary elaboration of baleen currently are poorly characterized. Review and meta-analysis of available data suggest that the teeth-to-baleen transition in Mysticeti ranks among the best characterized macroevolutionary shifts due to the diversity of data from the genome, the fossil record, comparative anatomy, and ontogeny that directly bears on this remarkable evolutionary transformation.

## Introduction

The transition in extinct stem Mysticeti (Cetacea) from tooth-assisted feeding on individual prey items to baleen-assisted filtering of small prey ranks among the most dramatic evolutionary transformations in mammalian history (Pivorunas 1979; Gatesy et al. 2013). In extant crown mysticetes (baleen whales), tooth buds are expressed and then resorbed in fetal stages as baleen tissue begins to form (Ridewood 1923; Karlsen 1962; Ishikawa and Amasaki 1995; Thewissen et al. 2017; Lanzetti 2019). In completely edentulous neonates and adults, the keratinous baleen plates are arranged in racks that continuously grow from the left and right margins of the palate (Utrecht 1965; Fudge et al. 2009; Young et al. 2015), fray through abrasive wear, and permit efficient batch-filter feeding via an array of behaviors – engulfment (’lunge’), benthic suction, and skimming (Werth 2000; Kienle et al. 2017) that are utilized singly or in combination by extant mysticetes.

The evolution of gigantic body size in this clade presumably was enabled by the ability to batch feed at low trophic levels on large aggregations of prey (Slater et al. 2017; Goldbogen and Madsen 2018). Engulfment feeding bouts of the enormous blue whale (*Balaenoptera musculus*) have been described as the world’s largest biomechanical events (Pivorunas 1979; Fig. 1e). These include high-speed lunges at batches of prey (Goldbogen et al. 2007, 2017; Shadwick et al. 2019), intake of huge volumes of prey-laden seawater into their elastic, longitudinally pleated throat pouch (Orton and Brodie 1987), expulsion of >50 tons of seawater through the baleen sieve, and retention of numerous tiny prey items in the oral cavity (Gaskin 1982; Goldbogen et al. 2017). Adult blue whales in the eastern North Pacific have been shown to consume ∼16 tonnes of krill per day on feeding days (Savoca et al. 2021). It is unlikely that this efficient, highly derived mode of prey capture evolved rapidly in a single saltatory transition, because this system requires a neomorphic keratinous sieve, numerous anatomical changes that reorganized the skull, and coordinated behavioral modifications (Lambertsen 1983; Lambertsen et al. 1995; Werth 2000).

**Fig. 1.**
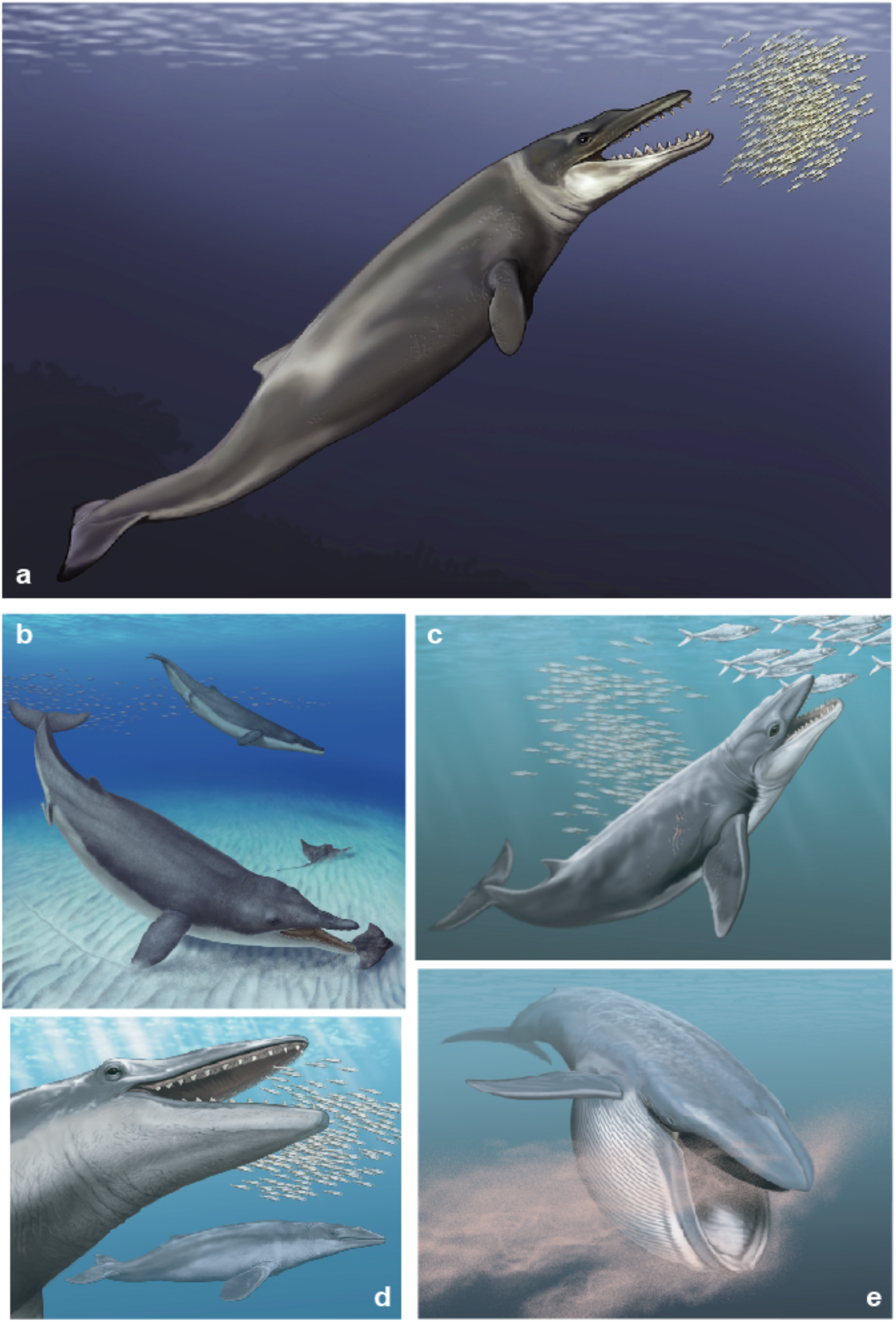
Hypothesized feeding modes in extinct toothed mysticetes (**a-d**) and batch filter feeding in extant mysticete whale (**e**). Batch-filter feeding with teeth instead of baleen, benthic suction, raptorial snapping of prey, and batch filter feeding with retention of primitive tooth-assisted feeding have been proposed as behaviors employed by Eocene and Oligocene toothed mysticetes. These modes are illustrated in *Coronodon* (**a**), *Mystacodon* (**b**), *Janjucetus* (**c**), and *Aetiocetus* (**d**), respectively. *Balaenoptera musculus*, the extant blue whale, is an obligate filter feeder (**e**), the default feeding mode for all extant mysticetes that uniformly bear baleen and lack a postnatal dentition. Illustrations are by A. Gennari (**b**) and C. Buell (**a**, **c-e**).

Despite the collection and analysis of diverse comparative data, the evolutionary sequence of change that enabled modern mysticetes to feed on large aggregations of prey remains contentious (Berta et al. 2016; Marx et al. 2016; Peredo et al. 2017). In the Eocene and Oligocene, the fossil record of Mysticeti includes diverse forms that retained functional dentitions and early representatives of Chaeomysticeti, mysticetes that are edentulous or nearly so (Fig. 1; Table 1). Representative toothed taxa include: *Coronodon havensteini* that may have filtered small prey with its closely-spaced teeth (Fig. 1a; Geisler et al. 2017), *Mystacodon selenensis* – the earliest known mysticete –interpreted as a suction feeder (Fig. 1b; Lambert et al. 2017; de Muizon et al. 2019), *Janjucetus hunderi* (Fig. 1c) a raptorial predator that likely fed on relatively large individual prey (Fitzgerald 2006), the small benthic suction feeder *Mammalodon colliveri* (Fitzgerald 2010), *Aetiocetus weltoni* and other members of the family Aetiocetidae that arguably deployed both teeth and some form of baleen to snap individual prey and filter smaller prey items (Fig. 1d; Deméré et al. 2008), and *Llanocetus denticrenatus* - an early large-bodied species with widely spaced teeth (Fordyce and Marx 2018). Extinct chaeomysticetes that fall just outside of crown clade Mysticeti also are critical for interpreting the transition to modern filter-feeding modes. Some members of the extinct Eomysticetidae show intriguing but still controversial anatomical evidence for the simultaneous expression of baleen and vestigial teeth (Boessenecker and Fordyce 2015a, b). By contrast, the recently described *Maiabalaena nesbittae* has been interpreted as a suction feeder that lacked teeth and baleen (Peredo et al. 2018), yet another contested interpretation in the field (Ekdale and Deméré 2022).

**Table 1.** Significant extinct species featured in recent phylogenetic analyses of Mysticeti

| **Extinct species** | **Family** | **Significance** | **References** |
| --- | --- | --- | --- |
| *Janjucetus hunderi*<br>(Fig. 1c) | Mammalodontidae | Oligocene toothed mysticete interpreted as a macrophagous predator | Fitzgerald (2006) |
| *Aetiocetus weltoni*<br>(Fig. 1d) | Aetiocetidae | Oligocene toothed mysticete with well-preserved lateral palatal foramina | Deméré et al. (2008); Deméré and Berta (2008); |
| *Mammalodon colliveri* | Mammalodontidae | Oligocene toothed mysticete interpreted as a benthic suction feeder | Fitzgerald (2010) |
| *Waharoa ruwhenua* | Eomysticetidae | Oligocene mysticete interpreted as a skim filter feeder with evidence for a vestigial dentition | Boessenecker and Fordyce (2015a) |
| *Tokarahia* spp. | Eomysticetidae | Oligocene mysticete interpreted as a skim filter feeder with evidence for a vestigial dentition | Boessenecker and Fordyce (2015b) |
| *Mystacodon selenensis* (Fig. 1b) | Mystacodontidae | oldest known mysticete (Eocene, ~36.4 Ma); interpreted as a benthic suction feeder | Lambert et al. (2017); de Muizon et al. (2019) |
| *Coronodon havensteini* (Fig. 1a) | unnamed clade | Oligocene mysticete interpreted as tooth-assisted filter feeder | Geisler et al. (2017) |
| *Llanocetus denticrenatus* | Llanocetidae | large-bodied Eocene toothed mysticete interpreted as a suction feeder | Fordyce and Marx (2018) |
| *Maiabalaena nesbittae* | unnamed clade | Oligocene mysticete interpreted as suction feeder with no teeth and no baleen | Peredo et al. (2018); Peredo and Pyenson (2018) |

Systematic analyses of living and extinct taxa are central to any evolutionary reconstruction of the teeth-to-baleen transition in Mysticeti. A phylogenetic hypothesis is a hierarchical map of evolutionary novelties. Genetic, behavioral, and anatomical features are used to infer phylogenetic relationships. In turn, the resulting phylogenetic hypothesis informs us on how the various characteristics of species were assembled over millions of years. Such comparative frameworks are essential for constraining evolutionary scenarios to those that are the simplest and best supported given the observed data. Without these scientific ‘rules of engagement’, wild speculations that are unnecessarily complicated can be proposed. From this perspective, systematic data matrices are ‘libraries’ of potential homologues. Cladograms derived from numerous characters then sort these primary homology statements through the critical test of congruence among characters to estimate common ancestry and the sequence of evolutionary change (Brower and Schawaroch 1996).

How diverse extinct genera (Fig. 1) branch from the stem lineage to crown Mysticeti strictly guides interpretations of functional and behavioral shifts during the transition from feeding with teeth to batch-filtering with baleen. Paleontologists have used phylogenetic analyses of living and extinct mysticetes to make detailed evolutionary maps of this macroevolutionary sequence of change. Explicitly cladistic analyses emerged in the 21st century (e.g., Geisler and Sanders 2003), with ever-expanding samplings of characters and taxa over the past 15 years (Fitzgerald 2006, 2010; Steeman 2007; Deméré et al. 2008; Marx 2010; Boessenecker and Fordyce 2015a, b; Marx and Fordyce 2015; Peredo and Uhen 2016; Geisler et al. 2017; Lambert et al. 2017; Fordyce and Marx 2018; Peredo and Pyenson 2018; Peredo et al. 2018; de Muizon et al. 2019). Simultaneously, ontogenetic and genomic data bearing on the teeth-to-baleen transition have buttressed macroevolutionary interpretations based primarily on fossils, enriching the overall historical reconstruction (e.g., Deméré et al. 2008; Meredith et al. 2009, 2011; Springer et al. 2016, 2019; Thewissen et al. 2017; Lanzetti 2019; Lanzetti et al. 2020).

Here, we review diverse phylogenetic studies of both morphology and molecules that have contributed to a deeper understanding of the complex evolutionary sequence in Mysticeti that culminated in batch filter-feeding at an awe-inspiring scale (Fig. 1e). Utilizing systematic matrices published over the past 15 years, we compare basic homology assessments, alternative character coding schemes, and contrasting evolutionary interpretations within a general parsimony framework. We map six essential morphological features related to filter feeding in modern Mysticeti onto competing phylogenetic hypotheses, contrast sequences of evolutionary change across 11 prominent studies, and review emerging ontogenetic and genomic contributions to understanding the evolutionary loss of teeth and elaboration of the neomorphic baleen sieve in mysticete whales. This overall meta-analytical comparison establishes what we consider the simplest explanatory hypothesis for the evolution of filter feeding in Mysticeti and also helps delineate what future research is required to provide further insights.

## Materials and Methods

### Phylogenetic analyses and character mapping

We reanalyzed morphological datasets from studies published between 2006 and the present that have focused on the teeth-to-baleen transition and have coded a common set of critical filter-feeding characters (Fig. 2). These 11 studies are listed in Appendix 1 with counts of taxa and characters sampled in each; studies that have built on these data matrices by adding just a few taxa and/or characters are noted but not reanalyzed here. The majority of phylogenetic work on the teeth-to-baleen transition in Mysticeti has utilized equally weighted parsimony to choose among competing tree topologies. This research provides the basic foundation for all downstream evolutionary interpretations regarding changes in anatomy, behavior, physiology, function, and molecules. We contend that it is essential to compare published work using the same basic ground-rules and generally employ a cladistic parsimony framework. However, we comment on any important differences between our parsimony reanalyses and published trees when our inferences conflict with alternative approaches utilized in the original publications (e.g., combined analysis with molecular data [de Queiroz and Gatesy 2007], Bayesian analysis [Nylander et al. 2004], or parsimony analysis with implied weighting [Goloboff 1993]).

**Fig. 2.**
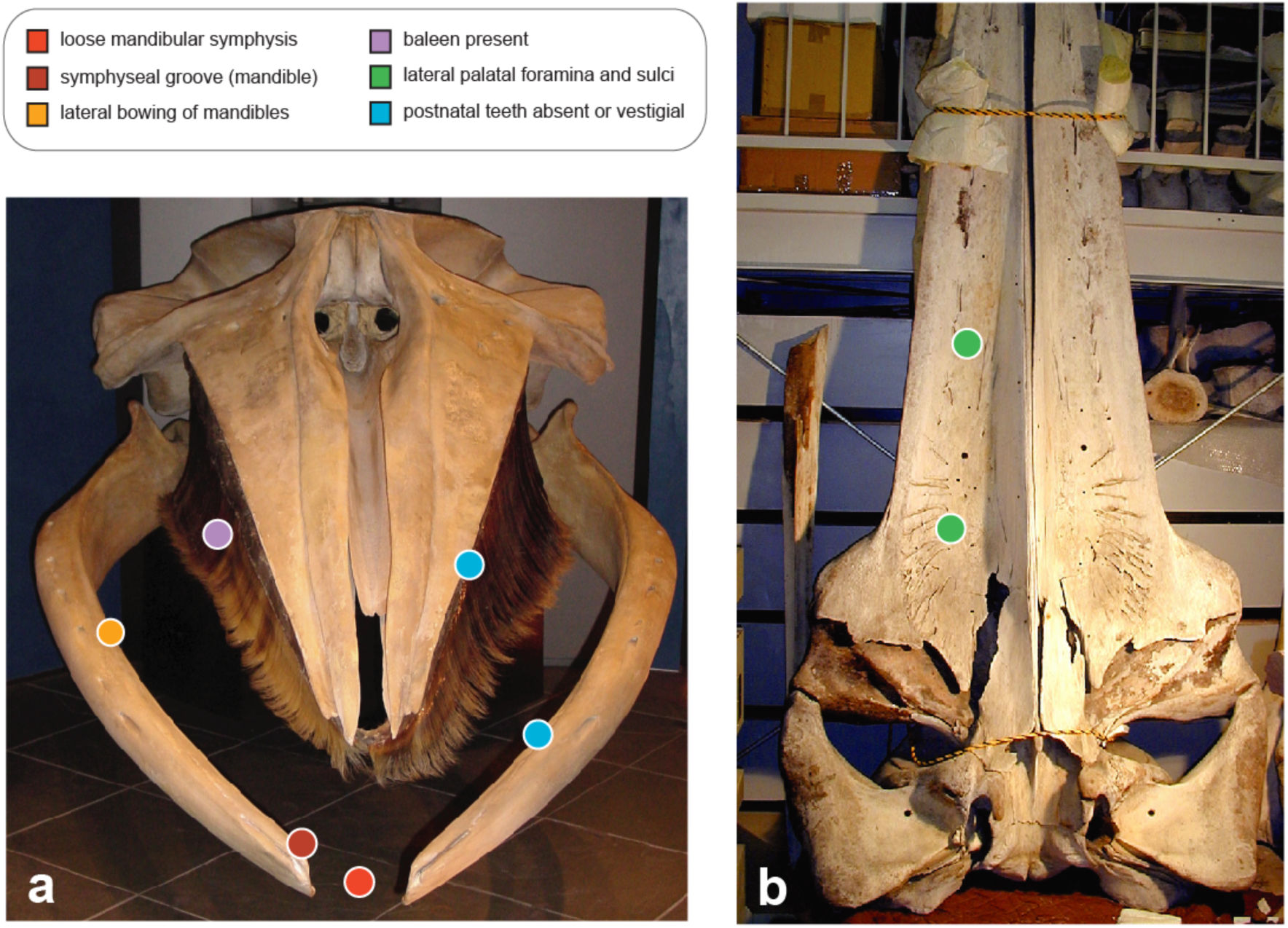
Six morphological characters related to filter feeding in modern mysticetes that were mapped onto alternative phylogenetic hypotheses: mandibular symphysis not sutured or fused (red), symphyseal groove (dark red), lateral bowing of mandibles (orange), baleen (purple), lateral palatal foramina and sulci (green), postnatal teeth absent (blue). **a.** Anterior view of *Balaenoptera acutorostrata* skull. **b.** Palatal view of *B. borealis* cranium. All extant baleen whales express these six character states that have been encoded as phylogenetic characters in most systematic studies of Mysticeti (Figs. 3, 5, 7-8; Appendix 2). Skeletal material is from National Science Museum, Tokyo (**a**) and Iwate Prefectural Museum (**b**).

Parsimony analyses of the 11 morphological data matrices were executed in PAUP* v 4.0a build 169 (Swofford 2002). All character state transformations were given equal weight, and the treatment of characters as ordered versus unordered followed the recommendations of the original authors for each data matrix. Searches were heuristic with at least 1000 random taxon addition replicates and tree-bisection-reconnection (TBR) branch swapping. To increase the efficiency of tree searches, branches with minimum length of zero were collapsed using the ‘amb-’ option of PAUP*. Strict consensus trees were constructed from all minimum length cladograms for each dataset, and these are shown in Appendix 1.

To broadly compare reconstructions of the teeth-to-baleen transition across multiple studies, we mapped character state changes on the strict consensus of minimum length cladograms for each dataset. For consistency with our phylogenetic searches, character mapping was via parsimony with equal weighting and ordering of character state changes as in tree searches. Delayed transformation (DelTran) and accelerated transformation (AccTran) parsimony optimizations were compared using PAUP*; in our figures, DelTran optimizations are shown. Six traits (Fig. 2) that can be simply coded and are key features of the filter-feeding apparatus in all extant mysticete species (see Deméré et al. 2008) were mapped onto each tree and include: 1) presence versus absence of a ‘loose’ mandibular symphysis that is not fused/sutured, 2) presence versus absence of a symphyseal groove (mandible), 3) presence versus absence of laterally bowed mandibles, 4) presence versus absence of baleen, 5) presence versus absence of lateral palatal foramina and sulci, and 6) presence versus absence of postnatal teeth. For each of the 11 reanalyzed studies, character descriptions for these six traits are given in Appendix 2. Many additional characters, such as the relative width and length of the palate, shape of the mandibular condyle, arching of the rostrum, patterns of tooth wear that imply different feeding modes, robustness/shape of the hyoids, etc. were not mapped here because these traits were commonly coded in incongruous ways across studies or were included in just a subset of studies. Thus, the present study presents a starting point for more detailed comparative analyses of the teeth-to-baleen transition in future work.

The first two characters are directly related to the unique mandibular-symphysis anatomy of modern mysticetes and were commonly coded as a single joint character in the reviewed studies (Appendix 2) as they are functionally related. The loose mandibular symphysis of mysticetes, unique among extant cetaceans, enables alpha and omega rotation of the mandibles, expansion of the oral cavity, and manipulation of the well-developed lower lips relative to the racks of baleen (Lambertsen et al. 1995; Lambertsen et al. 2005). A prominent symphyseal groove marks the attachment site for a fibrous connection between the loose mandibles that may counteract stress at the symphysis during feeding (Lambertsen et al. 2005; Fitzgerald 2012; Pyenson et al. 2012).

The third character, lateral bowing of the mandibles relative to the primitive condition, generally has been coded as multistate in published systematic work (Appendix 2). Lateral bowing of the mandibles is correlated with an increased oral cavity, and longitudinal rotation of the mandibles further increases width of the mouth in engulfment feeding mysticetes (Brodie 2001). Because this is a continuously varying character that is evolutionarily labile, optimization is often homoplastic and equivocal. In our figures, we highlight evolutionary changes from the plesiomorphic, ‘medially-bowed’ condition to ‘straightened’ mandibles that record initial lateral bowing relative to the ancestral state (see Fitzgerald 2012). Character states that describe more extreme lateral bowing of the mandibles in chaeomysticetes were merged in our figures that illustrate character optimizations (see Appendix 2). We did not map associated changes in the relative width of the palate because multiple characters in published matrices document variation in palatal width, and these characters are not consistent across different studies.

Surprisingly, several detailed systematic studies of Mysticeti did not score the fourth character, baleen (Marx and Fordyce 2015; Lambert et al. 2017; Fordyce and Marx 2018; de Muizon et al. 2019), perhaps because the presence or absence of this soft tissue character cannot be confidently coded in fossils, except when preservation is exceptional (e.g., Bisconti 2012; Marx et al. 2017). If baleen was not encoded in a morphological dataset, we assigned appropriate character states for all extant taxa, coded most extinct taxa as ‘ambiguous’ (‘?’), and scored ‘present’ for the few extinct mysticetes with fossilized baleen.

Lateral palatal foramina and sulci (character 5 above) historically have been treated as an osteological correlate for baleen (Kellogg 1965; Fordyce and de Muizon 2001; Ekdale and Deméré 2022). All extant baleen whales have these prominent nutrient foramina and grooves on the palate, albeit in alternative forms across genera (Deméré et al. 2008). These structures provide passage for vessels that nourish the ever-growing baleen plates. Despite the various phenotypic expressions of the trait in different baleen whale species, lateral palatal foramina and sulci are thought to be homologous across extant mysticetes (Ekdale et al. 2015). By contrast, the determination of which palatal structures in extinct taxa should (or should not) be encoded in systematic character matrices as potential homologues to the lateral palatal foramina in extant mysticetes is both contentious (see **Results and Discussion: Summaries of phylogenetic trees and character mappings**) and crucial for reconstructing the evolutionary history of baleen (see **Results and Discussion: The evolution of baleen and filter feeding**). The critical morphological issue for recognition of lateral palatal foramina and sulci is their anatomical position on the lateral margin of the alveolar process of the maxilla and an association via internal canals with the superior alveolar canal. Importantly, more medially placed foramina and sulci associated internally with the canal for the greater palatine artery and nerve are not related to baleen in extant mysticete species. However, CTscans or natural breaks in fossils that might reveal internal anatomy have been rare (e.g., Ekdale and Deméré 2022), so coding of this character has been controversial.

Functional mineralized postnatal teeth are completely absent in modern baleen whales, which are all obligate filter feeders. The presence versus absence of a postnatal dentition (character 6) would seem to be an easy character to code in extinct mysticetes, but there are complications due to imperfect fossil preservation. The absence of teeth in an incomplete fossil might indicate true absence or instead be due to the fact that critical regions of the jaw where teeth were present have not been preserved due to taphonomic processes (Boessenecker and Fordyce 2015a, b). Extinct crown mysticetes with incompletely preserved jaws often have been coded as edentulous even when the jaws are fragmentary. Such character codings are not based on direct observations of specimens but instead on assumptions about clade membership and the typical character state for species in that clade.

For extinct stem mysticetes, this problem of incomplete preservation is even more profound as phylogenetic bracketing based on extant species provides no clues, and the coding of character states can strongly influence phylogenetic reconstructions of the teeth-to-baleen transition. Detection of small vestigial teeth or dental alveoli is particularly problematic for incomplete fossils, and some studies have therefore coded the derived ‘absence’ state as ‘postnatal teeth absent or vestigial’ (Appendix 2). For our current analysis, we followed the original authors’ codings for presence/absence of teeth but comment on problematic codings that have implications for interpreting the teeth-to-baleen transition.

We compared character-optimization results for the feeding characters (Fig. 2) to summarize common evolutionary patterns and trends hypothesized by various research groups over the past 15 years. In particular, we focused on the relative sequence of change in the six characters along the stem mysticete lineage, determined whether homoplasy in the feeding characters was required to fit particular phylogenetic hypotheses or whether character states were uniquely derived, and noted which transformations in key characters were mapped to the same internal branch of a tree. We then used the overall cladistic framework of each previous study to evaluate functional and behavioral interpretations of the teeth-to-baleen transition that have been proposed in the literature. Throughout, we stress the importance of general criteria for assessing the homology of anatomical, molecular, and behavioral characters in a parsimony context (Patterson 1982, 1988; de Pinna 1991; Wenzel 1993; Brower and Schawaroch 1996).

In our reanalyses and discussion, we avoided published studies in which potentially interesting extinct taxa were not integrated into an explicit phylogenetic framework. For example, Marx et al. (2016) described a putative suction-feeding aetiocetid (NMV P252567) with unique tooth wear. However, despite the assertion of the authors that, “…it is confidently assigned to Aetiocetidae”, their brief list of six characters includes traits that are not synapomorphies for the clade (e.g., loose mandibular symphysis with attendant symphyseal groove, anterolaterally directed orbits), it is not clear whether Aetiocetidae is even monophyletic (Fitzgerald 2006; Boessenecker and Fordyce 2015b; Geisler et al. 2017; Fordyce and Marx 2018), and an explicit phylogenetic tree was lacking. Meaningful evolutionary analysis of the feeding apparatus in this unnamed species therefore awaits more detailed systematic work like that reviewed here.

### Inactivating mutations in tooth-specific genes

We used published literature and new analyses of genome sequences to map inactivating mutations in ten ‘dental genes’ that are tooth-specific. Inactivating mutations were procured from the literature for seven of ten tooth-specific genes (Meredith et al. 2011 [*AMEL*, *AMBN*, *ENAM*, *MMP20*]; Springer and Gatesy 2016 [*ODAPH*]; Springer et al. 2019 [*ODAM*]; Mu et al. 2021 [*ACP4*]). For the three other tooth-specific genes (*AMTN*, *DSPP*, *KLK4*), query sequences from toothed whales (*Orcinus orca* [killer whale], *Tursiops truncatus* [common bottlenose dolphin]) were BLASTed against RefSeq/WGS databases at NCBI and the Bowhead Whale Genome Resource (Keane et al. 2015) in sequence similarity searches to recover orthologous genes from five mysticetes that represent the families Balaenidae (*Balaena mysticetus* [bowhead whale], *Eubalaena japonica* [North Pacific right whale]), Eschrichtiidae (*Eschrichtius robustus* [gray whale]), and Balaenopteridae (*Balaenoptera musculus* [blue whale], *B. acutorostrata* [common minke whale]). Orthologous DNA sequences for each tooth gene were aligned in Geneious (Kearse et al. 2012) with MAFFT (Katoh et al. 2002). Alignments were visually inspected, and inactivating mutations (premature stop codons, frameshift insertions and deletions, splice site mutations at the 5’ and 3’ ends of introns, start codon mutations, complete gene deletions) were annotated. Inactivating mutations were mapped to Steeman et al.’s (2009) timetree for extant cetaceans using PAUP* with DelTran parsimony optimization (Swofford 2002).

### Homology of baleen peptides

Baleen currently is poorly characterized at the molecular level, but preliminary work on short peptides derived from baleen tissues and reference to the growing database of mysticete genome assemblies provide some insights into which genes are highly expressed in this keratinous epidermal appendage (Solazzo et al 2013, 2017). For Figure 13 (see **Results and Discussion: Repurposed keratin genes and the teeth-to-baleen shift**), we BLASTed baleen peptides from Solazzo et al. (2013) against the RefSeq/WGS databases at NCBI. Perfect matches to translations of cetacean keratin genes in this genomic database were recorded. Amino acid translations for two keratin loci (*KRT31*, *KRT36*) from the common minke whale (*B. acutorostrata*) genome were aligned using MAFFT to show representative divergence between different keratin paralogs and also to pinpoint the positions of seven perfectly-matching baleen peptides in the overall sequence alignment.

## Results and Discussion

### Summaries of phylogenetic trees and character mappings

We first provide summaries of 11 systematic studies of Mysticeti by reviewing key contributions of each work, noting critical character coding decisions that impacted evolutionary interpretations, describing evolutionary sequences for the six feeding characters that were coded across studies (Fig. 2), and documenting homoplasy (implied convergence and reversal) in these critical traits. Many of the studies focused on particularly well-preserved fossil specimens of extinct species that record unique character state combinations (Fig. 1; Table 1). So as not to misrepresent authors of the reviewed papers, we employ direct quotes from the original works to highlight controversies in reconstructing the teeth-to-baleen transition. For each study, we also include basic information on the data matrix, tree search results, and our inferred parsimony tree (strict consensus in Newick format) in Appendix 1.

#### Fitzgerald (2006)

Fitzgerald (2006) described the Oligocene toothed mysticete *Janjucetus hunderi* (Fig. 1c; Table 1) and included this taxon in an early numerical cladistic analysis of Mysticeti. His taxon/character matrix builds on the character list and codings of a more taxonomically inclusive data matrix for Neoceti (= crown Cetacea) (Geisler and Sanders 2003). Parsimony analysis (Fig. 3a) positions Oligocene aetiocetids (paraphyletic) as the toothed mysticete lineages that are most closely related to Chaeomysticeti - mysticetes that are toothless as adults (or nearly so) and generally viewed as baleen-bearing filter feeders. Fitzgerald (2006) interpreted *Janjucetus* as a large-eyed “macrophagous predator” that may have captured prey utilizing its teeth as in modern leopard seals (*Hydrurga leptonx*). This small mysticete (skull length = 46 cm) and additional Oligocene toothed mysticetes in the study (*Mammalodon*, two undescribed Charleston Museum specimens) branch from more basal positions in the overall tree, prior to the divergence of the aetiocetids (Fig. 3a).

**Fig. 3.**
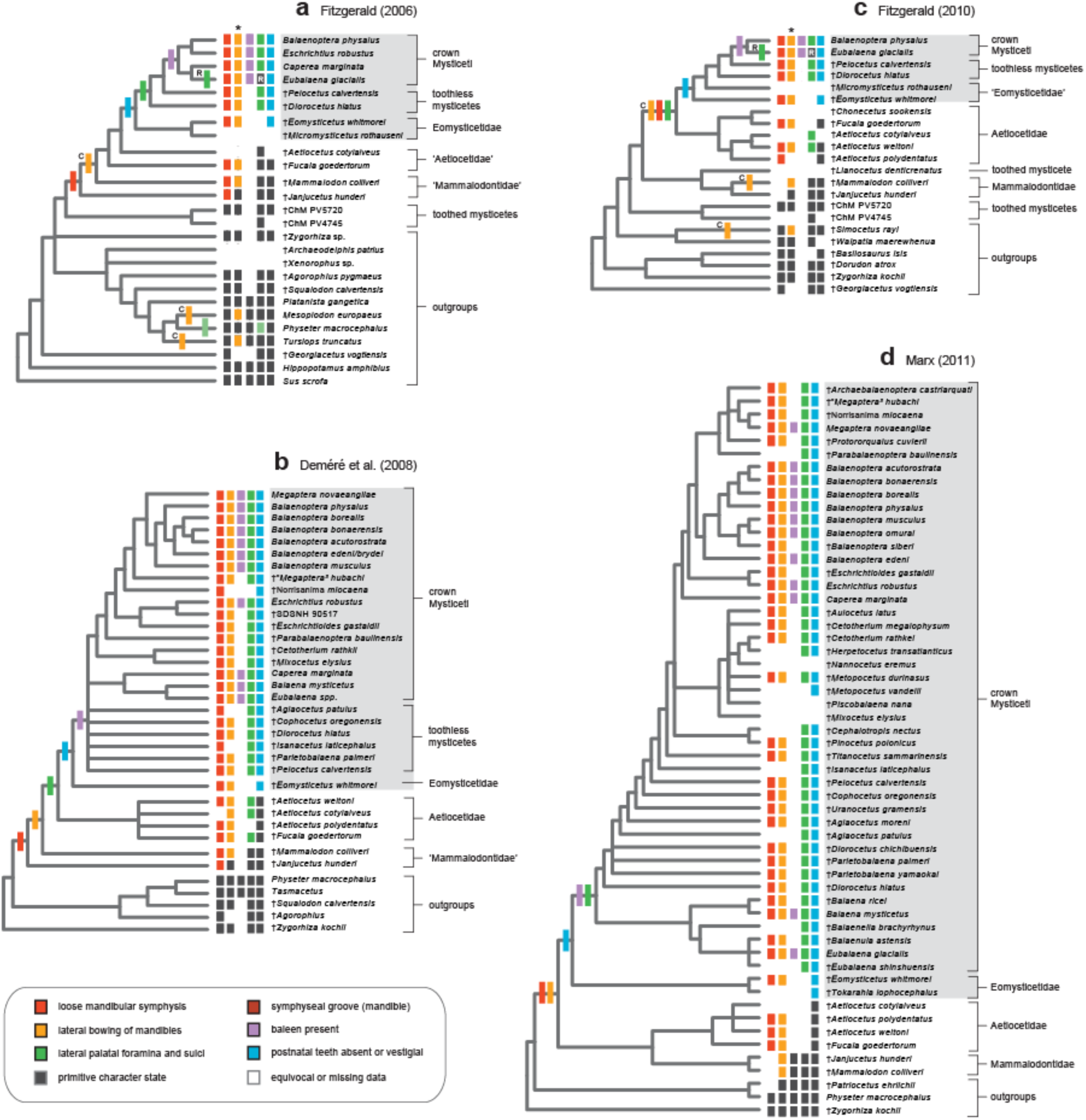
Phylogenetic interpretations of the teeth-to-baleen transition in Mysticeti (2006-2011) based on Fitzgerald (2006; **a**), Deméré et al. (2008; **b**), Fitzgerald (2010; **c**), and Marx (2011; **d**). The strict consensus of minimum length cladograms is shown for each morphological data matrix. Character state distributions are indicated for five features (dark gray rectangles = primitive state for crown Cetacea, colored rectangles = derived states, white = missing/ambiguous) with character state changes mapped onto each topology (DelTran optimization). ‘R’ indicates character reversals and reversed states, and ‘C’ indicates convergences. Asterisks (*) mark characters with alternative patterns of change for AccTran optimization (for example, convergence versus reversal). Brackets to the right of terminal taxa delimit particular clades or sets of species, and the light gray shading delineates Chaeomysticeti. Note that for Fitzgerald (2006), the palatal foramina state for *Physeter* (light green) is derived relative to the inferred primitive condition but is a different state from that shared by most extant baleen whales (darker green). See Appendix 2 for further explanation. Note that for presence versus absence of baleen, the single character state change can be mapped to many different internodes along the stem lineage to crown Mysticeti.

Filter feeding characters (Fig. 2) generally map onto the strict consensus tree in a sequential (’stepwise’) pattern. *Janjucetus* groups well within Mysticeti (Fitzgerald 2006) but lacks many of the typical filter-feeding characters of modern baleen whales. The genus instead expresses a fully developed dentition with shear and apical wear facets, dentaries that are bowed medially, and evidence for large jaw muscles that would enable feeding on large prey items. Fitzgerald noted that, “The palatal surface of the maxilla of *Janjucetus* lacks the nutrient foramina and sulci associated with vascular supply to baleen plates (Slijper 1979; Fordyce and de Muizon 2001; Deméré 2005). This is strong evidence against *Janjucetus* possessing baleen, even in an incipient form.” However, he coded *Janjucetus* as having mandibles that are “not sutured, [with] shallow longitudinal fossa for fibrocartilaginous connection” (Appendix 2). The early evolution of this trait (Fig. 3a), the first indication of lateral bowing of the mandibles at the next most apical internode (convergent with some odontocetes), and the subsequent derivation of lateral palatal foramina suggest that various toothed mysticetes expressed intermediate combinations of ‘filter-feeding’ characters. The implication is that some characters associated with batch filter feeding in extant taxa are ‘exaptations’ (Gould and Vrba 1982), traits that evolved for an initial function early in mysticete evolutionary history but were subsequently coopted for new functional roles related to batch filter feeding later in phylogenetic history. This theme of shifting functions for the same trait over evolutionary time remains a strong (and contentious) paradigm in interpretations of the teeth-to-baleen transition in Mysticeti (Marx et al. 2016; Peredo et al. 2017, 2018; Fordyce and Marx 2018; Ekdale and Deméré 2022).

For the Fitzgerald (2006) data matrix, several critical character-coding choices impact optimization of feeding characters on the strict consensus tree (Fig. 3a). First, the coding of *Janjucetus* as expressing a loose mandibular symphysis in Fitzgerald (2006) is a decision that was revised by the same author in subsequent publications to ambiguous (Fitzgerald 2010) and then to ‘sutured’ based on a new fossil specimen (Fitzgerald 2012). Likewise, the coding of this character in *Mammalodon* shifted from a loose mandibular symphysis (Fitzgerald 2006) to missing/ambiguous (?) in Fitzgerald (2010). The original coding of this character was adjusted by the author, and these changes impacted the stepwise pattern of character acquisition hypothesized in Fitzgerald (2006).

Second, because character state definitions from the broader phylogenetic study of Geisler and Sanders (2003) were repurposed to focus on mysticete relationships (Fitzgerald 2006), lateral palatal foramina in all extant mysticetes were not scored, a priori, as the same state. Instead, foramina and sulci on the palate were coded as three states that were unordered (Appendix 2). *Balaenoptera physalus* (fin whale), *Eschrichtius robustus* (gray whale), *Caperea* (pygmy right whale), and two extinct ‘cetotheres’ were coded as ‘palatal surface of maxilla bears many, large vascular foramina that open laterally and anterolaterally into long sulci’ (state 1), while *Eubalaena* (right whale) was coded as ‘palatal surface of maxilla bears few vascular foramina, those that are present are small’ (state 0), the same state as all toothed mysticetes and most outgroup taxa (Fig. 3a). *Physeter macrocephalus* (giant sperm whale) was assigned the third state (2), ‘bears numerous small vascular foramina that lack sulci’ (Appendix 2). These coding decisions imply a reversal in *Eubalaena* (state 1 to state 0) and also indicate no special similarity between the lateral palatal foramina and sulci of aetiocetids (state 0) (Deméré 2005) and most modern mysticetes (state 1). Both aetiocetid species were assigned the same primitive state as other toothed mysticetes, yet Fitzgerald (2006) then summarized the teeth-to-baleen transition by reconstructing aetiocetids as toothed mysticetes that fed with their teeth but also bore baleen for filter feeding (figure 3 in Fitzgerald [2006]), presumably based on earlier publications (e.g., Deméré 2005) and/or interpretations based on logic that was not elaborated upon. His figure 3 legend notes that, “Characters relevant to the evolution of feeding in mysticetes are optimized on to the tree at nodes where they appear”, but optimization of baleen to the common ancestor of ‘aetiocetids’ and chaeomysticetes is ambiguous given that Fitzgerald coded presence versus absence of baleen only for extant taxa in his matrix (Fig. 3a)

Note that in this phylogenetic hypothesis and all of those discussed below, the evolution of baleen, a soft tissue character, is ambiguously optimized. Because baleen generally does not preserve in fossils, extinct taxa typically are coded as missing data (?). The evolutionary transformation from baleen absent to baleen present therefore can be mapped to many different internodes along the stem lineage to crown Mysticeti with each reconstruction implying just one step in each case (i.e., multiple equally parsimonious character optimizations). Much of the controversy in reconstructing the transition to filter feeding with baleen centers on reconstructing precisely where this change occurred and justifying why one interpretation is a better explanation than alternatives (see **Results and Discussion: The evolution of baleen and filter feeding**).

#### Deméré et al. (2008)

Deméré et al. (2008) further prepared the palate of the holotype skull of *Aetiocetus weltoni* (Figs. 1d, 4; Table 1), and executed phylogenetic analyses of morphological and molecular data, including the first comparative data for mysticete ‘tooth genes’ (*ENAM*, *AMBN*, *DMP1*). Parsimony analysis of the morphological data supports a strict consensus that is largely consistent with the hypothesis of Fitzgerald (2006) as well as Deméré et al.’s (2008) tree based on combined morphological and molecular data. A monophyletic Aetiocetidae is the sister taxon of Chaeomysticeti, with the toothed mysticetes *Mammalodon* and *Janjucetus* branching from more basal nodes in the tree. Feeding characters map onto the tree in a stepwise pattern with no homoplasy (Fig. 3b).

**Fig. 4.**
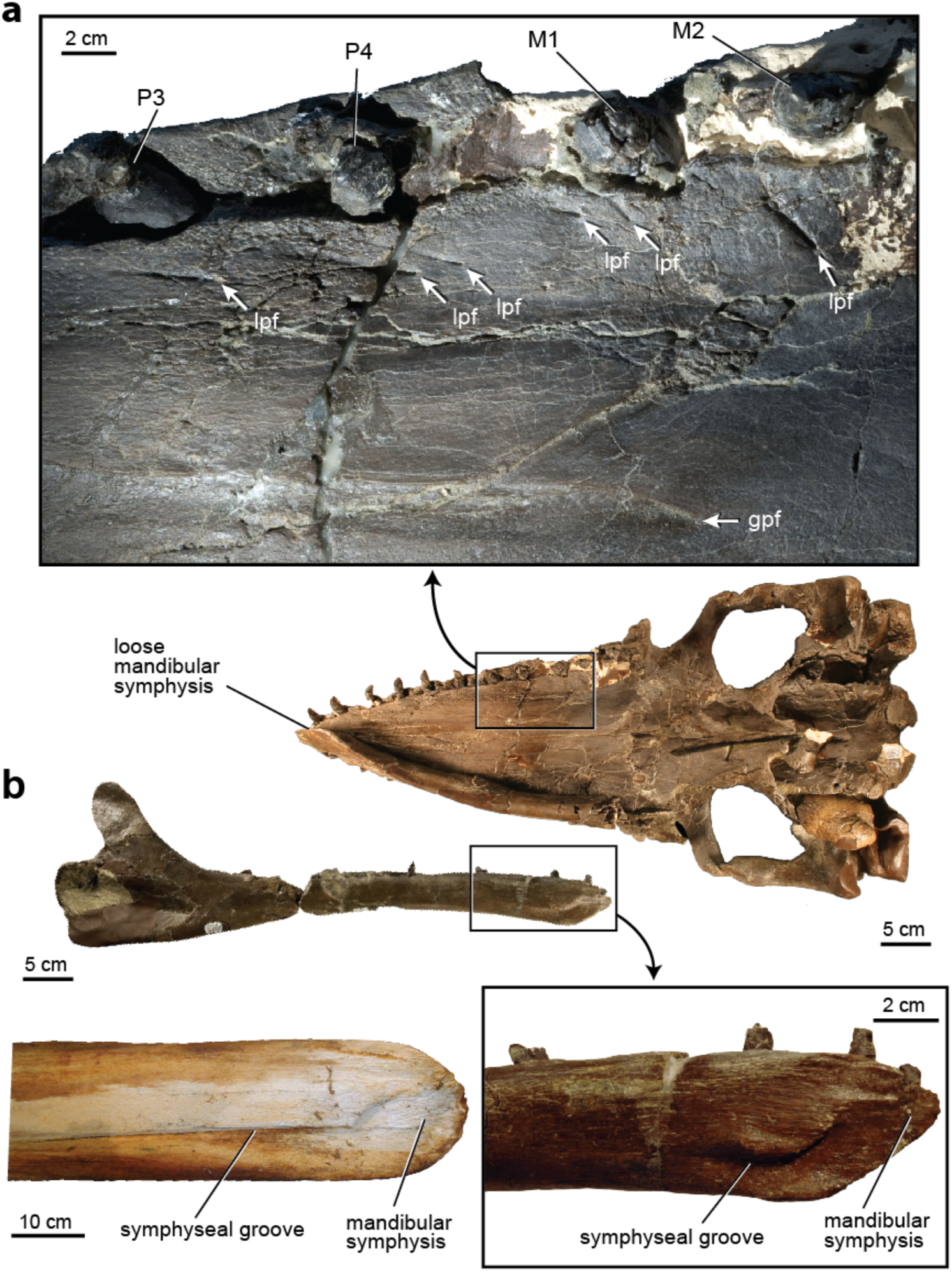
Anatomy of the Oligocene toothed mysticete, *Aetiocetus weltoni*. Skull and partial right dentary of *A. weltoni* (UCMP 122900) in ventral view (**a**). Inset shows lateral palatal foramina (lpf) medial to the tooth row and greater palatine foramen (gpf) close to the midline of the palate. Comparison of dentaries from *A. weltoni* (**b**), and extant fin whale, *Balaenoptera physalus* (**c**) shows shared derived character states. Left dentary of *A. weltoni* (UCMP 122900) is in medial view with inset of anterior end that shows similarity to the condition in *Balaenoptera physalus* (LACM 86020). The unfused, ‘loose’ symphysis and prominent groove for attachment of symphyseal ligament are synapomorphies for the clade composed of Chaeomysticeti and Aetiocetidae (= Kinetomenta). Specimens are from University of California Museum of Paleontology and the Natural History Museum of Los Angeles County.

Deméré et al. (2008; also see Deméré and Berta 2008) documented small lateral palatal foramina and sulci that are medial to the toothrow and exquisitely preserved in *A. weltoni* (Fig. 4a). Similar structures were described from several additional aetiocetids and compared to the palatal foramina and sulci of extant edentulous mysticete genera. Ultimately, they suggested homology of these structures across living and extinct species based on special similarity, relative position, and congruence tests of homology; phylogenetic analysis supports a single origin of these similar features in the common ancestor of Aetiocetidae + Chaeomysticeti (Fig. 3b). Deméré et al. (2008) argued that teeth and some form of baleen, even if only simple bristles, likely occurred simultaneously in aetiocetid mysticetes. This hypothesis minimizes changes in function and morphology on the most parsimonious trees, and assumes that the primary function of the lateral palatal foramina in extant mysticetes is to supply blood and innervation to the constantly growing baleen apparatus (also see Ekdale and Deméré 2022).

In combination with limited lateral bowing of the mandibles relative to the primitive state and a mobile mandibular symphysis, these small-bodied aetiocetids (*A. weltoni* skull length = 62 cm) express a relatively broad palate with relatively thin lateral margins, additional features sometimes associated with filter-feeding and expansion of oral volume, but with retention of a full adult dentition (Figs. 1d, 4). Deméré et al. (2008) suggested that aetiocetids are ‘mosaic’ intermediates that could have used both primitive (tooth-assisted) and derived (baleen-assisted) feeding behaviors to forage at multiple levels of a food chain, concluding that, “… by extending the size range of prey that could be captured efficiently, the composite feeding anatomy of aetiocetids may have facilitated the transition from ancestral toothed forms to derived filter-feeders that are toothless.” Their overall phylogenetic hypothesis suffers, however, from utilization of inaccurate character codes for the mandibular symphysis taken from Fitzgerald (2006). These errors were later corrected in the supermatrix analysis in Gatesy et al. (2013), and the original tree topology from Deméré et al. (2008) was still best supported.

#### Fitzgerald (2010)

Fitzgerald (2010) focused on detailed description and phylogenetic placement of the toothed mysticete *Mammalodon colliveri*, a bizarre small-bodied late Oligocene species of Australasia (Table 1). Parsimony analysis of the morphological data matrix once again positions the toothed mysticete clade Aetiocetidae sister to Chaeomysticeti, but *Janjucetus* and *Mammalodon* (Mammalodontidae) group with the large-bodied late Late Eocene *Llanocetus denticrenatus* from Antarctica (Mitchell 1989) in the single minimum length tree (Fig. 3c). Previous character codes for the mandibular symphysis in *Mammalodon* and *Janjucetus* were adjusted (now coded as ‘?’), and the coding of palatal foramina and sulci also was altered relative to Fitzgerald (2006; Fig. 3a). Fitzgerald (2010) scored the aetiocetid *Fucaia goedertorum* as missing/ambiguous (contra Deméré et al. 2008) for the presence/absence of lateral palatal foramina and grooves, but the homology of palatal nutrient foramina in two other aetiocetids and chaeomysticetes was supported by parsimony analysis. *Eubalaena glacialis* (North Atlantic right whale) was coded as ‘absent’ for this character, resulting in an inferred reversal on the tree. Lateral bowing of the mandibles relative to the primitive condition also was characterized by homoplasy, both within Mysticeti and convergently between some mysticetes and the stem odontocete *Simocetus rayi* (Fig. 3c).

Fitzgerald (2010) developed initial arguments for the importance of suction feeding in toothed mysticetes, a theme that subsequently gained strong momentum (e.g., Marx 2010; Lambert et al. 2017; Peredo et al. 2018; Fordyce and Marx 2018). In particular, Fitzgerald (2010) interpreted *Mammalodon* as a suction feeder based on tooth wear and a unique rostral morphology, suggesting an early diversification of feeding modes in Mysticeti (suction feeding, snapping of small prey, macrophagous ripping, batch filter feeding). The possibility that suction feeding was the initial impetus for the evolution of baleen-assisted batch feeding was explored, drawing on earlier, more speculative work (Darwin 1872; Werth 2000). Because baleen is an ever-growing keratinous structure, wear from the abrasive effects of sediment can be accommodated (Werth et al. 2020), which could have been a selective advantage relative to teeth that simply wear away and are not replaced, as seen in the holotype skull of *Mammalodon colliveri* (NMV P199986) (Fitzgerald 2010) as well as in the earliest described mysticete, *Mystacodon selenensis,* another putative benthic suction feeder (Lambert et al. 2017; de Muizon et al. 2019).

#### Marx (2011)

Marx (2011) assembled a large morphological data matrix that significantly extended taxonomic sampling of crown mysticetes relative to previous work. Equally weighted parsimony (Fig. 3d), parsimony with Goloboff weighting, and Bayesian analysis support the same branching sequence at the base of Mysticeti. In contrast to previous cladistic analyses, aetiocetids were not sister to chaeomysticetes. Instead, a monophyletic Aetiocetidae groups with Mammalodontidae (*Janjucetus* + *Mammalodon*) to form a diverse clade of Oligocene stem toothed mysticetes that is sister to Chaeomysticeti (Fig. 3d).

Feeding characters map to the tree without homoplasy and in a sequential pattern (for DelTran optimization) because several characters were assigned different codes relative to earlier studies. For shape of the mandibles, *Janjucetus* had previously been scored as ‘bowed medially’, the state that matches most outgroup taxa and is presumably plesiomorphic (Fig. 3a-c). Marx (2011) instead coded the mandibles in *Janjucetus* as ‘straight’ (Fig. 3d; Appendix 2). This discrepancy likely reflects an ambiguous anatomical condition in *Janjucetus* that is at the margin between two different character state definitions. In other cases, more controversial decisions were made. Most importantly, lateral palatal foramina and sulci were coded as ‘ambiguous’ for all aetiocetids. Marx (2011) explained this cautious choice, noting that the homology of lateral palatal foramina with chaeomysticetes “…has been questioned (Ichishima et al. 2008), as has the presence of these grooves in some of the taxa in which they were originally proposed to occur (Fitzgerald 2011). Because of this uncertainty, this character was coded as uncertain (‘?’) for the toothed mysticetes in question.” Coding aetiocetids as ‘uncertain’ as in Max (2011) could be justified on the grounds that lateral palatal foramina and sulci show a range of forms in both living and extinct crown mysticetes and that these features are relatively smaller in aetiocetids (Deméré et al. 2008), drawing doubt in some researchers as to whether similarity is compelling enough to make an initial hypothesis of homology, or not.

The presence versus absence of baleen also was coded as ‘uncertain’ in aetiocetids (Fig. 3d) but was coded as ‘absent’ (state ‘0’) in extinct outgroups and two extinct toothed mysticetes (*Janjucetus* and *Mammalodon*) that completely lack foramina and sulci on the lateral margins of the palate. These coding decisions are not justified, given that it is unknown whether incipient ‘proto-baleen’ was developed in these Oligocene toothed mysticetes prior to the evolution of derived canals that, in extant mysticetes, enhance blood flow and provide passage for innervation to the baleen plates. Given the poor fossilization potential of baleen, absence cannot be reliably coded in extinct taxa. However, this character-coding choice does not impact minimum tree length (still one step) and just refines the set of internodes at which baleen is inferred to have evolved in the tree (Fig. 3d).

The initial codings of anatomical structures are where valid disagreements among different systematists commonly are articulated. In this particular case, Deméré et al. (2008) and Deméré and Berta (2008) based their primary homology decisions for putative lateral palatal foramina and sulci in aetiocetids on relative position (i.e., the structures are where they should be on the lateral palate) and on detailed similarity (i.e., the structures are reminiscent of scaled-down lateral palatal foramina and sulci of modern balaenopterids); these details of similarity were later augmented by even more anatomical specificity using CTscans (Ekdale et al. 2017; Ekdale and Deméré 2022). The uncertainty codings in Marx (2011) potentially impact the choice among alternative trees, because presence of lateral palatal foramina and sulci has been interpreted as a synapomorphy for the Aetiocetidae + Chaeomysticeti clade in prior cladistic analyses (Fig. 3b, C).

This grouping of Aetiocetidae + Chaeomysticeti is contradicted by parsimony analysis of Marx’s (2011) data matrix, but recoding lateral palatal foramina and sulci as present in aetiocetids (as opposed to ambiguous ?) and redoing the parsimony analysis still does not support this clade, because multiple characters in the matrix group Aetiocetidae with Mammalodontidae to the exclusion of Chaeomysticeti (Fig. 3d). Marx (2011) listed six synapomorphies for this novel clade of Oligocene toothed mysticetes, “…the presence of a palatal window [exposing the vomer] (char. 19), large (char. 27) and deeply notched (char. 35) orbits, a short anterior border of the supraorbital process of the frontal (char. 29) bordered by the lacrimal only (char. 32), and a supraorbital process with a distinctly shortened medial portion (char. 33).” Five of the six characters are related to the orbits and orbital rim, with multiple characters that characterize the shape of the supraorbital process of the frontal. Some of these ‘orbital characters’ could be correlated developmentally due to their close proximity in the skull and potentially are also functionally related, given that the sampled aetiocetids and mammalodontids are all small-bodied with large eyes that are anteriorly directed.

Indeed, a major narrative in Marx (2011) is that the toothed mysticete clade (Aetiocetidae + Mammalodontidae) is defined by adaptive features related to active vision-based predation. He noted that, “…large and forward-pointing eyes are found in all aetiocetids, as well as *Janjucetus hunderi* and *Mammalodon colliveri* (figure 5). In this respect, toothed mysticetes clearly differ from all extinct and extant mysticetes, instead resembling pinnipedimorphs.” Marx (2011) noted that, if accurate, the rearranged relationships at the base of Mysticeti potentially overturn previous interpretations of aetiocetids as ‘transitional taxa’ that record a unique ancestral combination of traits in the transformation to obligate batch filter feeding with baleen (Deméré and Berta 2008; Deméré et al. 2008). Thus, in Marx’s (2011) tree, aetiocetids instead are interpreted as a dead-end side-branch that convergently evolved small lateral palatal foramina (Fig. 4a). Given the ambiguity in character codes for mammalodontids, the tree topology implies that a loose mandibular symphysis is a more general character that evolved deeper in mysticete evolutionary history. The phylogenetic results could “…imply the absence of baleen in *both* mammalodontids and aetiocetids” (Marx 2011). Indeed, for this data matrix, the most parsimonious interpretation requires no homoplasy for presence versus absence of baleen, with a single origin of baleen on the stem lineage of Chaeomysticeti. The new tree (Fig. 3d) fueled the perspective developed in subsequent papers that suction-feeding without baleen, not the simultaneous presence of both teeth and baleen, defines the critical transitional anatomy on the path to edentulous batch feeding with just baleen.

**Fig. 5.**
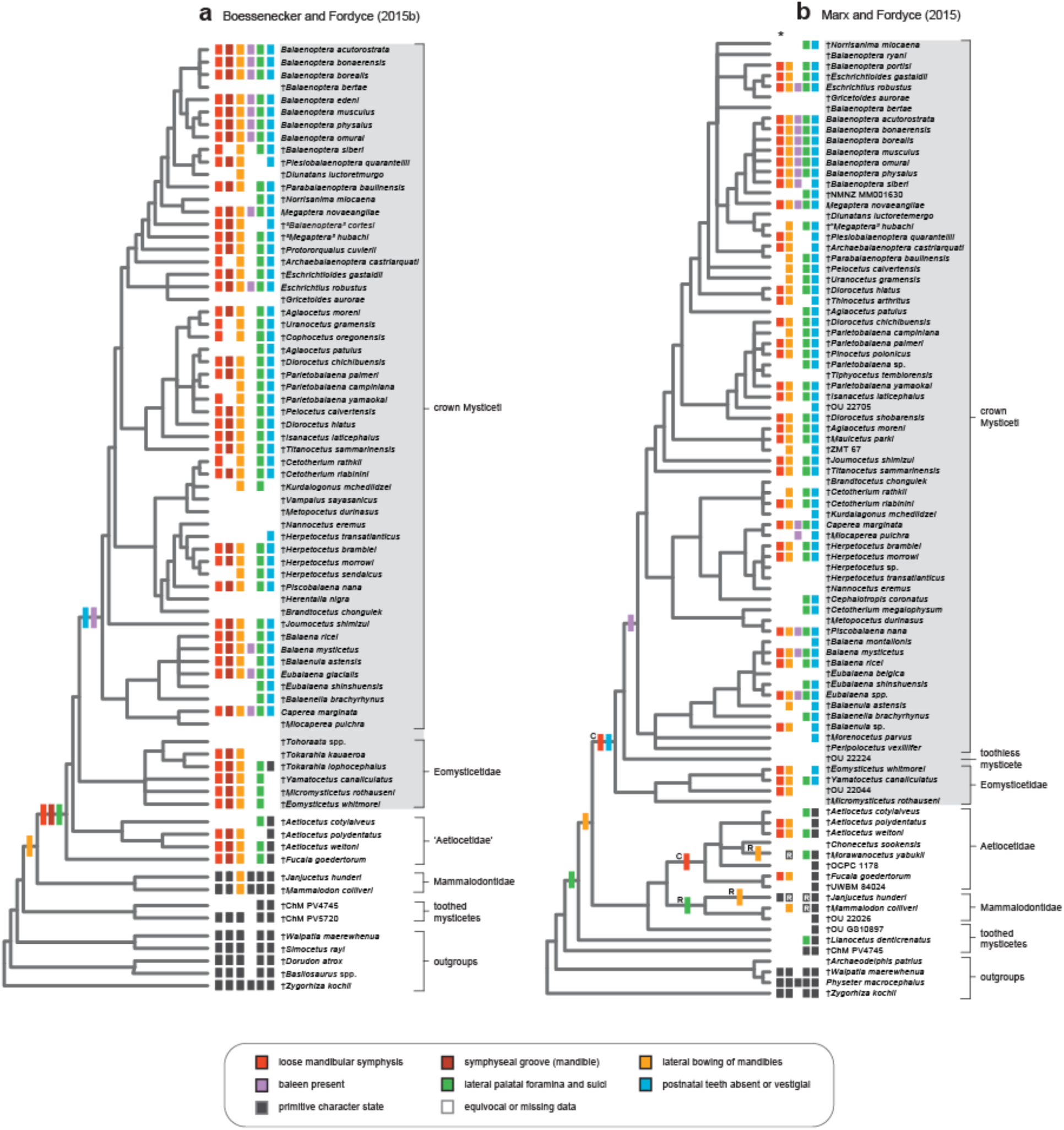
Phylogenetic interpretations of the teeth-to-baleen transition in Mysticeti (2015) based on Boessenecker and Fordyce (2015b; **a**) and Marx and Fordyce (2015; **b**). The strict consensus of minimum length cladograms is shown for each morphological data matrix. See Figure 3 legend for additional explanation/information relevant to interpretation of trees.

#### Boessenecker and Fordyce (2015b)

Boessenecker and Fordyce (2015b; also see 2015a) focused on detailed description of several eomysticetid fossils (Figs. 5a, 6; Table 1) and developed a new, comprehensive character/taxon matrix to explore the phylogenetic placement of these Oligocene stem mysticetes. Character and taxon sampling was broad, but the selection of toothed mysticetes was not expanded relative to prior work. In earlier systematic studies, the type eomysticetids from the western North Atlantic were interpreted as edentulous, baleen-bearing chaeomysticetes that were batch filter feeders (Fig. 3; Sanders and Barnes 2002; but see Okazaki 2012). The new eomysticetids from New Zealand described by Boessenecker and Fordyce (2015a, b) preserve lateral palatal foramina and sulci on the posterior two thirds of the elongated palate and evidence for vestigial teeth at the tips of the upper and lower jaws (Fig. 6a-c), suggesting an intermediate morphology with baleen posteriorly and teeth anteriorly (Fig. 6d). Based on these fossils, Boessenecker and Fordyce (2015a, b) interpreted eomysticetids as batch filter feeders that likely skimmed or perhaps gulped aggregations of small prey. Evidence for the joint expression of teeth and baleen in these early chaeomysticetes implies that the complete resorption and loss of teeth is not required for baleen to develop, an idea based on dubious extrapolations from ontogenetic series of extant baleen whales – highly derived species that have completely lost postnatal teeth (Peredo et al. 2017; see **Results and Discussion: Ontogeny recapitulates phylogeny in the teeth-to-baleen transition?**).

**Fig. 6.**
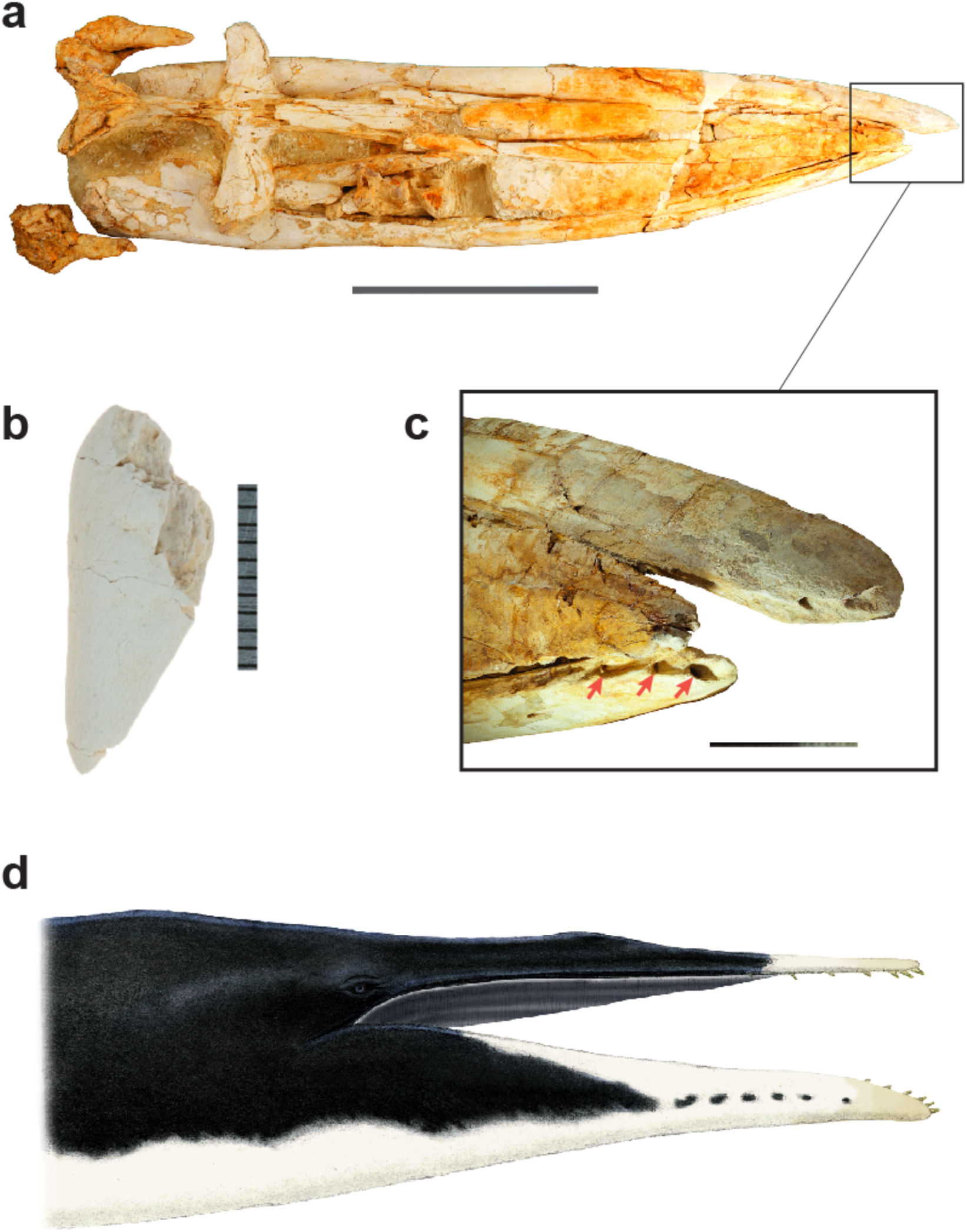
Evidence for the simultaneous occurrence of teeth and baleen in eomysticetids. Dorsal view of holotype cranium (OU 22044) of *Waharoa ruwhenua* is shown in **a** with focus on tip of the snout (**c**) that highlights lower incisor alveoli on the right dentary (red arrows). An isolated partial tooth closely associated with skull of the eomysticetid *Tokarahia* sp., cf. *T. lophocephalus* (OU 22081; Boessenecker and Fordyce 2015b) is shown in **b**. Scale bars are 50 cm (**a**), 1 cm (**b**), and 10cm (**c**). A reconstruction of *Waharoa ruwhenua* shows putative baleen and vestigial teeth (**d**). Fossils preserve lateral palatal foramina and grooves on the posterior two thirds of the palate as well as alveoli on the premaxilla, maxilla, and mandible of this species (Boessenecker and Fordyce 2015a). Photos and painting are by R. Boessenecker. Fossil material is from Geology Museum, University of Otago.

Boessenecker and Fordyce’s (2015b) coding decisions for several key characters diverged from previous systematic work (Fig. 3). Fewer feeding characters were scored as ambiguous (‘?’) in extinct stem mysticetes. For example, *Mammalodon* and *Janjucetus* were scored as ‘absent’ for both derived mandibular symphysis traits, in part based on description of new fossils (Fitzgerald 2012), and multiple eomysticetids and aetiocetids were scored as ‘present’ for lateral palatal foramina and sulci (Fig. 5a). Furthermore, no extant outgroups were included in the data matrix, so three extinct taxa that lack lateral palatal foramina (*Mammalodon*, *Janjucetus*, *Zygorhiza*) were coded ‘absent’ for presence versus absence of baleen, even though the absence of lateral palatal foramina does not unequivocally deny the presence of primordial baleen.

Parsimony analysis of Boessenecker and Fordyce’s (2015b) data matrix supports ‘aetiocetids’ (paraphyletic) as the closest fully-toothed relatives of Chaeomysticeti, and the strict consensus implies a sequential acquisition of ‘filter-feeding’ characters. Undescribed Charleston Museum stem toothed mysticetes (ChM PV4745, ChM PV5720) group in an early diverging clade that is sister to remaining mysticetes that were sampled in the study. The complete evolutionary loss of postnatal teeth can be delineated more precisely relative to previous systematic studies. Absence of postnatal teeth maps to the base of crown mysticetes, postdating the divergence of eomysticetids (Fig. 5a). This interpretation of character evolution assumes that eomysticetids did indeed have vestigial teeth (Fig. 6). A loose mandibular symphysis and lateral palatal foramina are interpreted as synapomorphies for an “Aetiocetidae” + Chaeomysticeti clade (Fig. 4). Boessenecker and Fordyce (2015b) described the unique morphology of the mandibular symphysis in mysticetes using two characters instead of just one. Their first character records the nature of the symphysis as either ‘fused/sutured’ or ‘not sutured’, and the second character records ‘presence’ or ‘absence’ of a symphyseal groove. Both mandibular symphysis characters map cleanly to the common ancestor of “Aetiocetidae” + Chaeomysticeti. This coding choice provides even more character support for this key relationship in the teeth-to-baleen transition (Fig. 5a). In studies where these two characters are distinguished, they are always associated and presumably are functionally related in all extant mysticetes (Lambertsen et al. 2005, 2015).

#### Marx and Fordyce (2015)

Marx and Fordyce (2015) expanded on Marx’s (2011) dataset and sampled many additional extinct stem mysticetes. The study is notable for its detailed archiving and photo-documentation of morphological character states (www.morphobank.org, project 687) and for comprehensive registry of stratigraphic ranges for all extinct taxa in the dataset. Morphological characters for living and extinct taxa were merged with the molecular supermatrix of McGowen et al. (2009). Marx and Fordyce (2015) coded ‘palatal nutrient foramina and sulci’ as ‘present’ in three aetiocetids instead of ‘uncertain’ as in Marx (2011). They also coded the Eocene toothed mysticete *Llanocetus*, the oldest known mysticete at the time, as ‘present’ for this character (Fig. 5b). In contrast to Marx (2011), the mandibular symphysis in *Janjucetus hunderi* was coded as ‘sutured or fused’ instead of ‘missing/ambiguous’ (Fig. 3d), and mandibular shape was changed from ‘straight or slightly bowed laterally’ to ‘medially bowed’ (Fitzgerald 2012; Marx and Fordyce 2015).

The authors’ combined Bayesian timetree is largely congruent with our parsimony analysis of just the morphological characters, but for relationships of extinct basal lineages, weakly supported placements of several taxa in the combined Bayesian timetree were not resolved in the parsimony strict consensus tree. In particular, *Llanocetus* was weakly supported as sister to Chaeomysticeti in Bayesian analysis (with low posterior probability of 0.58) but assumes a more basal position in the parsimony tree (Fig. 5b). Given that this dataset is an expanded version of Marx (2011), it is not surprising that the same ‘large-eyed clade’ of toothed mysticetes composed of aetiocetids and mammalodontids is once again supported in both parsimony (Fig. 5b) and Bayesian (Marx and Fordyce 2015) trees by a suite of characters mostly associated with the orbits and orbital rim.

It is noteworthy, that with adjusted character coding and more complete taxon sampling relative to Marx (2011), the strict consensus of minimum length cladograms now implies extensive homoplasy in characters associated with filter feeding in modern mysticetes. Lateral palatal foramina and sulci, lateral bowing of the mandibles, and a loose mandibular symphysis each require two or more changes on the parsimony strict consensus tree with many reversals and convergences (Fig. 5b). This pattern of widespread homoplasy is a striking contrast to the homoplasy-free mapping of essentially the same characters on the consensus tree for Boessenecker and Fordyce (2015b) that was published in the same year (Fig. 5a). The conflicting topologies highlight how particular character coding choices, as well as differences in taxon sampling can generate contradictory interpretations of mysticete phylogeny and the teeth-to-baleen transition, even for datasets assembled by overlapping sets of authors.

#### Lambert et al. (2017)

Lambert et al. (2017) provided the initial description of the oldest known mysticete, *Mystacodon selenensis* (Fig. 1b; Table 1), from the late Eocene (∼36.4 Ma) of Peru. They placed this early diverging toothed species into a phylogenetic context using a modified version of Marx and Fordyce’s (2015) morphological data matrix. The articulated partial skeleton of the type specimen (MUSM 1917) preserves the skull and parts of the pectoral girdle, forelimb, vertebral column, and pelvis. A highly worn dentition and other features of the skull suggest that *Mystacodon* may have been a benthic suction feeder.

Parsimony analysis of Lambert et al.’s (2017) dataset positions *Mystacodon* as sister to all remaining mysticetes (Fig. 7a). The authors interpreted this topology as indicating an initial behavioral shift in the mysticete lineage from raptorial feeding as in ‘archaeocetes’ to benthic suction feeding on single, small prey items. The strict consensus tree does not resolve a monophyletic Chaeomysticeti. The phylogenetic position of the stem toothed mysticete, ChM PV5720, disrupts this clade and implies that postnatal teeth were lost twice within Mysticeti or were lost and then re-evolved in ChM PV5720. In addition to this incongruence with other studies (Figs. 3, 5, 7-8), three other characters related to filter feeding in mysticetes (lateral palatal foramina, loose mandibular symphysis, lateral bowing of mandibles) are homoplastic (Fig. 7a). Because character codes were recycled from Marx (2011) and also Marx and Fordyce (2015), the same Aetiocetidae + Mammalodontidae clade is supported by the same suite of characters related to the orbits in Lambert et al. (2017). *Llanocetus* groups with chaeomysticetes and ChM PV5720, which results in lateral palatal foramina and sulci optimizing as a synapomorphy for this combined clade with a reversal in ChM PV5720 as well as a convergence with aetiocetids. Overall, the parsimony tree for this dataset supports an extremely complicated pattern of evolution for the feeding characters (Fig. 2), with Aetiocetidae only distantly related to extant baleen whales.

**Fig. 7.**
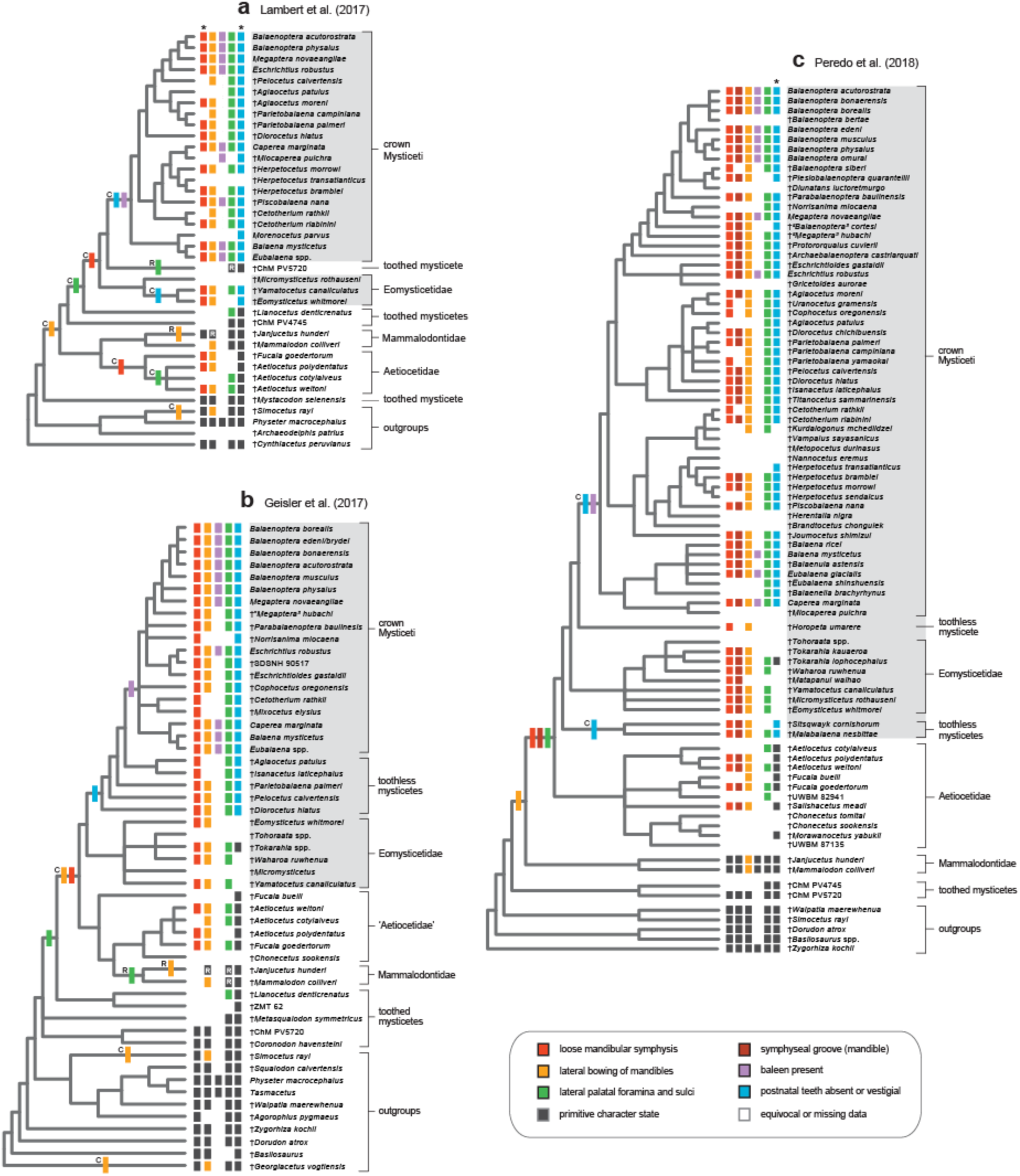
Phylogenetic interpretations of the teeth-to-baleen transition in Mysticeti (2017-2018) based on Lambert et al. (2017; **a**), Geisler et al. (2017; **b**), and Peredo et al. (2018; **c**). The strict consensus of minimum length cladograms is shown for each morphological data matrix. See Figure 3 legend for additional explanation/information relevant to interpretation of trees. Note that for presence versus absence of baleen, a single origin of baleen can be mapped to many different internodes along the stem lineage to crown Mysticeti due to missing data. DelTran optimizations are shown. For (**c**), the DelTran optimization in (**c)** is consistent with baleen evolving after the divergence of *Maiabalaena* + *Sitsqwayk* from other chaeomysticetes (see Peredo et al. 2018). Given that lateral palatal foramina and sulci map to the common ancestor of Aetiocetidae + Chaeomysticeti, however, an alternative interpretation is that baleen also was present in the common ancestor of this clade, and there was an autapomorphic loss of connections between the lateral palatal foramina and the superior alveolar canal in *Maiabalaena* (see main text). In this same tree (**c**), the convergent loss of postnatal teeth shown in the DelTran optimization is ambiguous; loss of teeth in the common ancestor of Chaeomysticeti and a reversal in Eomystidae is equally parsimonious.

**Fig. 8.**
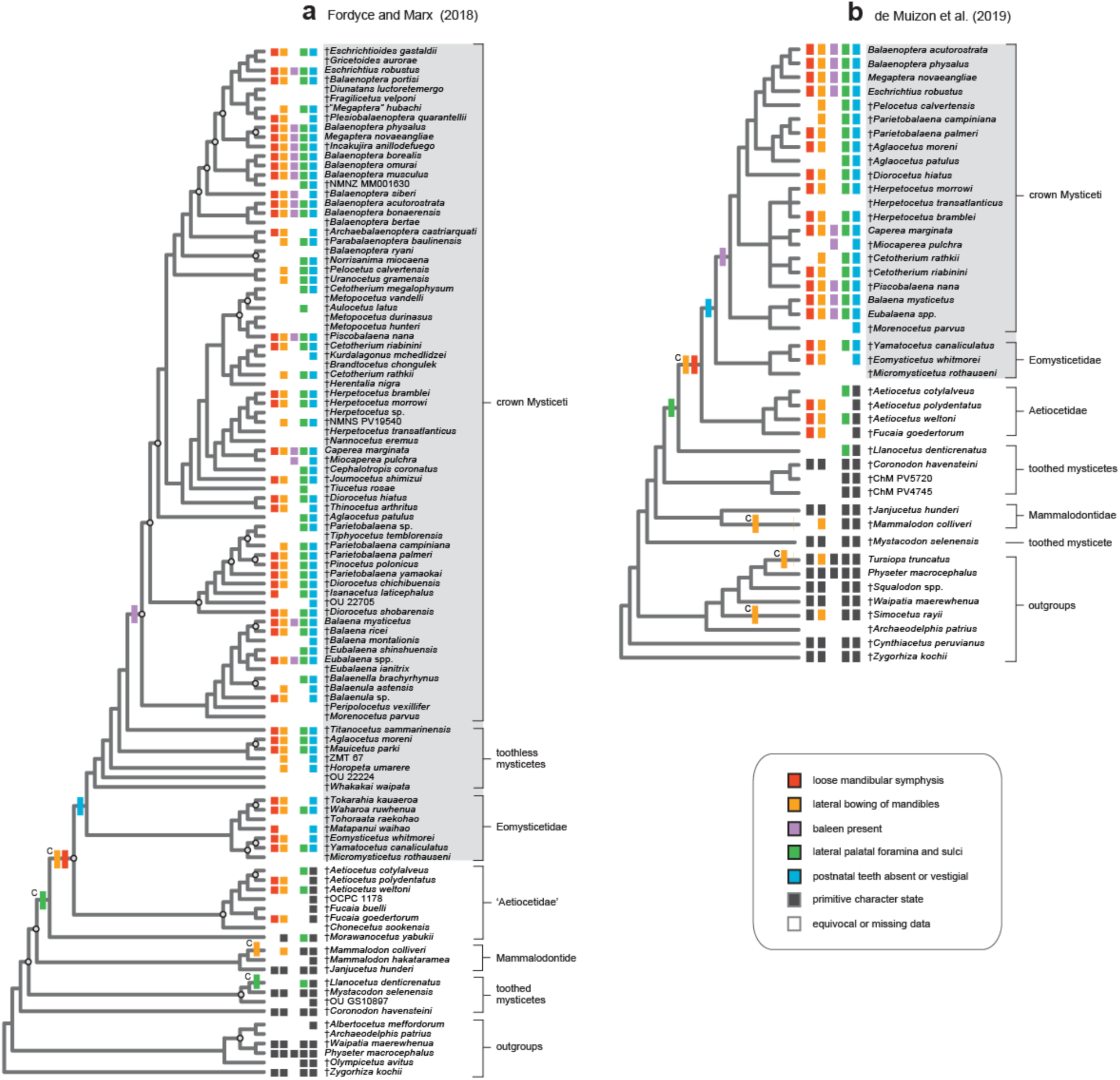
Phylogenetic interpretations of the teeth-to-baleen transition in Mysticeti (2018-2019) based on Fordyce and Marx (2018; **a**) and de Muizon et al. (2019; **b**). The Bayesian consensus for combined DNA and morphological data is shown for **a**, and the strict consensus of minimum length cladograms based on the morphological matrix is shown in **b**. Circles at nodes in the Bayesian tree mark clades with very low support (posterior probabilities < 0.50). The parsimony strict consensus tree for the morphological matrix of Fordyce and Marx (2018) is given in Appendix 1; this tree also positions aetiocetids as the closest fully-toothed relatives of Chaeomysticeti. See Figure 3 legend for additional explanation/information relevant to interpretation of trees.

#### Geisler et al. (2017)

Geisler et al. (2017) described *Coronodon havensteini* (Fig. 1a; Table 1), an Oligocene mysticete with broad, partially overlapping, multicusped teeth and used phylogenetic analyses of a large supermatrix of morphological and molecular characters to place this intriguing species. Based on morphology of the skull and teeth, as well as specific tooth-wear and abrasion patterns, the authors hypothesized that *Coronodon*’s unique dentition enabled raptorial or suction capture of single prey items and also batch filter feeding on smaller prey, without the aid of a baleen filter. This scenario of tooth-aided filter feeding harkens back to much earlier speculations on possible pathways for the teeth-to-baleen transition in Mysticeti (Mitchell 1989; Fordyce 1984; Fordyce and de Muizon 2001). In particular, postero-laterally angled gaps between the cheek teeth were suggested as possible filtering channels. In describing the holotype of *Coronodon havensteini* (CCNHM 108), Geisler et al. (2017) noted that a Charleston Museum specimen ChM PV4745 from the same geologic unit as the holotype of *Coronodon* “…is clearly a juvenile specimen, and may be conspecific with *Coronodon havensteini*.” This individual has been included in many phylogenetic studies (Figs. 3a, c, 5a-b, 7b, 8b), but Geisler et al. (2017) did not incorporate this specimen in their trees because it is subadult.

Parsimony analysis of Geisler et al.’s (2017) morphological data groups aetiocetids (paraphyletic) with Mammalodontidae, and this combined clade is sister to Chaeomysticeti, which implies homoplasy in feeding characters (Fig. 7b). *Llanocetus* branches off at the next more basal node and is sister to the Chaeomysticeti + “Aetiocetidae” + Mammalodontidae clade. In the strict consensus tree, the evolution of lateral palatal foramina and sulci optimizes to a relatively basal internode, with homology of this trait inferred for *Llanocetus*, aetiocetids, and chaeomysticetes. The overall topology requires the secondary loss of lateral palatal foramina and sulci in mammalodontids and the reversal of mandible shape from ‘straight’ to ‘laterally concave’ in *Janjucetus*. *Coronodon* clusters with ChM PV5720 near the base of the mysticete tree in a trichotomy with *Metasqualodon* and a clade that includes all remaining mysticetes (Fig. 7b).

In comparison to our equally weighted parsimony analysis of morphology, the combined data tree with characters weighted by fit with concavity function k = 3 (figure 3 in Geisler et al. 2017) positions the late Eocene *Llanocetus* in a more crownward position as the sister group to Chaeomysticeti. This relationship, in combination with an early divergence of *Coronodon*, suggested a teeth-to-baleen scenario in which filtering with slots between the teeth evolved early, subsequent increases in the spacing of the cheek teeth with the development of baleen improved filter feeding, and this was followed by the complete evolutionary loss of postnatal teeth in modern baleen-bearing whales. Their hypothesis outlines an evolutionary feedback loop in which increased oral cavity dimensions, less dependence on teeth, and improved filtering efficiency drove increases in both body size and the oral cavity that then enabled even larger body size, as seen in truly giant modern mysticetes with elaborate baleen racks (Geisler et al. 2017).

The sequence of evolutionary transitions outlined in Geisler et al. (2017) makes some sense for their preferred tree based on Goloboff weighting with k = 3 (figure 4 in Geisler et al. [2017]), but as in all phylogenetic analyses of Mysticeti that include a dense sampling of extinct taxa with extensive missing data, this topology is highly unstable as indicated by shifting relationships with different concavity functions for parsimony-weighting of characters (i.e., k ranging from 2 to 9; see supporting online materials for Geisler et al. 2017) or equal weighting of characters (Fig. 7b). It is therefore a challenge to determine whether the unique dentition of *Coronodon* is autapomorphic for this taxon or perhaps represents the plesiomorphic condition for Mysticeti.

#### Peredo et al. (2018)

Peredo et al. (2018) (also see Peredo and Uhen 2016; Peredo and Pyenson 2018) expanded the morphological dataset of Boessenecker and Fordyce (2015a, b) by sampling additional extinct stem mysticetes (Fig 10c) and presented a novel, highly controversial interpretation of the teeth-to-baleen transition in Mysticeti. The authors described the Oligocene (33 Ma) chaeomysticete *Maiabalaena nesbittae* (Table 1) from Washington State (USA) and concluded that, “*Maiabalaena* had neither teeth nor baleen—it represents a surprising intermediate stage between modern filter-feeding whales and their toothed ancestors.” Furthermore, they noted that, “Phylogenetic analysis shows *Maiabalaena* as crownward of all toothed mysticetes, demonstrating that tooth loss preceded the evolution of baleen,” and that a suction feeding intermediate without teeth or baleen enabled the shift from feeding with teeth to batch filter feeding with just baleen as in extant mysticetes. If true, this scenario completely upends previous interpretations of the teeth-to-baleen transition in Mysticeti. Given the potentially profound evolutionary implications of this study, we scrutinize the evidence in detail below (also see Ekdale and Deméré 2022).

**Fig. 9.**
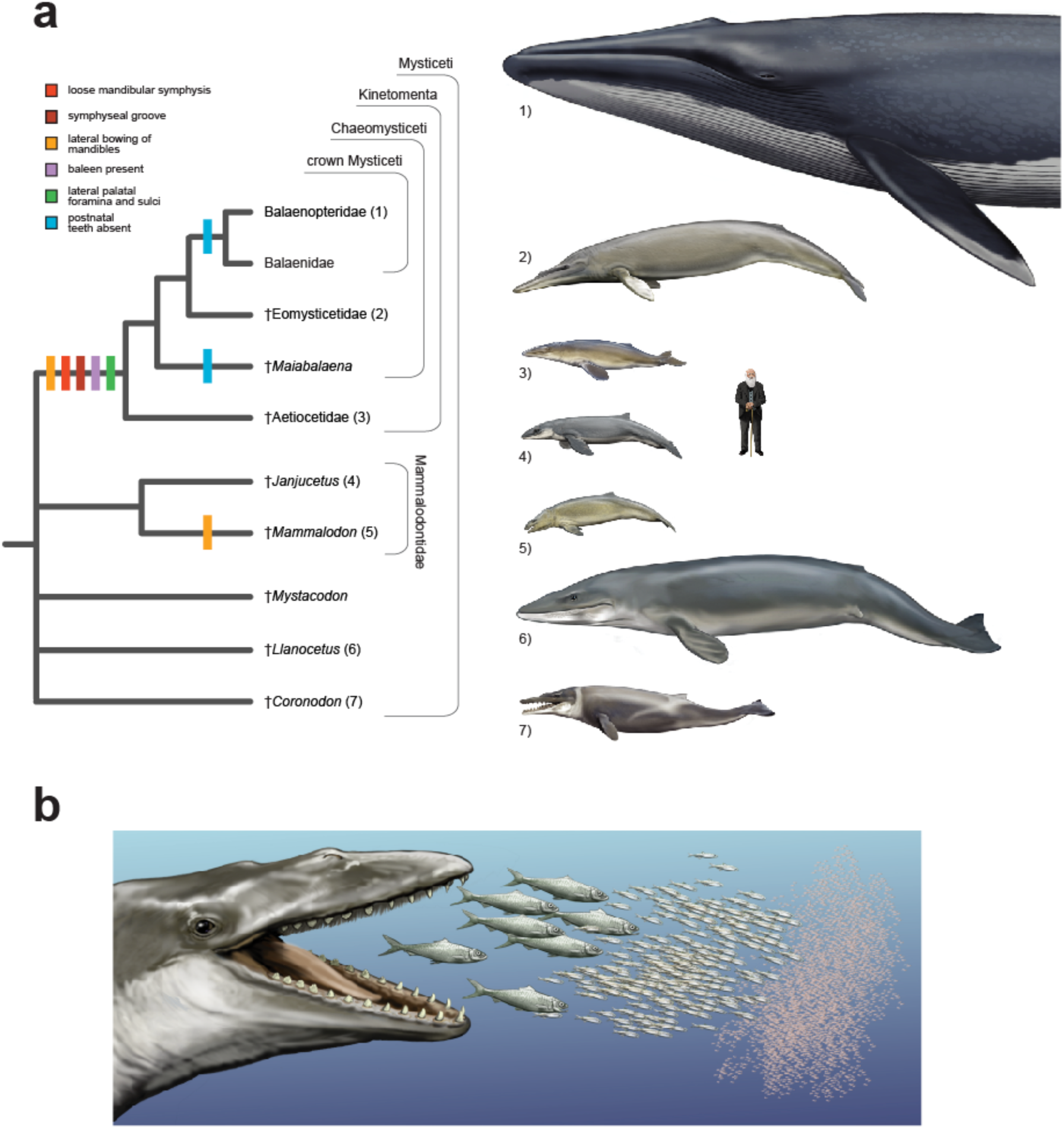
Tree that summarizes our interpretation of phylogenetic work on Mysticeti over the past 15 years (**a**) and one possible reconstruction of *Aetiocetus weltoni* (**b**) given this phylogenetic hypothesis. Early (Fig. 3a-c) and more recent studies (Figs. 5a, 7c, 8a-b) support Kinetomenta, the clade that includes Aetiocetidae and Chaeomysticeti. Kinetomenta is supported by a loose mandibular symphysis, a symphyseal groove, lateral palatal foramina, and perhaps baleen. Postnatal teeth were lost convergently in crown mysticetes and in *Maiabalaena* or were lost in the last common ancestor of Chaeomysticeti, followed by re-evolution of vestigial teeth in Eomysticetidae (here mapped as convergent loss). Baleen is reconstructed in the summary tree as initially evolving in the common ancestor of all kinetomentans with the evolution of lateral palatal foramina and a loose mandibular symphysis, but this interpretation remains controversial. Phylogenetic relationships among early-diverging toothed forms conflict in published studies, which hinders reconstruction of the earliest transitions related to feeding anatomy in Mysticeti. Human is shown to gauge approximate body size of seven representative mysticetes: 1) *Balaenoptera musculus*, 2) *Eomysticetus whitmorei*, 3) *Aetiocetus weltoni*, 4) *Janjucetus hunderi*, 5) *Mammalodon colliveri*, 6) *Llanocetus denticrenatus*, 7) *Coronodon havensteini*. In (**b**), *A. weltoni* is reconstructed as having a composite feeding apparatus with functional teeth and baleen that may have enabled feeding at a variety of trophic levels. Illustrations are by C. Buell.

**Fig. 10.**
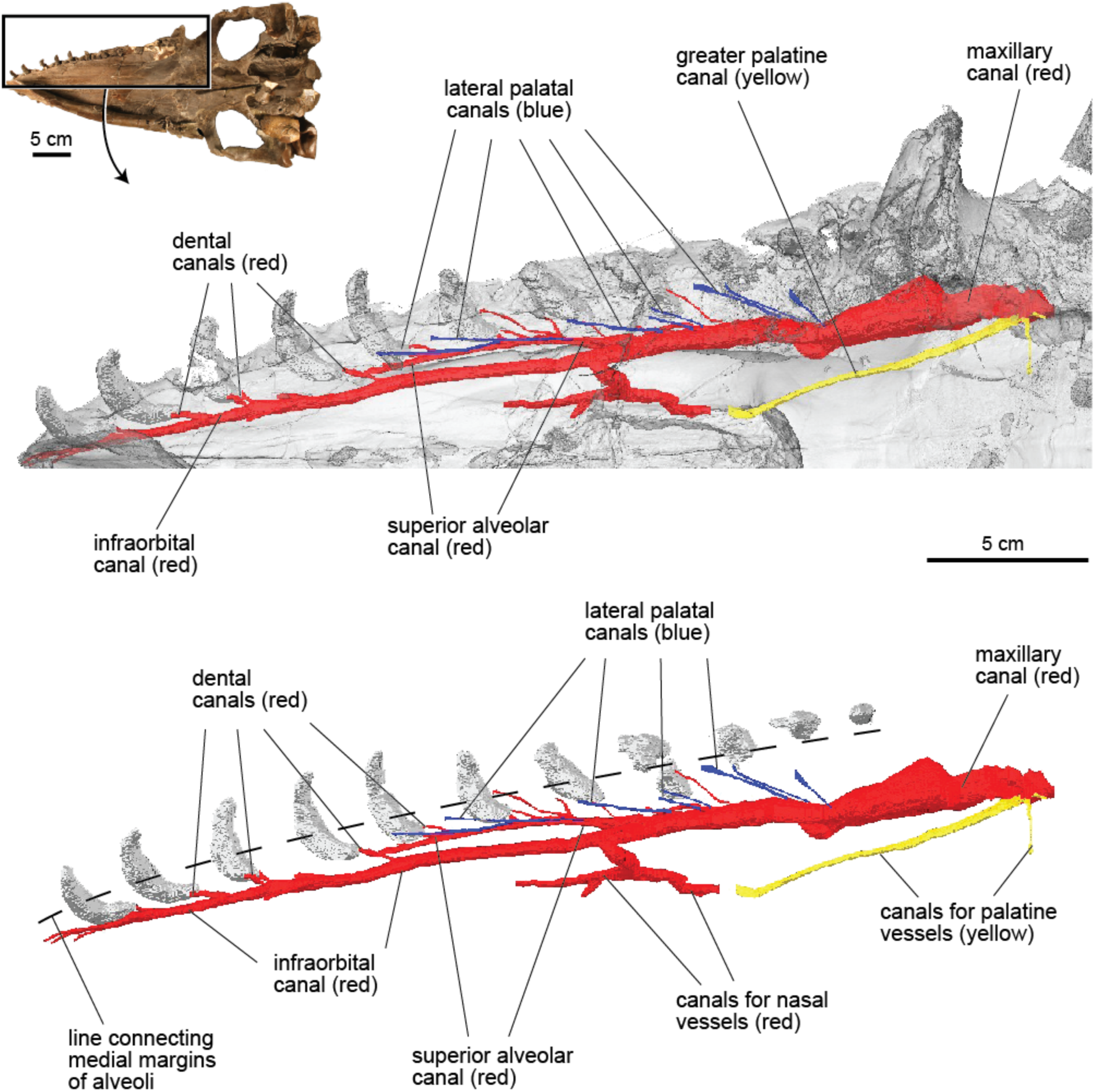
Digital models of rostrum and rostral canals of *Aetiocetus weltoni* (UCMP 122900) based on CT images. The ventral view of the skull (top left) shows the region of the skull that is illustrated below. The palate is rendered semi-transparent in the middle image, and bone is removed to reveal internal rostral canals in the bottom image. Image processing and digital segmentation of rostral canals is as described by Ekdale and Deméré (2021). Note that both the lateral palatal canals (blue) and the small dental canals from the alveoli connect with the superior alveolar canal (for the superior alveolar artery). This suggests a common source of vessels that deliver blood to teeth as well as vessels that deliver blood to baleen via the lateral palatal foramina. Also, see Figure 4a for a close-up photo of lateral palatal foramina and sulci in this specimen.

At first glance, the holotype skull of *Maiabalaena* (USNM 314627) appears to be an ordinary edentulous baleen-bearing mysticete, albeit an early one, with character states reminiscent of an eomysticetid-grade cetacean. The authors noted that, “The right maxilla preserves at least four, and as many as eight palatal foramina as identified by Deméré and Berta (2008) in various aetiocetids. These foramina lie between 23 and 30 cm from the anterior tip of the rostrum. All are shallow, laterally narrow foramen [*sic*] with anteroposteriorly elongated sulci that are angled between 20 and 45 degrees from the sagittal plane.” Given the detailed similarity of these features to lateral palatal foramina and sulci in other chaeomysticetes, the authors coded this critical character as ‘present’ in *Maiabalaena* (Fig. 7c).

A CTscan of the fossil skull, however, did not reveal connections between the palatal structures and the superior alveolar canal as seen in modern baleen whales (Sawamura 2008a, b; Ekdale et al. 2017) and in Oligocene aetiocetids (Peredo et al. 2018; Ekdale and Deméré 2022). In *Maiabalaena*, the shallow foramina at the edges of the palate penetrate only ∼5mm into the maxilla and then appear to terminate, but the authors noted that, “Our observations may be limited by CT resolution, or it may be attributed to the loss of alveolar bone and subsequent remodeling of the palatal margin”. Based on this CTscan with poor resolution, Peredo et al. (2018) concluded that *Maiabalaena* definitively lacked baleen because, “there is insufficient evidence for inferring baleen in stem mysticetes based solely on the absence of teeth…” and that, “…there is no evidence for using palatal foramina to exclusively infer the presence of baleen”, because several fossil cetaceans outside of Mysticeti have foramina on their palates similar to those in aetiocetids. Importantly, the only potentially compelling evidence presented *<u>for</u>* absence of baleen in *Maiabalaena* is the apparent lack of connection between the lateral palatal foramina and the superior alveolar canal. Peredo et al. (2018) essentially argued that the unique internal anatomy of the lateral palatal foramina and sulci in *Maiabalaena* indicated that baleen was unequivocally absent in this taxon but did not explain why the ‘dead-end’ lateral palatal foramina in this specimen do not connect to the superior alveolar canal or any other major canals in the rostrum. This condition has never been recorded previously or since in any mysticete.

Peredo et al. (2018) coded crown mysticetes, *Maiabalaena*, four aetiocetids, and five eomysticetids as ‘present’ for lateral palatal foramina and sulci but coded two extinct outgroup taxa, the odontocete *Simocetus rayi* and the basilosaurid *Zygorhiza kochii*, as ‘absent’ in their data matrix (Fig. 7c). In their text, however, Peredo et al. (2018) asserted that the presence of lateral palatal foramina in *Simocetus* and *Zygorhiza* demonstrates that these structures extend outside Mysticeti and nullifies the utility of these features for inferring baleen in extinct species. Given that the first test of anatomical homology is similarity (Patterson 1982; de Pinna 1991), it is apparent that for Peredo et al. (2018), the structure, relative position, and detailed similarity of palatal foramina in *Simocetus* and *Zygorhiza* were not compelling enough to hypothesize homology to the lateral palatal foramina of extant mysticetes. By contrast, the foramina on the palates of *Maiabalaena*, eomysticetids, and aetiocetids were considered similar enough to score as potential homologs to the lateral palatal foramina of extant mysticetes, which is indicated by the character codings in their data matrix (Fig. 7c).

Initial conjectures of anatomical homology are confirmed or rejected by assessing congruence with other characters via cladistic analysis (Patterson 1982; de Pinna 1991; Brower and Schawaroch 1996). The strict consensus tree based on Peredo et al.’s (2018) morphological dataset (Fig. 7c) positions *Maiabalaena* sister to *Sitsqwayk cornishorum*, another Oligocene chaeomysticete from Washington State. In an earlier paper, Peredo and Uhen (2016) reported that in the fragmentary skull of *Sitsqwayk*, “grooves along the palate suggest the presence of baleen” and interpreted the species as “… a toothless, baleen-bearing mysticete”, but they coded lateral palatal foramina and sulci as ‘uncertain’ (’?’) in this species. At any rate, *Sitsqwayk*, the purported sister species to *Maiabalaena,* was previously interpreted as an early edentulous baleen-bearing whale with lateral palatal sulci (Peredo and Uhen 2016). In minimum length trees, *Maiabalaena* + *Sitsqwayk* is sister to a clade including eomysticetids and all other chaeomysticetes sampled, which reinforces the view that *Maiabalaena* is a relatively ‘standard’ toothless, baleen-bearing mysticete, just like *Sitsqwayk* (Peredo and Uhen 2016). Indeed, for the feeding characters that were mapped, character states in *Maiabalaena* are identical to those of extant mysticetes (Fig. 7c), thus providing few insights regarding the teeth-to-baleen transition for these traits. Convinced that *Maiabalaena* lacked baleen, however, Peredo et al. (2018) concluded, “Given that *Maiabalaena* forms a clade with *Sitsqwayk*, we tentatively infer *Sitsqwayk* as lacking both teeth and baleen, as well”, a remarkable 180-degree reversal relative to the authors’ previous view just two years earlier in which *Sitsqwayk* was interpreted as a “…a toothless, baleen-bearing mysticete” (Peredo and Uhen 2016).

After reevaluating the published data, we find no compelling evidence for the absence of baleen in *Maiabalaena* or *Sitsqwayk* and instead interpret these taxa as baleen-bearing, toothless (or functionally toothless) mysticetes that filter fed. We predict that future higher-resolution CTscans of *Maiabalaena* and *Sitsqwayk* specimens (that are perhaps better preserved) will reveal connections between lateral palatal foramina and the superior alveolar canal as found previously in members of crown Mysticeti and Aetiocetidae (Ekdale et al. 2017; Peredo et al. 2018; Ekdale and Deméré 2022). It also will be of interest to determine whether CTscans of *Simocetus rayi* and *Zygorhiza kochii* record any detailed similarity to the lateral palatal foramina and sulci of mysticetes (i.e., connections to the superior alveolar canal).

As in several prior phylogenetic analyses of Mysticeti (Figs. 3a-c, 5a), Peredo et al.’s (2018) dataset supports Aetiocetidae as the sister group to Chaeomysticeti, with Mammalodontidae and Charleston Museum stem toothed mysticetes (ChM PV4745, ChM PV5720) branching from more basal nodes in the tree. The strict consensus tree supports a sequential evolution of features related to filter feeding (Fig. 7c). Incipient bowing of the mandibles appeared first, then a cluster of potential ‘filter-feeding’ characters evolved in the common ancestor of Aetiocetidae + Chaeomysticeti. Parsimony optimization reconstructs a single evolution of lateral palatal foramina at this node, with inheritance of lateral palatal foramina and sulci in all descendants for which this character can be coded, including aetiocetids, eomysticetids, *Maiabalaena*, and crown mysticetes (Fig. 7c). Likewise, a loose mandibular symphysis and a distinct symphyseal groove evolved at this critical point in the tree, prior to loss of the dentition. The overall mix of primitive and derived states at this ancestral node is exemplified in the mosaic feeding anatomy of aetiocetids (Fig. 4).

The presence versus absence of teeth does not map cleanly to the strict consensus tree (Fig. 7c). Following Boessenecker and Fordyce (2015b; Fig. 5a), Peredo et al. (2018) scored the eomysticetid *Tokarahia lophocephalus* as having a vestigial adult dentition (‘teeth: present in adult’). Like Boessenecker and Fordyce (2015a, b), Peredo et al. (2018) coded most eomysticetids as ‘uncertain’ (‘?’) for presence versus absence of teeth in adults, but definitively coded *Maiabalaena* and the highly fragmentary *Sitsqwayk* as ‘teeth: absent in adult’. It clearly is not possible to determine whether *Sitsqwayk* had upper teeth based on the holotype, and the margins of the palate are not complete in the holotype of *Maiabalaena* (Peredo and Uhen 2016; Peredo et al. 2018), which hinders definitive scoring of dental features in this species as well (Ekdale and Deméré 2022). The questionable character coding choices imply convergent losses of teeth in *Sitsqwayk + Maiabalaena* and in crown mysticetes (Fig. 7c), or alternatively the loss of postnatal teeth in the common ancestor of chaeomysticetes with the re-evolution of small teeth in *Tokarahia lophocephalus*, an equally parsimonious optimization.

Assuming that it is more probable to lose a complex anatomical feature two times in evolutionary history than to lose the structure and then re-evolve it anew (Gould 1970), the first parsimony optimization (Fig. 7c) is more credible and reconstructs the common ancestor of *Maiabalaena* and other chaeomysticetes as retaining at least vestigial postnatal teeth. In this scenario, the toothless state of modern baleen whales (Figs. 1e, 2) did not evolve until after the divergence between Eomysticetidae and crown Mysticeti. The hypothetical common ancestor of Chaeomysticeti is reconstructed as a species with postnatal teeth as well as functional lateral palatal foramina and sulci (Deméré and Berta 2008; Deméré et al. 2008) with connections to the superior alveolar canal, as seen in aetiocetids (Peredo et al. 2018; Ekdale and Deméré 2022). In this character mapping, *Maiabalaena* is just an evolutionary side-branch that independently lost its dentition with autapomorphic loss of connections between its lateral palatal foramina and the superior alveolar canal – assuming these traits are accurately coded (Fig. 7c). Alternatively, if the second parsimony optimization is accurate, the loss of postnatal teeth occurred prior to the split of *Maiabalaena* from other chaeomysticetes but after the evolution of lateral palatal foramina and sulci. In this second reconstruction, functional lateral palatal foramina are again reconstructed in the common ancestor of Aetiocetidae + Chaeomysticeti, and *Maiabalaena* is again interpreted as an aberrant side-branch in which the ‘plumbing’ for the lateral palatal foramina and sulci was lost (or perhaps was distorted by post-mortem processes). In summary, the interpretation of *Maiabalaena* as a representation of the hypothetical toothless, baleenless, suction-feeding ancestor that enabled the shift to batch filter feeding with baleen (Peredo et al. 2018) falls apart following scrutiny of the authors’ character codings, trees derived from these observations, parsimony optimization of character states, and reassessments of homology based on the overall cladistic analysis (Fig. 7c).

#### Fordyce and Marx (2018)

Fordyce and Marx (2018) presented a description of the holotype partial skeleton (USNM 183022) of *Llanocetus denticrenatus*, a large-bodied toothed mysticete from the Eocene of Antarctica (Table 1) that had been included in several previous systematic studies (Figs. 3c, 5b, 7a-b). They executed the most comprehensive phylogenetic study of Mysticeti to date that featured an expanded version of the Marx and Fordyce (2015) morphological dataset and Bayesian combined analysis with molecular data compiled by McGowen et al. (2009). *Llanocetus* presents a unique combination of anatomical features including extremely large body size for a stem toothed mysticete (∼8 meters), widely spaced teeth, and scattered bundles of fine sulci that surround alveoli for some of the upper premolars.

As in most previous (Figs. 5b, 7a-b) and subsequent (Fig. 8b) studies, *Llanocetus* was scored as ‘present’ for lateral palatal foramina and sulci. This coding is contentious, because no lateral palatal foramina have actually been observed in the holotype skull (Ekdale and Deméré 2022). Fordyce and Marx (2018) noted that, “Clusters of sulci are associated with a specific tooth, forming four distinct bundles of subparallel sulci each bound for a particular alveolus (figure 2). Medially, all of the bundles converge posteriorly into a parasagittally oriented area of fractured bone. This area is marked by longitudinal striations, interpreted as crushed and ventrally eroded vascular canals and sulci; however, there are no unequivocal foramina.” The condition in *Llanocetus* does not match the anatomy of lateral palatal foramina and sulci in aetiocetids in any specific way, differing in both shape and relative position (Deméré and Berta 2008; Peredo et al. 2018; Ekdale and Deméré 2022). To our knowledge, there also is no detailed similarity or compelling positional homology to lateral palatal foramina and sulci of any living or extinct chaeomysticete (e.g., Ekdale et al. 2015; Deméré et al. 2008). Fordyce and Marx (2018) described the holotype as a “virtually complete skull”, but several critical feeding characters (lateral palatal foramina, loose mandibular symphysis, symphyseal groove, lateral bowing of mandibles) currently cannot be coded unequivocally from the available fossil material (Fig. 8a).

Fordyce and Marx (2018) interpreted *Llanocetus* within the context of three previously proposed hypotheses for the teeth-to-baleen transition and concluded that *Llanocetus* used suction and/or suction-assisted raptorial feeding modes. They suggested that extremely widely spaced cheek teeth preclude the possibility of *Llanocetus* batch filtering small prey with its dentition as proposed for *Coronodon* (Geisler et al. 2017). Despite the fragmentary lower jaw and a fractured, incompletely preserved palate, the authors dismissed filter feeding using baleen or a convergently evolved keratinous filter. The fine sulci close to the upper teeth suggested to them that, if present, baleen would have to be positioned between the upper cheek teeth and thus interfere with the tight occlusion of *Llanocetus*’ upper and lower teeth (see Geisler et al. 2017; de Muizon et al. 2019 for alternative interpretations). Fordyce and Marx (2018) instead hypothesized that *Llanocetus* was a suction feeder that had “well-developed gums” that were nourished by blood vessels coursing through the palatal sulci and noted that several other early diverging toothed mysticetes have been interpreted as suction feeders with enlarged palatal gingivae (Fitzgerald 2010; Marx et al. 2015, 2016; Lambert et al. 2017).

Parsimony analysis of Fordyce and Marx’s (2018) morphology dataset surprisingly supports a strict consensus tree in which Mysticeti, as usually construed, is not monophyletic. Instead, a clade that includes the earliest known mysticetes (*Llanocetus*, *Mystacodon*) is sister to a much larger clade that includes Odontoceti and all remaining mysticetes (Appendix 1). By contrast, the combined data Bayesian consensus of Fordyce and Marx (2018) supports monophyly of Mysticeti (posterior probability = 0.95), so we mapped feeding characters onto this topology (Fig. 8a). In direct contrast to previous work by these authors (Figs. 3d, 5b; Marx 2011; Marx and Fordyce 2015), Aetiocetidae and Mammalodontidae do not form a clade, and aetiocetids (paraphyletic) instead are the closest relatives of Chaeomysticeti. Mammalodontidae (*Janjucetus* + *Mammalodon*) is the next most basal branch on the mysticete stem lineage, and the general pattern of relationships is replicated in the strict consensus parsimony tree based on just morphological characters (Appendix 1). Divergences among the earliest lineages of Mysticeti are more muddled, however, due to nodes with extremely low Bayesian posterior probabilities (Fig. 8a) that conflict with the parsimony analysis of morphology (Appendix 1). This instability at the base of the tree hinders evolutionary reconstructions, but even if the weakly supported nodes in the Bayesian consensus are collapsed, the tree still implies convergence between the incompletely preserved sulci around the upper dentition of *Llanocetus* and the lateral palatal foramina and sulci observed in aetiocetids and chaeomysticetes (Fig. 8a). Optimization of suction feeding on the tree is ambiguous given the mix of putative suction feeders (*Mammalodon*, *Llanocetus*, *Mystacodon*), a possible dental filterer (*Coronodon*), and a macrophagous predator (*Janjucetus*) among the earliest-diverging clades of Mysticeti.

#### de Muizon et al. (2019)

de Muizon et al. (2019) provided an expanded description of the oldest known mysticete, *Mystacodon selenensis* (Fig. 1b; Table 1), and extended earlier phylogenetic work by Lambert et al. (2017). Parsimony analysis yielded a strict consensus tree (Fig. 8b) that implies a much simpler pattern of evolution for feeding characters relative to their initial study (Fig. 7a). In the new tree, the five feeding characters generally map in a sequential, stepwise pattern on the mysticete stem lineage as in several previous studies (Fig. 3a-c, 5a, 7c), with only incipient lateral bowing of the mandibles evolving convergently multiple times. Aetiocetidae is sister to Chaeomysticeti, with several lineages (*Llanocetus*, a clade of *Coronodon* and other Charleston Museum toothed mysticetes, Mammalodontidae, and *Mystacodon*) branching from progressively more basal positions on the stem lineage of Mysticeti. As in Lambert et al. (2017) (Fig. 7a), *Mystacodon* from the late Eocene is the sister group to all remaining mysticetes in the tree. The authors inferred a central importance for suction feeding early in mysticete evolution based on the phylogenetic positions of *Mystacodon* and *Mammalodon* that have been interpreted as benthic suction feeders (Fitzgerald 2010; Lambert et al. 2017; de Muizon et al. 2019). *Llanocetus*, the second Eocene toothed mysticete in de Muizon et al.’s (2019) tree, did not group with *Mystacodon* as in Fordyce and Marx’s (2018) phylogenetic study (Fig. 8a; Appendix 1). Instead, the presence of lateral palatal foramina and sulci supports a sister group relationship between *Llanocetus* and Aetiocetidae + Chaeomysticeti (Fig. 8b). The authors concluded that, “…the position of *Llanocetus* as a sister group of the clade Aetiocetidae and Chaeomysticeti renders possible a single acquisition of palatal keratinous appendages in the clade (*Llanocetus*, (Aetiocetidae, Chaeomysticeti)).”

### Summary of phylogenetic results

Clearly, research on mysticete phylogeny over the past 15 years has yielded a broad spectrum of results in parsimony analyses of morphological characters (Figs. 3, 5, 7, 8; Appendix 1). However, some important common patterns have emerged (Fig. 9a). Early studies (Fitzgerald 2006, 2010; Deméré et al. 2008) supported aetiocetids (Figs. 1d, 4) as the closest relatives of Chaeomysticeti (Fig. 3a-c). The most recent and largest phylogenetic analyses of Mysticeti to date (Fordyce and Marx 2018; Peredo et al. 2018; de Muizon et al. 2019) corroborate the close relationship between aetiocetids and Chaeomysticeti as well as the generally stepwise, sequential evolution of feeding characters on the stem lineage of crown Mysticeti (Figs. 7c, 8a-b). Importantly, in two of these recent studies (Fordyce and Marx 2018; de Muizon et al. 2019), the grouping of aetiocetids with Chaeomysticeti contradicts earlier, less comprehensive phylogenetic work by the same authors (Marx 2011; Marx and Fordyce 2015; Lambert et al. 2017) that instead grouped aetiocetids with Mammalodontidae in a clade of stem toothed mysticetes. The early trees (Fig. 3c, 8b, 7a) were all based on variations of the matrix compiled by Marx (2011) that provided the initial support for the Aetiocetidae + Mammalodontidae clade. This grouping implies wholesale homoplasy in feeding characters that are found in all extant mysticetes (Figs. 5b, 7a; also see Geisler et al. [2017], Fig. 7b). A suite of characters related to the supraorbital process of the frontal and the orbital rim provide the majority of character support for this controversial ‘large-eyed’ clade, and the recycling of similar characters in multiple studies generated the same phylogenetic result. With better character and taxon sampling, however, the Aetiocetidae + Mammalodontidae clade disappeared (Figs. 8a-b), suggesting that correlated homoplasy driven by developmental or adaptive factors related to the enlarged eyes of aetiocetids and mammalodontids could explain resolution of this unnatural grouping in the earlier work.

Given that multiple research groups have converged on the same phylogenetic solution in which aetiocetids cluster with chaeomysticetes and that several key feeding characters provide support for this clade (Figs. 3a-c, 5a, 7c, 8a-b), it is useful at this time to provide a name for the mysticete clade composed of Aetiocetidae and Chaeomysticeti. We propose the clade name Kinetomenta (‘moveable chin’) for the last common ancestor of *Balaenoptera musculus* and *Aetiocetus weltoni* and all of its descendants (Fig. 9a). We employed *A. weltoni* as the representative aetiocetid in defining the name because the holotype cranium and mandible of this species are well preserved, relatively complete, and have been CT scanned (Deméré and Berta 2008; Ekdale and Deméré 2022). We note, however, that with this phylogenetic definition, the content of Kinetomenta varies for different cladistic hypotheses, depending on whether Aetiocetidae is resolved as monophyletic (Figs. 3b-c, 7c, 8b) or paraphyletic (Figs. 3a, 5a, 8a).

Another strong consensus in phylogenetic studies reviewed here is a close relationship between Eomysticetidae and crown-clade Mysticeti (Fig. 9a). This grouping is evidenced by a variety of character states, including more extreme lateral bowing of the mandibles, loss of a functional adult dentition, and prominent lateral palatal foramina and grooves (when preserved). As in aetiocetids, eomysticetid fossils preserve evidence for the simultaneous presence of both postnatal teeth and baleen, implying transitional phenotypes, but for both groups, the inference is still debated in the literature (e.g., Peredo et al. 2018; de Muizon et al. 2019). All recent trees (Figs. 3, 5, 7-8) suggest a large phenotypic gap between the branching point for Aetiocetidae and the node where Eomysticetidae split with crown group Mysticeti. On this internal branch, a large reduction in the dentition is inferred, along with an expansion of the palatal vasculature, perhaps signifying the increased reliance on filter feeding with baleen and decreased reliance on tooth-assisted predation. In the cladistic analysis of Peredo et al. (2018), an unnamed clade composed of *Sitsqwayk* + *Maiabalaena* branches in this sector of the mysticete tree (Figs. 7c, 9a), but these species already show a highly reduced dentition and do not close the large phenotypic gap between aetiocetids and eomysticetids.

Studies that have argued for the centrality of suction feeding in early stages of the transition to filter-feeding with baleen are common in the recent literature (e.g., Fitzgerald 2010; Marx et al. 2015, 2016; Hocking et al. 2017a; Lambert et al. 2017; Fordyce and Marx 2018; Peredo et al. 2017, 2018). Yet, the phylogenetic relationships among toothed mysticetes that have been interpreted as suction feeders, such as *Mystacodon*, *Mammalodon*, and *Llanocetus,* are highly unstable across different studies, with the possible dental filterer, *Coronodon*, adding to the disarray among the earliest branching events in Mysticeti (Figs. 3, 5, 7-8). In part, this confusion may be due to missing/ambiguous data for the feeding apparatus and inadequate primary homology assessment. For example, the known fossils of both *Llanocetus denticrenatus* and *Mammalodon colliveri* do not preserve key features of the mandibular symphysis that link aetiocetid toothed mysticetes to Chaeomysticeti, and the coding of mandibular curvature is uncertain in *Llanocetus denticrenatus* due to the fragmentary lower jaw of the holotype. As discussed earlier, the presence of lateral palatal foramina and sulci represents another likely synapomorphy for Kinetomenta. In some studies, *Llanocetus* has grouped close to Kinetomenta, suggesting homology of the palatal vasculature in these taxa (e.g., 11B), but more commonly, *Llanocetus* is positioned much deeper in the mysticete tree, consistent with its late Eocene age and with convergent evolution of its superficially similar palatal features (e.g., 11A).

### The evolution of baleen and filter feeding

Unequivocal assertions about the presence of baleen in individual mysticete taxa can only be made for specimens with preserved baleen. Obviously, all extant mysticetes can be coded for the presence of baleen, while to date only four extinct mysticete taxa with fossilized baleen can be scored unambiguously as ‘baleen-bearing’. These include *Piscobalaena nana* (Marx et al. 2017), *Miocaperea pulchra* (Bisconti 2012), *Balaenoptera siberi* (Pilleri 1989), and *Incakujira anillodefuego* (Marx and Kohno 2016). However, based on parsimony optimization, it is possible to add a number of extinct taxa to the list of baleen-bearing mysticetes. All extinct crown mysticetes are descended from the last common ancestor of the crown clade and are bracketed by extant balaenids and extant balaenopterids that express baleen. It is therefore simplest to assume that all fossil crown mysticetes possessed baleen, which they inherited from a common baleen-bearing progenitor. Inferences regarding the presence of baleen at more basal nodes in the tree have been guided by phylogenetic interpretations of stem mysticetes and the characters that they share with baleen-bearing crown mysticetes.

The presence of baleen in extinct mysticetes most likely extends stemward to all eomysticetids based on their highly reduced adult dentition (Fig. 6), the presence of lateral palatal foramina and sulci in specimens with well-preserved palates, the presence of a loose mandibular symphysis and symphyseal groove, as well as an elongated rostrum with a dorso-ventrally thin palatal margin that likely enabled skim filter feeding (Boessenecker & Fordyce 2015a, b; 2017). However, when it comes to extinct mysticetes that branch from more basal nodes on the stem lineage, statements about the presence of baleen become more controversial, while at the same time the demand for unequivocal evidence for baleen becomes more vehement. At the center of this controversy are the aetiocetids (Fig. 4) and whether or not members of this long extinct, tooth-bearing mysticete group possessed some form of baleen (Figs. 1c, 9b). The proposal that members of the chaeomysticete clade consisting of *Maiabalaena nesbittae* and *Sitsqwayk cornishorum* (Fig. 7c) lacked both teeth and baleen (Peredo et al. 2018) also is controversial (e.g., Peredo and Uhen 2016).

For aetiocetids, the argument for the presence of proto-baleen medial to the upper toothrow is primarily based on phylogenetic placement sister to Chaeomysticeti (Fig. 9a), the occurrence of lateral palatal foramina and sulci (Fig. 4a), and the proposed homology between these structures and the neomorphic neurovascular palatal anatomy associated with the baleen apparatus in crown mysticetes (Fig. 2b; Deméré and Berta 2008; Deméré et al. 2008). This homology is strongly supported by recent high resolution CT imaging of the holotype skull of *Aetiocetus weltoni* that found direct connections between branches of the superior alveolar canal and the lateral palatal foramina (Fig. 10; Ekdale and Deméré 2022), which matches the vascular supply pattern to baleen found in the extant gray whale, *Eschrichtius robustus* (Ekdale et al. 2015). Importantly, the CT imaging of *A. weltoni* also revealed retention of connections between the superior alveolar canal and dental canals of the upper dentition, which is consistent with the plesiomorphic pattern of blood supply to teeth in odontocete cetaceans and terrestrial mammals (Ekdale and Deméré 2022). This pattern suggests that the rostral dental vasculature of stem toothed mysticetes was co-opted to supply blood to baleen in the common ancestor of aetiocetids and chaeomysticetes (Kinetomenta). Peredo et al. (2017) noted that, “While recent work suggests the baleen in extant gray whales (*Eschrichtius robustus*) may be vascularized via branches of the superior alveolar artery (Ekdale et al. 2015), it remains unclear how these branches could supply both teeth and baleen, as would be necessary for extant mysticetes *in utero*, or for aetiocetids if these possessed both types of feeding structures.” The new CT imaging of *A. weltoni* clearly shows how the superior alveolar artery could supply both teeth and baleen by branching further to supply blood through the neomorphic lateral palatal canals (Fig. 10; Ekdale and Deméré 2022).

Several alternative interpretations of the palatal anatomy of stem toothed mysticetes have been proposed, including suggestions that the lateral palatal foramina and sulci are not universal in aetiocetids (Peredo et al. 2017, 2018), that these nutrient foramina evolved very early on the mysticete stem and then were lost in mammalodontids (Marx and Fordyce 2015), that the structures evolved convergently within a ‘big-eyed clade’ of toothed mysticetes (Lambert et al. 2017), in the early toothed mysticete *Llanocetus* (Fordyce and Marx 2018), or in non-mysticete cetaceans (Peredo et al. 2018), that the structures are not homologous with the baleen-related neurovascular anatomy of extant mysticetes (Peredo et al. 2017), or that the palatal vasculature of aetiocetids represent an adaptation for suction feeding with subsequent exaptation (co-option) for baleen-assisted filter feeding (Marx et al. 2016; Fordyce and Marx 2018; Peredo et al. 2018). As summarized in various phylogenetic studies, support for these competing hypotheses is mixed, as is evidence for lateral palatal foramina and sulci (and perhaps baleen) evolving only once (Figs. 3b-d, 5a, 7c, 8b), evolving once with loss in some taxa (Figs. 3a, c, 5b, 7b), or evolving convergently within Mysticeti (Figs. 7a, 8a).

Importantly, a majority of these studies coded lateral palatal foramina and sulci as present in all aetiocetids and all chaeomysticetes in which palatal anatomy is well preserved enough to permit coding of presence versus absence (Figs. 3b, 5a-b, 7a-c, 8a-b). Therefore, in trees that support aetiocetids as the closest relatives of Chaeomysticeti, it is parsimonious to infer that the lateral palatal foramina function as in extant mysticetes, providing innervation and nourishment to the continuously growing baleen. An exception is the proposal by Peredo et al. (2018) that *Maiabalaena* (USNM 314627), positioned apical to aetiocetids and basal relative to eomysticetids in their tree (Fig. 7c), has lateral palatal foramina and sulci (coded ‘present’) but these nutrient foramina seem to be ‘dead ends’ that do not connect to the superior alveolar canal. Given their character codes, the most parsimonious solution is a single origin of lateral palatal foramina and sulci (and perhaps the first appearance of proto-baleen) in the last common ancestor of Kinetomenta (Fig. 7c) with a later autapomorphic disruption of the connection between the lateral nutrient foramina and the superior alveolar canal in *Maiabalaena*, or perhaps in the common ancestor of *Maiabalaena* + *Sitsqwayk.* This reconstruction is wholly consistent with previous and subsequent phylogenetic studies that provide systematic evidence for a single origin of baleen-related foramina and sulci in Mysticeti (Fig. 9a), but also begs the question of why the connection to the superior alveolar canal was lost in this taxon. CTscans of additional fossils with key positions along the stem lineage of Mysticeti are required to fill gaps in our current understanding of palatal vasculature evolution in this clade.

Although perhaps tangential to the question of whether baleen was present in aetiocetids, eomysticetids, *Maiabalaena*, and *Sitsqwayk*, it is noteworthy that all of these taxa also possess an unsutured mandibular symphysis with a longitudinal symphyseal groove (Figs. 2a, 4b, 6a, c), apomorphic features shared by all kinetomentans that are adequately preserved (Deméré and Berta 2008; Fitzgerald 2012). Functionally, this anatomy permits enhanced mobility at the symphysis, which has been recognized as critical in accommodating the specialized bulk filter-feeding modes seen in extant mysticetes (Lambertsen et al. 1995; Pyenson et al. 2012). In aetiocetids, mobility at the mandibular symphysis could have permitted controlled manipulation of the fully-toothed mandibles in relation to a primitive keratinous filter medial to the upper tooth row. Viewed in a phylogenetic context, it therefore may be no coincidence that a kinetic mandibular symphysis (Fig. 4b) and evidence for neurovascular structures related to baleen (Figs. 4a, 10) are associated in aetiocetids, lending further support to the idea that these Oligocene toothed mysticetes possessed some form of proto-baleen (Figs. 1d, 9b).

From a functional standpoint, competing phylogenetic hypotheses (Figs. 3, 5, 7, 8) suggest alternative interpretations of the transition from raptorial feeding with teeth as in stem cetaceans (’archaeocetes’) to baleen-assisted filter feeding as in modern mysticetes. These include the following four scenarios that take into account recent evidence that eomysticetids likely retained a vestigial dentition at the tips of their jaws (Fig. 6: Boessenecker and Fordyce 2015a, b) and the suggestion that several early toothed mysticetes were likely suction feeders (Fitzgerald 2010; Lambert et al. 2017; Fordyce and Marx 2018; de Muizon et al. 2019):

1. <u>’Transitional chimaeric feeder’ scenario</u>: Raptorial feeding with teeth, to raptorial/suction feeding with teeth, to a multifarious feeding system with both raptorial/suction feeding with teeth and early filtering with primordial baleen, to filter feeding with baleen and retention of vestigial teeth, to complete loss of postnatal teeth and filter feeding with baleen (Fitzgerald 2006; Deméré et al. 2008).
2. <u>’Early dental filterer</u>’ scenario: Raptorial feeding with teeth, to raptorial/suction feeding on single prey and batch filtering of prey with teeth, to raptorial/suction feeding with teeth on single prey and batch filtering with primordial baleen, to filter feeding with baleen and retention of vestigial teeth, to complete loss of postnatal teeth and filter feeding with baleen (Geisler et al. 2017).
3. <u>’Transitional toothed suction feeder’</u> scenario: Raptorial feeding with teeth, to raptorial/suction feeding with teeth and enlarged gums, to filter feeding with baleen and retention of vestigial teeth, to complete loss of postnatal teeth and filter feeding with baleen (Fordyce and Marx 2018).
4. <u>’Transitional toothless suction feeder’</u> scenario: Raptorial feeding with teeth, to suction-assisted raptorial feeding with teeth, to complete loss of postnatal teeth and suction feeding (i.e., suction feeding without teeth or baleen), to filter feeding with just baleen (Peredo et al. 2018).

These scenarios can be evaluated based on their compatibility with hypotheses of homology implied by particular phylogenetic trees. Some scenarios provide simpler, more efficient explanations than others. The first (’transitional chimaeric feeder’) is broadly consistent with multiple phylogenetic studies, in particular those that position aetiocetids as the closest relatives of Chaeomysticeti (Figs. 3a-c, 5a, 7c, 8a-b, 9a). Depending on the specifics of character coding, these trees imply a generally stepwise evolution of feeding characters in which aetiocetids express the composite feeding anatomy of the inferred kinetomentan ancestor (Fitzgerald 2006; Deméré et al. 2008). Possessing a well-developed, functional adult dentition, aetiocetids also express multiple character states that are associated with baleen-assisted filter feeding in modern mysticetes (Figs. 4, 10). The ‘transitional chimaeric feeder’ scenario does not require extensive exaptation in these feeding characters, but instead posits that a loose mandibular symphysis, a symphyseal groove, and lateral palatal foramina and sulci were utilized by aetiocetids in filter-feeding with proto-baleen, similar to how these features are utilized by extant mysticetes for this function. This scenario therefore posits a straightforward transition from tooth-assisted predation on single larger prey to filter-feeding on batches of smaller prey items via a transitional mosaic anatomy that could accomplish both (Fig. 9b). A composite feeding apparatus that combines both primitive and derived feeding modes could expand the range of prey items and facilitate the evolutionary transition to sole reliance on baleen-assisted bulk filter feeding (Deméré et al. 2008). This ‘step-wise’ evolutionary hypothesis (Marx 2006; Deméré et al. 2008) does not preclude the possibility that ancestral mysticetes with both teeth and baleen utilized suction feeding in their repertoire of predatory behaviors.

The most controversial aspect of the ‘transitional chimaeric feeder’ hypothesis is whether primordial baleen could function adequately in a whale with a functional adult dentition. Some researchers have suggested that baleen medial to the upper tooth row would interfere with the dentition when grasping or manipulating large prey and that a well-developed dentition with tight occlusion would interfere with baleen-assisted filter feeding (Marx et al. 2016; Peredo et al. 2017, 2018; Fordyce and Marx 2018). Others have countered that any interference would depend on the shape and positioning of the baleen. Early baleen may have been organized as simple bundles of keratinous bristles rather than racks of much larger and structurally complex baleen plates (Deméré et al. 2008; de Muizon et al. 2019; Ekdale and Deméré 2022). Depending on how the jaws were positioned, a loose collection of fairly short baleen bristles that are medial to the toothrow could, in our opinion, function as a rudimentary sieve for capture of small prey. Even with the limited gaps between the upper and lower cheek teeth of some aetiocetids, filtering would be possible if the jaws were not completely closed and if medially positioned proto-baleen was the appropriate length and shape. Keep in mind that in extant mysticetes the baleen rack lies medial to the lower jaw when the mouth is closed and the upper and lower lips are occluded.

In the end, it is likely that only exquisitely preserved aetiocetid specimens with fossilized baleen would convince skeptics that the ‘transitional chimaeric feeder’ scenario is correct. We therefore suggest that whenever relatively complete toothed mysticete fossils are discovered in the future, the palate should be prepared with great care to document any evidence of keratinous structures in the rock matrix. Researchers generally have not asserted the presence of baleen in taxa with prominent teeth when lateral palatal foramina are absent (e.g., Fitzgerald et al. 2006, 2010; Geisler et al. 2017; Peredo et al. 2018; de Muizon et al. 2019). This is perhaps a conservative approach, but we see no reason to believe that the earliest baleen, which may have been just sparse patches of short keratinous bristles, required an extensive neomorphic palatal vasculature. Incipient baleen may have been derived initially in a toothed mysticete that lacked lateral palatal foramina, with enhanced palatal vasculature evolving only later as the baleen increased in size and extent.

The second scenario, ‘early dental filterer’, posits that a key initial step in the transition to filter feeding with baleen included mysticetes that batch filtered small prey by utilizing slots between their denticulate teeth (Geisler et al. 2017). Only one toothed stem mysticete has been seriously considered to be a dental filterer - *Coronodon havensteini* (Fig. 1a). The ‘early dental filterer’ hypotheses (Geisler et al. 2017) requires an initial shift to dental filtering, a subsequent shift to filtering with incipient baleen in toothed mysticetes (e.g., aetiocetids and perhaps *Llanocetus*), and a final transition to toothless (or nearly toothless) chaeomysticetes that filter with well-developed baleen. Like the ‘transitional chimaeric feeder’ scenario, the ‘early dental filterer’ scenario does not entail extensive exaptational shifts in function for the five feeding characters examined here (Fig. 2) but does require the evolution of a novel feeding mode not observed in any extant cetaceans - batch filter feeding with the dentition.

Some of the phylogenetic trees presented in Geisler et al. (2017) fit this pattern, but the authors noted that the evolution of dental spacing on their preferred tree was “complicated”. *Coronodon* has been included in two subsequent phylogenetic analyses - Fordyce and Marx (2018) and de Muizon et al. (2019). The former positions *C. havensteini* as a basal branch in the mysticete tree, which is consistent with the hypothesis of very early evolution of dental filtering in the clade, but progressively apical branches in the tree include a mix of putative suction-feeders and the macrophagous raptorial predator, *Janjucetus*. This topology does not reconstruct a simple transformational sequence from filtering with teeth, to filtering with baleen and prominent teeth, to filtering solely with baleen (Fig. 8a). The tree based on morphological data from de Muizon et al. (2019) is a better fit to this scenario, because in this case, *Coronodon* is more closely related to chaeomysticetes, aetiocetids, and *Llanocetus*, all of which have been interpreted as baleen-bearing mysticetes by some authors (Fig. 8b). In addition to currently ambiguous phylogenetic support for the ‘early dental filtering’ scenario, the interpretation of *Coronodon* as a tooth-assisted filterer has been challenged based on a poor morphometric match between the teeth of *Coronodon* and those of distantly related filter-feeding seals (Hocking et al. 2017b), but this critique hinges on the assumption that dental filtration can be achieved in just one way in all marine mammals.

The third scenario, ‘transitional toothed suction feeder’, stresses the central importance of suction feeding in the teeth-to-baleen transition and has gained support from studies of early stem toothed mysticetes that have been interpreted as suction feeders (Fitzgerald 2010; Marx et al. 2016; Lambert et al. 2017; Fordyce and Marx 2018; de Muizon et al. 2019). In this scenario, aetiocetids are seen as suction feeders or suction-assisted raptorial predators that did not have baleen (Marx et al. 2016; Fordyce and Marx 2018). Instead of being associated with baleen and filter-feeding, lateral palatal foramina and sulci in aetiocetids are interpreted as evidence for increased blood flow to enlarged gums that facilitated suction-assisted feeding. A mobile mandibular symphysis and symphyseal groove (Fig. 4b) also are reinterpreted as characters that evolved prior to the evolution of filter feeding in the ‘transitional toothed suction feeder’ scenario.

A primary impetus for this hypothesis is the opinion that a primitive baleen filter in aetiocetids is not biomechanically possible, because baleen medial to the upper toothrow would surely hinder a composite feeding system (Marx et al. 2015, 2016; Fordyce and Marx 2018). As noted above, this opinion is contentious and depends on the position of baleen on the palate, the morphology of early baleen, how far apart the upper and lower jaws were positioned during filter-feeding bouts, as well as the shape of the gums, tongue, and lips. All of these factors are basically unknown (and perhaps unknowable) without future fossil finds that preserve soft tissue anatomy. Therefore at present, any biomechanical modeling that attempts to interpret the complex interactions of these various anatomical unknowns would be highly speculative.

Phylogenetic hypotheses that fit the ‘transitional chimaeric feeder’ scenario (Figs. 3a-c, 5a, 7c, 8a-b) also are generally compatible with the ‘transitional toothed suction feeder’ scenario. In these trees, various purported suction feeding mysticetes branch from basal lineages (albeit with certain taxa that may not be suction feeders) and aetiocetids group close to Chaeomysticeti. If aetiocetids are interpreted as suction feeders with enlarged gums instead of baleen, then the critical importance of suction-feeding toothed taxa at the teeth-to-baleen transition is credible. Relative to the ‘transitional chimaeric feeder’ hypothesis, however, the ‘transitional toothed suction feeder’ hypothesis requires additional changes in function/behavior for various traits (exaptations), as well as changes that account for the gain and loss of unique morphologies in extinct taxa (e.g., enlarged gums utilized in suction feeding). These additional evolutionary derivations imply a more convoluted, less parsimonious pattern of evolution.

An example of possible exaptation in the feeding apparatus of mysticetes was outlined by Fordyce and Marx (2018). They rejected the early evolution of proto-baleen in stem toothed mysticetes and instead hypothesized that aetiocetids and *Llanocetus* (taxa with evidence for lateral palatal vasculature) had “well-developed gums”. They noted that, “Because palatal sulci directly supply the gingiva, they are best interpreted as an osteological correlate of enlarged gums, such as those that give rise to baleen in modern whales. This situation is analogous to the evolution of flight feathers in birds: although they are undoubtedly correlated with moving in air, they originally evolved for a different purpose (Brusatte et al. 2015). Well-developed gingivae have been inferred for virtually all archaic mysticetes (Fitzgerald 2010; Marx et al. 2015, 2016; Geisler et al. 2017).” Marx et al. (2016) made the same avian analogy in arguing that lateral palatal foramina and sulci initially functioned to supply blood to enlarged gums and were only later co-opted to also supply blood to ever-growing baleen, with large gums still present in extant mysticete species.

There are some problems with this line of reasoning. First, it should be noted that “well-developed gingivae” have not been inferred in “virtually all archaic mysticetes”, and unlike baleen, fossilized gums have not been preserved in any extinct mysticete. Instead, such soft anatomical structures have been hypothesized in a subset of early mysticetes (e.g., *Mammalodon*, *Mystacodon*, *Coronodon*), but not others (e.g., Fitzgerald 2006; Deméré et al. 2008; Solis-Añorve et al. 2019). Second, Fordyce and Marx (2018) suggested that the palatal sulci of aetiocetids and *Llanocetus* provided increased blood supply to enlarged gums, but other early mysticete taxa with putatively large gums lack lateral palatal foramina and sulci (Fitzgerald 2010; Marx et al. 2016; Geisler et al. 2017; Lambert et al. 2017; de Muizon et al. 2018), making this anatomical feature an especially poor “osteological correlate of enlarged gums”. Third, we are not aware of any phylogenetically-controlled allometric studies that have quantified the relative thickness of gingivae in extant odontocetes versus extant mysticetes. So, the “enlarged gums, such as those that give rise to baleen in modern whales” (Fordyce and Marx 2018) may not be especially large at all in extant large-bodied mysticetes. The prominent gingivae hypothesized in early suction feeding mysticetes would then represent a unique morphology, not the ancestral state inherited by living mysticetes. This is especially true if the derived gums of suction-feeding toothed mysticetes resembled the complex ‘gum teeth’ of the extant Dall’s porpoise (*Phocoenoides dalli*), which has been speculated in the literature (e.g., Miller 1929; Marx et al. 2016).

The early evolution and then loss of enlarged, modified gums would imply additional evolutionary steps, as would the proposed change of function for the lateral palatal vasculature from nourishment of large gums in aetiocetids to nourishment of constantly growing baleen in modern baleen whales. The same parsimony argument applies to exaptation of the loose mandibular symphysis. In the ‘transitional toothed suction feeder’ scenario, this derived feature did not evolve to assist filter-feeding with baleen, and surely must have evolved for some other function in suction-feeding aetiocetids that lacked baleen. This is of course possible, but such compounding of exaptational shifts should be minimized when reconstructing evolutionary patterns within a parsimony framework.

The fourth scenario, ‘transitional toothless suction feeder’, is similar to the third proposal but is more extreme. In this hypothesis, suction feeding without baleen also is inferred for stem mysticetes that are toothless or retain small postnatal teeth. Peredo et al. (2018) suggested that *Sitsqwayk* and *Maiabalaena* are completely edentulous mysticetes that lacked baleen and fed via suction. The authors also questioned the presence of baleen in eomysticetids, some of which show well developed lateral palatal foramina and sulci, laterally bowed mandibles, and an elongate rostrum with nearly complete tooth loss (Fig. 6). The point at which baleen first evolved is therefore murky in the ‘transitional toothless suction feeder’ scenario, but is posited to have occurred after the complete loss of a functional postnatal dentition (Peredo et al. 2018). Mapping of feeding characters on the phylogenetic hypothesis of Peredo et al. (2018) does not support their evolutionary interpretation (see **Results and Discussion: Summaries of phylogenetic trees and character mappings**).

Direct fossil evidence for baleen provides the most compelling evidence for the presence of this keratinous tissue in long extinct organisms, but as noted above, very few specimens record such evidence (e.g., Bisconti 2012), and phylogenetic analyses of skeletal fossils entail speculative reconstructions that hinge upon the relative efficiency or plausibility of different evolutionary explanations (e.g., Peredo et al. 2018 versus Ekdale and Deméré 2022). Researchers have therefore searched for additional clues about the origin of baleen and the loss of postnatal dentition in Mysticeti by examining complementary data from extant species. Although less direct, developmental and molecular evidence from modern taxa provide additional context for interpreting the teeth-to-baleen transition in Mysticeti.

### Ontogeny recapitulates phylogeny in the teeth-to-baleen transition?

Previous studies have tried to reconcile paleontological evidence of the teeth-to-baleen transition with developmental observations on modern species using a qualitative approach (e.g., Deméré et al. 2008; Peredo et al. 2017). Different morphologies, especially apomorphic features of extinct aetiocetids have been interpreted with reference to developmental evidence (e.g. Deméré and Berta 2008; Marx et al. 2016; Fordyce and Marx 2018).

Ontogenetic data document the coexistence of baleen structures and tooth germs based on both anatomical observations (Fig. 11) and gene expression data in fetal mysticetes. Several studies conducted on fetal balaenopterids and balaenids have demonstrated that tooth germs are still recognizable in both upper and lower jaws at the same developmental stage when baleen starts forming in the upper jaw (Ishikawa and Amasaki 1995; Sawamura 2008a, b; Thewissen et al. 2017; Lanzetti 2019). In addition, it has been shown that in the bowhead whale (*Balaena mysticetus*), the growth factor FGF-4 (associated with tooth growth in mammals) is expressed in the tissues surrounding developing baleen, even after all tooth germs are resorbed. This latter pattern reinforces the view that there is an ontogenetic connection between teeth and baleen, with an ancestral developmental pathway for teeth perhaps being co-opted for baleen growth in mysticetes (Thewissen et al. 2017). In Antarctic minke whales (*Balaenoptera bonaerensis*) – the focus of most developmental studies – the formation of baleen rudiments as a denser tissue medial to still clearly recognizable tooth germs is first observed in specimens that are in the last half of gestation (>90 cm in total body length). This dense tissue is linked to external anatomical changes in the paired longitudinal gingival ridges of the palate, which enlarge medially before forming a distinct keel that later in ontogeny becomes the baleen apparatus (Ishikawa and Amasaki 1995; Sawamura 2008a, b; Lanzetti 2019) (Fig. 11).

**Fig. 11.**
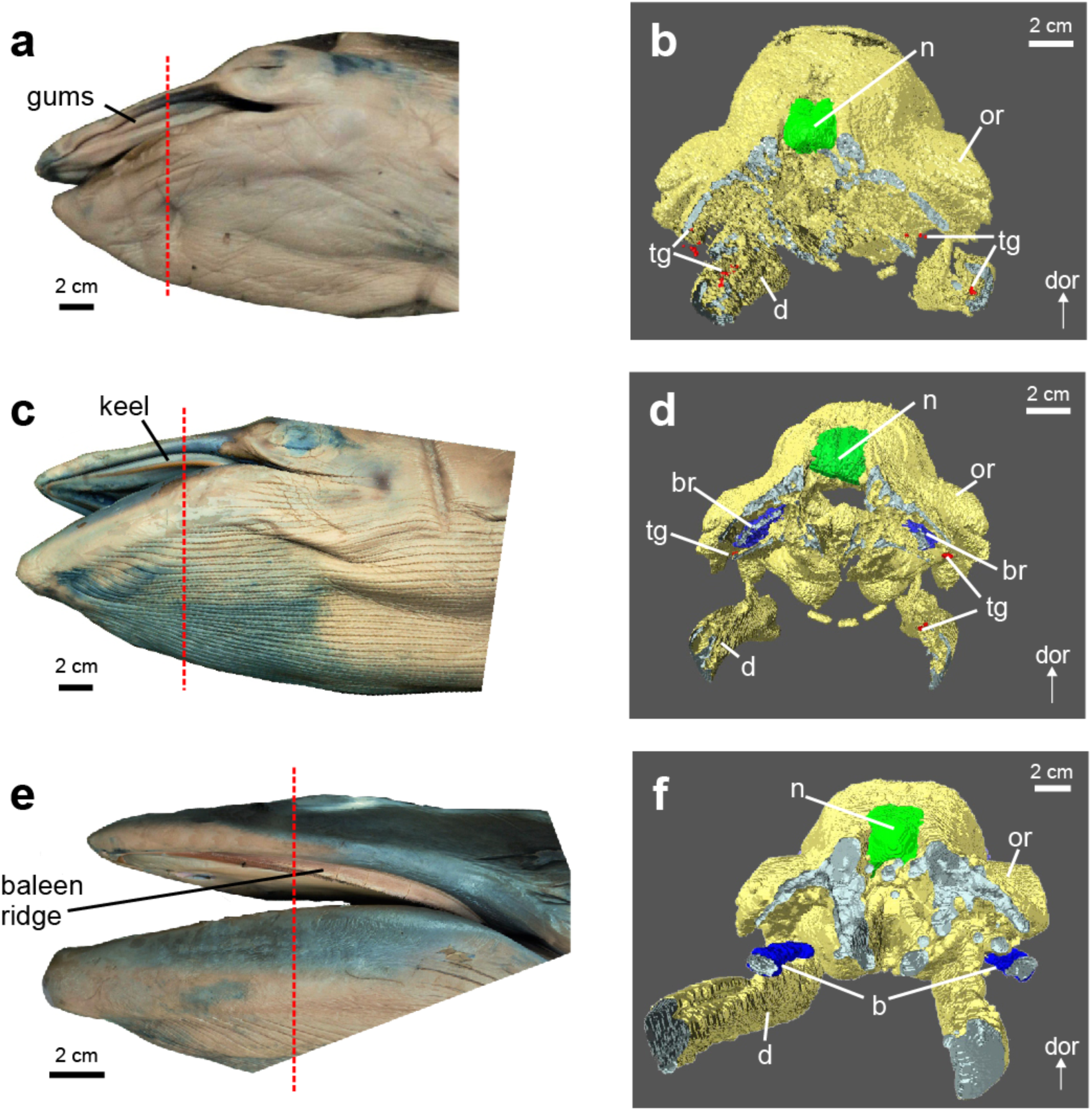
Teeth-to-baleen transition in the ontogeny of the Antarctic minke whale, *Balaenoptera bonaerensis.* **a-c.** “tooth germs” stage (NSMTblue10, TL 74 cm, 5.4 months, 57% gestation), **c-d.** “keel” stage (NSMT27171, TL 115 cm, 6.5 months, 68% gestation), **e-f.** “baleen ridge” stage (NSMT27174, TL 212.5 cm, 8.1 months, 85% gestation). Left column (**a**, **c**, **e**): lateral views of the head, right column (**b**, **d**, **f**): cross sections for 3D renderings of internal head morphology. Red dashed lines in left column show the location and orientation of each cross section to the right. Ossified skull bones (yellow), nasal bones (green), tooth germs (red), baleen rudiments and baleen (blue) are highlighted. Selected elements are labeled (b, baleen; br, baleen rudiments; d, dentary; n, nasal; or, orbit; tg, tooth germs). NSMT = National Museum of Nature and Science, Tsukuba, Japan, TL = total body length, dor = dorsal. Images and data are modified from Lanzetti (2019).

Overall, the ontogenetic series in *Balaenoptera bonaerensis* (Fig. 11) recapitulates the inferred evolutionary transformation in the ‘transitional chimaeric feeder ‘ scenario: from teeth, to teeth plus baleen, to just baleen (see **Results and Discussion: The evolution of baleen and filter feeding**). On the surface, the ontogenetic data are consistent with the hypothesis that both teeth and proto-baleen structures were present in aetiocetids (Deméré and Berta 2008; Deméré et al. 2008; Ekdale and Deméré 2022), as these data demonstrate that baleen and teeth can co-exist anatomically and are deeply linked through shared developmental pathways (Thewissen et al. 2017). Recently, morphological evidence also has been presented for the likely co-occurrence of baleen and vestigial teeth in eomysticetids (Fig. 6; Boessenecker and Fordyce 2015a, b), while Ekdale and Deméré (2022) used CTscanning to show that both the upper teeth and the lateral palatal foramina of the aetiocetid *Aetiocetus weltoni* connect to the superior alveolar canal. The latter result suggests that the neurovascular anatomy of the upper teeth was co-opted for use by baleen (Figs. 4a, 10). In other words, there is a convergence of evidence from ontogeny, paleontology, and phylogeny that is consistent with the ‘transitional chimaeric feeder’ scenario.

Many dental features of aetiocetids, however, do not resemble the tooth germs of prenatal modern mysticetes. This should not be surprising, because the rudimentary enamel-less tooth germs of extant mysticetes have been subjected to long-term relaxed selection since mineralized postnatal teeth were lost >20 million years ago following the elaboration of baleen. Extinct toothed stem mysticetes have heterodont teeth, in contrast with the mostly homodont rudimentary dentition found in modern mysticete fetuses (Karlsen 1962). Most aetiocetids retain tooth counts similar to archaeocetes, the paraphyletic stem group to crown Cetacea, with fewer than 40 teeth (Deméré et al. 2008; Marx et al. 2015; Marx et al. 2016). These plesiomorphic dental features of Aetiocetidae suggested to some researchers that polydonty of the prenatal teeth in extant mysticetes evolved independently in a separate group of early mysticetes. These same researchers go on to conclude that present developmental evidence does not support the co-occurrence of teeth and baleen in aetiocetids (Marx et al. 2016; Peredo et al. 2017; Fordyce and Marx 2018). Since baleen rudiments only appear when tooth germs are resorbing and because polydonty and homodonty are associated with baleen formation in fetal mysticetes, these authors suggested that the ontogenetic series is not consistent with a scenario in which adult aetiocetids possessed both proto-baleen and a well-developed dentition.

While ontogenetic data are meaningful for understanding macroevolutionary change, information from fetuses of modern species should be interpreted judiciously with reference to the adult morphologies of long extinct taxa. Proper evolutionary assessments of ontogeny require interpretations of complete developmental series for different taxa within a rigorous phylogenetic context (Fink 1982; Mabee 1993; Smith 2001). Unless exceptionally well-preserved specimens of fetal aetiocetids are found—an unlikely prospect—it is not possible to determine whether polydonty was expressed during aetiocetid development. It is therefore not particularly informative to compare the number of tooth germs in modern fetuses to the tooth counts of adult fossil specimens that diverged from extant mysticetes >30 million years ago. Higher expression levels of certain growth factors still active in modern species (Thewissen et al. 2017) or fusion of the many single rooted teeth observed in the lower jaw of fetal specimens with the progression to adulthood (e.g. Karlsen 1962) could explain the heterodont dentition seen in aetiocetid fossils, without precluding the possibility of prenatal polydonty and associated baleen tissue in adults.

Moreover, prenatal polydonty and homodonty might have been acquired after the loss of the postnatal dentition in chaeomysticetes as a necessary iterative developmental precursor to baleen rack formation (Thewissen et al. 2017) or simply due to relaxed selection on the dentition. The high tooth counts and unequal numbers of teeth in each side of the jaws observed in some aetiocetids, such as *Aetiocetus polydentatus* and *Salishicetus meadi*, also could be an indicator of relaxed selective pressures and altered developmental constraints in the dentitions of these polydont aetiocetid species (Barnes et al. 1994; Cobourne and Sharpe 2010; Thewissen et al. 2017).

Since modern baleen whales are completely edentulous at birth and already have fully erupted baleen laminae, it is not surprising that tooth germs resorb before birth (e.g. Karlsen 1962; Thewissen et al. 2017; Lanzetti 2019). However, this does not imply that tooth germs *must* be resorbed for baleen to grow, contrary to the assertions of Marx et al. (2016) and Peredo et al. (2017). Based on a more appropriate application of developmental evidence (Fig. 11) and more complete descriptions of extinct taxa (Figs. 6, 10), we see no conflict with the hypothesis that both postnatal teeth and proto-baleen structures were simultaneously present in aetiocetids (Figs. 1d, 9b) as well as in eomysticetids (Fig. 6d).

### Molecular decay of dental genes in crown Mysticeti

Tooth development is a complex process that involves hundreds of genes, most of which are pleiotropic and also perform essential functions outside of odontogenesis (Thesleff 2006). However, ten genes have been shown to be tooth-specific with respect to their crucial functions that are maintained by natural selection (Deméré et al. 2008; McKnight and Fisher 2009; Meredith et al. 2009, 2011, 2013, 2014; Gasse et al. 2012; Kawasaki et al. 2014, 2020; Springer et al. 2015, 2016, 2019; Mu et al. 2021) (Table 2). All of these genes have become pseudogenized in one or more clades of mammals that have lost their postnatal teeth (baleen whales, anteaters, pangolins, Stellar’s sea cow) or lost the enamel on their teeth (armadillos, sloths, aardvarks, pygmy and dwarf sperm whales). The molecular decay of these tooth-specific genes in mysticetes provides a genomic record of tooth loss that can be compared to and integrated with morphological evidence from the fossil record (Fig. 12).

**Fig. 12.**
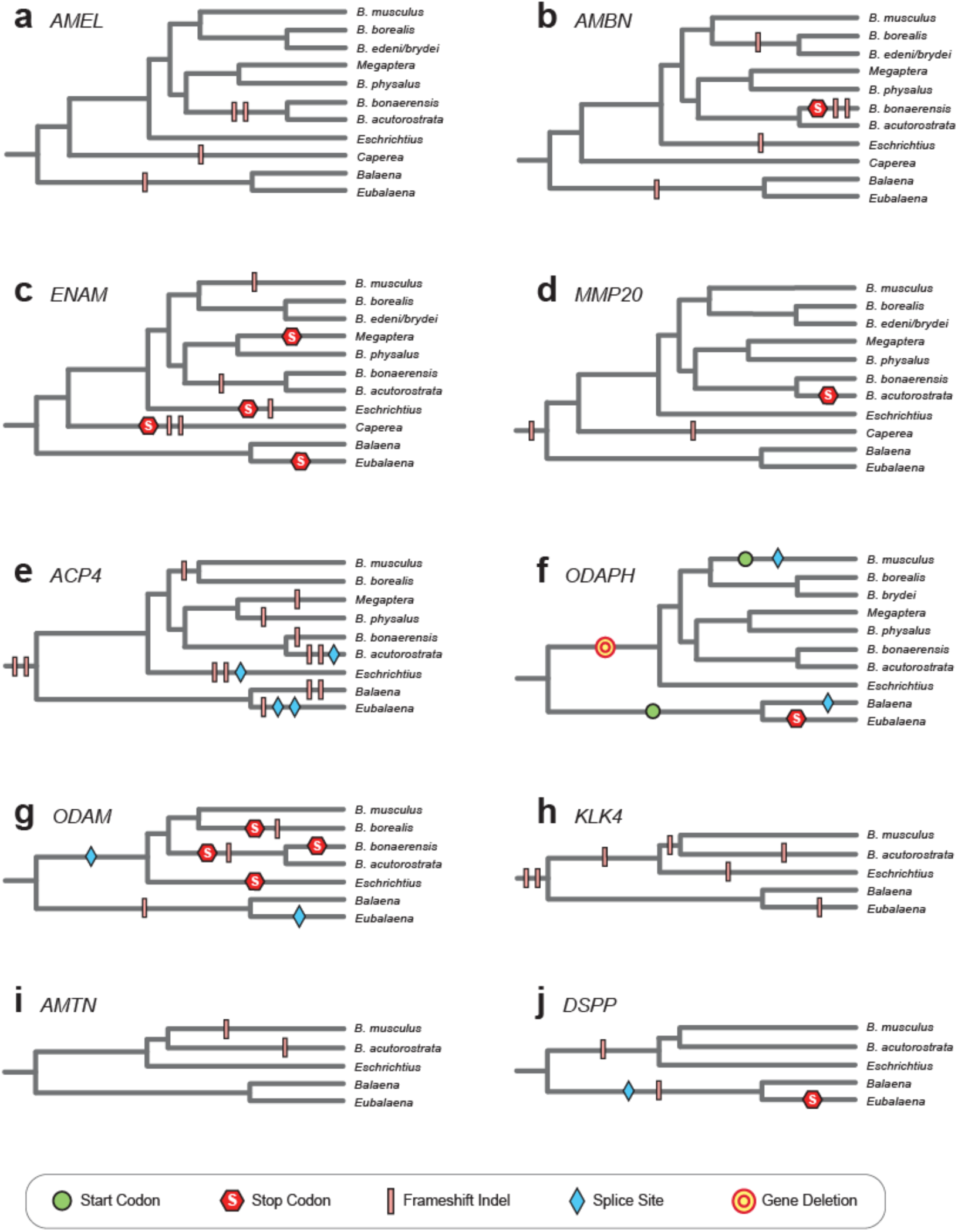
Molecular decay of dental genes in Mysticeti. Mutations in enamel-and tooth-specific loci (**a-j**) that are suggestive of pseudogenization mapped onto a species tree for extant baleen whales (Steeman et al. 2009). Parsimony optimization (DelTran) of various inactivating mutations (see key) are shown. Note the shared mutations that imply gene knockouts prior to the last common ancestor of crown Mysticeti for *MMP20*, *ACP4*, and *KLK4*. For the *ODAPH* (= *C4orf26*) locus in Balaenopteroidea (*Balaenoptera* + *Megaptera* + *Eschrichtius*), one mutant allele includes a large deletion that has completely excised *ODAPH* from the genome in most species, but another allele for this locus lacks the deletion and instead shows an intron splice site mutation and a disrupted start codon in *Balaenoptera musculus* (Springer et al. 2016). ‘*Balaenoptera*’ is abbreviated as ‘*B.*’ in all trees. For some dental genes (e.g., *AMBN*), mapping of inactivating mutations was based on PCR amplification and sequencing of selected exons and not complete protein-coding regions (see Deméré et al. 2008; Meredith et al. 2011; Springer et al. 2016).

**Table 2.**
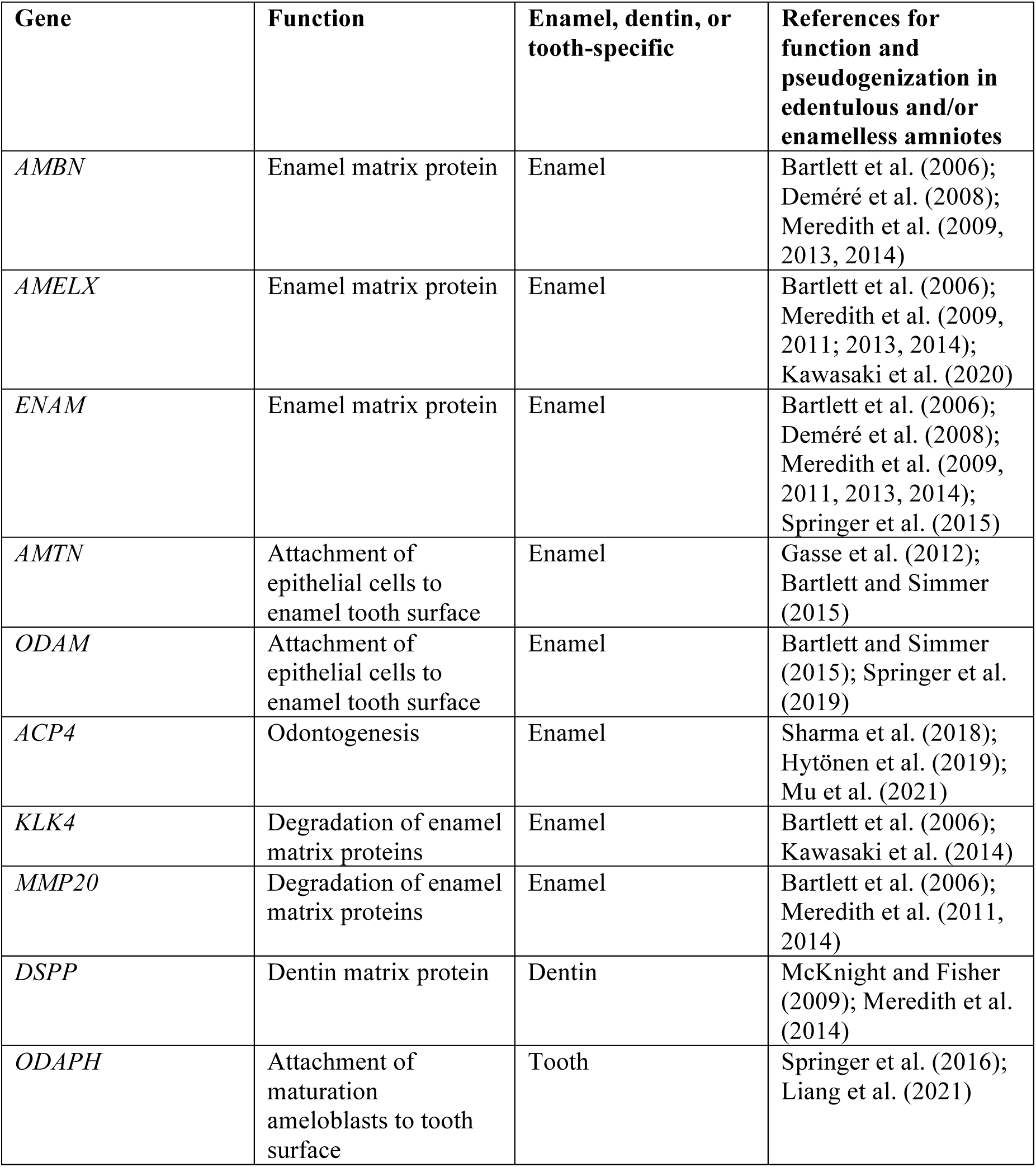
Ten tooth-specific genes and their functions related to the dentition

| **Gene** | **Function** | **Enamel, dentin, or tooth-specific** | **References for function and pseudogenization in edentulous and/or enamelless amniotes** |
| --- | --- | --- | --- |
| *AMBN* | Enamel matrix protein | Enamel | Bartlett et al. (2006); Deméré et al. (2008); Meredith et al. (2009, 2013, 2014) |
| *AMELX* | Enamel matrix protein | Enamel | Bartlett et al. (2006); Meredith et al. (2009, 2011; 2013, 2014); Kawasaki et al. (2020) |
| *ENAM* | Enamel matrix protein | Enamel | Bartlett et al. (2006); Deméré et al. (2008); Meredith et al. (2009, 2011, 2013, 2014); Springer et al. (2015) |
| *AMTN* | Attachment of epithelial cells to enamel tooth surface | Enamel | Gasse et al. (2012); Bartlett and Simmer (2015) |
| *ODAM* | Attachment of epithelial cells to enamel tooth surface | Enamel | Bartlett and Simmer (2015); Springer et al. (2019) |
| *ACP4* | Odontogenesis | Enamel | Sharma et al. (2018); Hytönen et al. (2019); Mu et al. (2021) |
| *KLK4* | Degradation of enamel matrix proteins | Enamel | Bartlett et al. (2006); Kawasaki et al. (2014) |
| *MMP20* | Degradation of enamel matrix proteins | Enamel | Bartlett et al. (2006); Meredith et al. (2011, 2014) |
| *DSPP* | Dentin matrix protein | Dentin | McKnight and Fisher (2009); Meredith et al. (2014) |
| *ODAPH* | Attachment of maturation ameloblasts to tooth surface | Tooth | Springer et al. (2016); Liang et al. (2021) |

Six tooth-specific genes belong to the secretory calcium-binding phosphoprotein (SCPP) gene family: amelogenin (*AMELX*), odontogenic ameloblast associated (*ODAM*), amelotin (*AMTN*), ameloblastin (*AMBN*), enamelin (*ENAM*), and dentin sialophosphoprotein (*DSPP*) (Kawasaki et al. 2011). Three of these SCPP genes (*AMELX, AMBN, ENAM*) encode enamel matrix proteins and *DSPP* encodes a dentin matrix protein. *AMTN* and *ODAM*, in turn, are expressed during the maturation stage of enamel development and help to facilitate the attachment of epithelial cells to the mineralized surface of the tooth (Bartlett and Simmer 2015). Two other tooth-specific genes encode enzymes (matrix metallopeptidase 20 [*MMP20*], kallikrein-related peptidase 4 [*KLK4*]) that break down enamel matrix proteins. Acid phosphatase 4 (*ACP4*) plays a role in odontogenesis although its precise function(s) are poorly understood (Hytönen et al. 2019). Finally, odontogenesis associated phosphoprotein (*ODAPH*) encodes a protein that plays a role in the attachment of maturation ameloblasts to the enamel surface (Liang et al. 2021). However, this gene remains intact in several enamelless taxa that have been investigated (nine-banded armadillo, Hoffmann’s two-toed sloth, aardvark), which suggests that ODAPH is tooth related but not enamel-specific (Springer et al. 2016). The function of these ten genes is summarized in Table 2.

Among the tooth-specific genes, a large CHR-2 SINE retroposon insertion in exon 2 of *MMP20* is shared by all extant mysticetes (Meredith et al. 2011). Mice that are deficient for *MMP20* exhibit amelogenesis imperfecta, a condition in which thin, hypomineralized enamel chips away from the underlying dentin (Caterina et al. 2002; Fanjul-Fernandez et al. 2009). The SINE insertion in *MMP20* that is shared by all extant mysticetes provides direct evidence that the genetic toolkit for enamel production was inactivated on the stem mysticete branch (Fig. 12d). More recently, Mu et al. (2021) provided evidence for the inactivation of a second enamel-related gene, *ACP4* (= *ACPT*), in the common ancestor of all baleen whales. Specifically, all mysticetes that were examined, including representatives of Balaenidae and Balaenopteroidea that diverged at the base of crown Mysticeti, share single-base frameshift deletions in exons 4 and 5 of this gene (Fig. 12e). Four other enamel-related genes (*ODAM, AMBN, ENAM, AMELX*) also show evidence of inactivation in some or all mysticetes that have been investigated (Deméré et al. 2008; Meredith et al. 2009, 2011; Springer et al. 2019), but there are no shared inactivating mutations in the protein-coding exons or intronic splice sites of these genes (Fig. 12a-c, g). *KLK4* has inactivating mutations in *Balaenoptera acutorostrata* (Kawasaki et al. 2014), but other mysticetes have not been examined, and it remains to be determined if any of these mutations are shared with other mysticetes. Also, we are unaware of published studies that report on *AMTN* in mysticetes.

We therefore examined five mysticetes with genome sequences (*Balaena mysticetus, Eubalaena japonica, Balaenoptera acutorostrata, Balaenoptera musculus, Eschrichtius robustus*) to determine if there are any shared inactivating mutations in *KLK4* and/or *AMTN*. In the case of *KLK4*, we discovered a 1-bp frameshift deletion in exon 3 that is shared by all five taxa (Fig. 12h). There is also a 1-bp frameshift deletion in exon 5 that is shared by the three balaenopteroids (*B. acutorostrata, B. musculus, E. robustus*) and *E. japonica*, but not *B. mysticetus*. This deletion could have resulted from independent mutations on the stem Balaenopteroidea and *E. japonica* branches or in the common ancestor of all extant mysticetes if there was incomplete lineage sorting of the intact and deletion alleles. There are also frameshift deletions in exon 1. Two of these are 1-bp deletions that are shared by the two balaenopterids and there is a 2-bp deletion in *E. robustus*. For *AMTN*, there is a 4-bp insertion in exon 4 in *Balaenoptera acutorostrata* and a large deletion that spans exons 7-9 in *B. musculus* (Fig 15i).

It is manifest from shared inactivating mutations in *MMP20*, *ACP4*, and *KLK4* that enamel loss occurred on the stem lineage to crown Mysticeti. By contrast, genomic evidence for the complete absence of teeth on the ancestral lineage of crown mysticetes is so far lacking. McKnight and Fisher (2009) reported partial sequences for exon 4 of *DSPP* in *Balaena mysticetus* and found that the DPP domain encoded by this gene is intact. However, we discovered a 1-bp frameshift insertion in exon 3 of *DSPP* in *B. mysticetus* (scaffold 876). This insertion is shared with another balaenid, *Eubalaena japonica* (RJWP010020556), which also has a mutation in the donor splice site of intron 1 (from GT to AT), a 5-bp frameshift deletion in exon 3, and a premature stop codon in exon 4 that is 1089 bp upstream from the usual stop codon (Fig. 12j). However, an ∼18.3 kb deletion that encompasses exons 1-3 and the 5’ end of exon 4 is present in three balaenopteroids that we examined (*Balaenoptera acutorostrata* [NW_006727686], *Balaenoptera musculus* [VNFD02000988], *Eschrichtius robustus* [NIPP01012418]). It is therefore unclear if the 1-bp deletion in exon 3 of *DSPP* occurred in the ancestor of balaenids or is a shared deletion in all extant mysticetes that was subsequently overprinted by the larger deletion that removed this exon from representative balaenopteroids. *ODAPH* is a second tooth-specific rather than enamel-specific gene that is inactivated in edentulous taxa such as mysticetes and pangolins, but not in enamelless taxa (*Dasypus novemcinctus* [nine-banded armadillo], *Choloepus hoffmanni* [Hoffmann’s two-toed sloth], *Orycteropus afer* [aardvark]) that have been investigated (Springer et al. 2016). We examined additional edentulous and enamelless taxa and found that this gene has four 1-bp frameshift deletions in exon 2 of the edentulous anteater *Tamandua tetradactyla* (PVIF010030063), but is intact in two armadillos (*Tolypeutes matacus* [PVIB010178092, PVIB010180256], *Chaetophractus vellerosus* [PVKH010014510], and a sloth (*Choloepus didactylus* [JADCNF010000003]) that now have genome sequences, consistent with the pattern observed previously (Springer et al. 2016). All mysticetes have inactivating mutations in this gene, but there are no inactivating mutations that are shared by all mysticetes (Springer et al. 2016) (Fig. 12f).

In summary, there is unambiguous molecular evidence for the loss of enamel, but not dentin (and teeth), on the stem mysticete branch. Unlike enamel, where there are eight different genes whose sole or primary function is related to enamel formation or the maintenance of a healthy junctional epithelium that attaches to the enamel surface of teeth, there are only two tooth-specific genes that are inactivated in conjunction with edentulism. Inactivations in these loci can tentatively be interpreted as molecular markers for the complete loss of postnatal teeth (i.e., enamel, dentin). As noted above, one of these genes has a large deletion that may have overprinted evidence for a shared inactivating mutation in all crown mysticetes.

An additional complication is that mysticetes have very slow rates of molecular evolution, perhaps the lowest among extant mammals (Jackson et al. 2009; Meredith et al. 2009). A low substitution rate may have provided limited opportunity for the accumulation of shared inactivating mutations in the few known dentin/tooth-specific genes. Even if *DSPP* and *ODAPH* were under relaxed selection on the stem mysticete branch for millions of years, a very slow rate of evolution might not register any mutations that mark loss of the dentition. Indeed, the majority of enamel-specific genes did not accumulate inactivating mutations on the stem mysticete branch even though enamel production was abrogated on this branch (Meredith et al. 2011). Thus, it is possible that dentin/tooth-specific genes were under relaxed selection on the stem mysticete branch even though there is no genomic evidence for shared inactivating mutations in these genes. An alternate explanation for the absence of shared mutations in dentin/tooth-specific genes (*DSPP, ODAPH*) is that edentulism evolved independently in two or more lineages of mysticetes rather than in their common ancestor.

This latter hypothesis currently is inconsistent with paleontological evidence and ancestral reconstructions of tooth loss in mysticetes (Fitzgerald 2006, 2010; Meredith et al. 2011). However, if small enamelless teeth were set in the gingivae rather than in bony alveoli as in some living beaked whales (Gomerčić et al. 2006), then these teeth could have become dissociated from the rest of the skeleton of an otherwise toothless mysticete by post-mortem taphonomic processes. In this context, there is equivocal evidence for the presence of small teeth at the anterior ends of the upper and lower jaws in at least some Eomysticetidae (Fig. 6; Okazaki 2012; Boessenecker and Fordyce 2015a, b), while prior studies on more fragmentary fossils suggested that members of this lineage were completely edentulous, baleen-bearing mysticetes (Sanders and Barnes 2002).

It is not known whether a partial non-functional postnatal dentition also extended to some early-diverging members of crown clade Mysticeti. Additional paleontological and molecular evolutionary studies will be required to determine if the enamel and dentin components of teeth were lost either: (1) simultaneously on the stem mysticete branch; (2) in a stepwise fashion on the stem mysticete branch (i.e., enamel first, then dentin); or (3) sequentially with enamel loss on the stem mysticete branch followed by independent dentin losses in two or more crown mysticete lineages. From the molecular perspective, one approach to distinguish these three possibilities would be a more comprehensive survey of dentin-specific and tooth-specific loci, including any regulatory regions of these genes and all protein-coding exons. Pseudogenization events on the stem lineage to crown Mysticeti could yield molecular clues regarding ancestral phenotypes and indicate when inactivation of these loci occurred via methods that estimate selection intensities on protein-coding sequences (e.g., Meredith et al. 2009).

For fossil evidence, more completely preserved extinct mysticetes that are close to or just within the crown clade could reveal the presence/absence of vestigial teeth, as well as whether such teeth were enamel-capped. Currently, a large gap in morphology remains between aetiocetids with fully-developed, presumably functional dentitions (Fig. 4) and their toothless (or nearly so) sister group that includes *Maiabalaena*, *Sitsqwayk*, Eomysticetidae (Fig. 6), and crown Mysticeti. It is possible that there was a saltatory evolutionary shift between these divergent dental conditions. Alternatively, reduction of the dentition may have been more gradual, and new fossil discoveries might illuminate key transitional stages. Finally, taxa that emerged just after the basal split in crown Mysticeti might reveal enamel-less vestigial teeth within the crown clade, which would hint that this condition is primitive for crown mysticetes. To date, no evidence of this nature has been published.

### Repurposed keratin genes and the teeth-to-baleen shift

Baleen (Fig. 13) is a neomorphic keratinous structure with unique biomechanical properties, in part due to calcification that enhances stiffness of the flexible keratin plates in seawater (Szewciw et al. 2010). Unlike teeth that are present in most mammalian species, including easily studied model species such as the domestic mouse, baleen is poorly characterized from a genomic perspective. Numerous studies have documented wholesale degradation of tooth genes that relates to the complete evolutionary loss of the ancestral functional dentition in Mysticeti (Fig. 12), but very little molecular-genetic work has focused on emergence of the apomorphic baleen filter. Because baleen is a keratinous epidermal appendage, the *de novo* evolutionary design of this unique mammalian sieve likely entailed a repurposing (exaptation; Gould and Vrba 1982) of existing genes that originally evolved for other functions, but also the derivation of new genes that are baleen-specific.

**Fig. 13.**
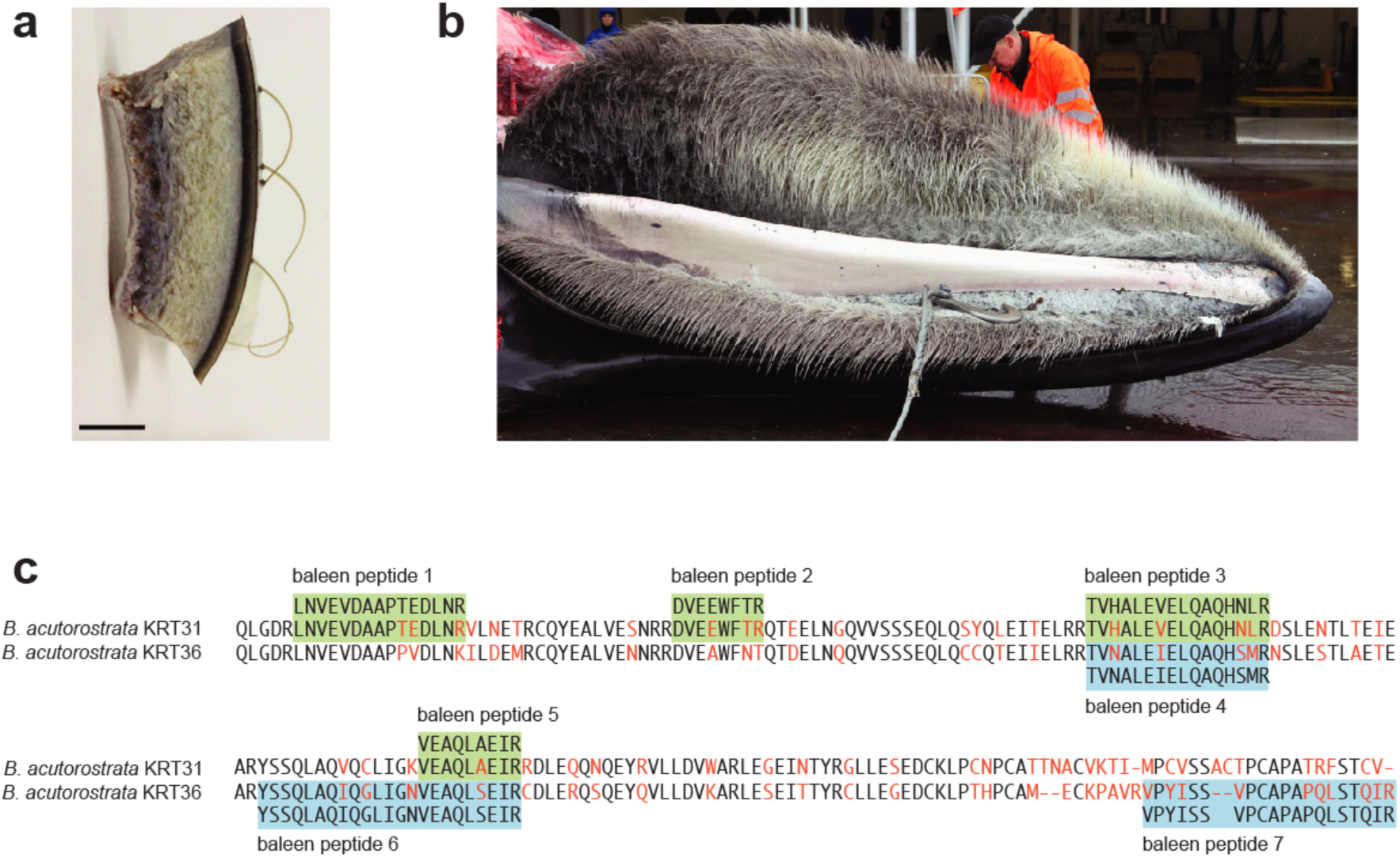
Vibrissae hairs in skin sample of newborn dolphin, palatal view of baleen in fin whale, and evidence for expression of keratin ‘hair’ genes in mysticete baleen plates. Some odontocete species, such as *Tursiops truncatus*, have sparse whiskers on the snout (**a**), but all mysticete cetaceans, such as *Balaenoptera physalus*, have keratinous baleen plates that fray to form an extensive hair-like filter-feeding apparatus (**b**). Short peptides (**c**) derived from the baleen of various mysticete species match amino acid sequences for keratins such as *KRT31* and *KRT36*, demonstrating that these repurposed structural proteins are expressed in baleen (Solazzo et al. 2013, 2017). The partial sequence alignment of KRT31 and KRT36 from minke whale (*Balaenoptera acutorostrata*) shows amino acid differences between keratin paralogs (red). For this alignment region, four baleen peptides perfectly match KRT31 (green) and three perfectly match KRT36 (blue). Scale bar in **a** is 5 mm. Photo of dolphin skin is by R. Ramos, and photo of fin whale is by A. Aguilar.

In mammals, the main structural components of keratinized structures - such as pelage hair, vibrissae (whiskers), nails, hooves, skin, and tongue papillae - are keratin proteins (Hesse et al. 2004; Schweizer et al. 2006; Bragulia et al. 2009) and keratin associated proteins (KRTAPs) (Wu et al. 2008). In hair, keratin filaments are surrounded by a matrix that consists of KRTAPs. The genes that encode keratins and KRTAPs comprise two gene families that are highly degraded in cetaceans relative to terrestrial mammals, likely correlated with the evolutionary loss of hooves, pelage hair, and the epidermal barrier of the skin that came with the shift from terrestrial to aquatic habitats (Khan et al. 2014; Nery et al. 2014; Oh et al. 2015; Sun et al. 2017; Sharma et al. 2018; Ehrlich et al. 2019; Espregueira Themudo et al. 2020; Springer et al. 2021). Although greatly reduced, some cetaceans (including mysticetes; Berta et al. 2015; Drake et al. 2015) retain sparse vibrissae (Fig. 13a), so the molecular components of hair are retained, even in some extant odontocete cetaceans that never evolved baleen (Slipjer 1976; Ling 1977; Mynett et al 2022; Springer et al. 2021).

Peptides derived from baleen-plate samples offer an initial glimpse at ‘hair genes’ that have been repurposed by the baleen-producing palatal epithelium of mysticetes (Fig. 13c). Solazzo et al. (2013, 2017) used short peptide sequences as molecular signatures to differentiate the baleen of different mysticete species from archaeological artifacts. Amino acid sequences were matched to translations of genes in published genome assemblies, and identities to multiple hair keratin paralogs (KRT31, KRT33A, KRT36, KRT81, KRT83, KRT85, KRT86) were revealed in the baleen samples (Solazzo et al. 2017). The peptide sequences suggest that baleen whales do in fact feed by using a mouthful of repurposed ‘hair’ that has been modified to strain small organisms from seawater.

Characterization of baleen at the genomic level is in its infancy, however, and more comprehensive proteomic surveys or RNA-sequencing studies of baleen-producing epithelia will be required to document the full catalog of genes that are expressed in baleen. Given the basic structure of baleen, which includes hair-like bristles embedded in an amorphous medullary keratin matrix that yields the distinctive medial fringe of the baleen plate upon abrasive wear (Fudge et al. 2009), we predict that KRTAP proteins will be components of baleen as well. Strictly baleen-specific loci that evolved synchronously with the emergence of this novel feeding structure are currently unknown. If such genes exist, it might be possible to date the origin of baleen by molecular clock estimates that specify the divergence of baleen-specific loci from their closest genetic relatives (paralogs). An analogous approach has been used to infer when feathers or feather-like structures evolved on the stem lineage to Aves (birds) based on the documented amplification of beta keratin genes (Greenwold and Sawyer 2010; Li et al. 2013; Lowe et al. 2015). Mysticete genomes do not indicate extensive keratin-gene duplications associated with the evolution of baleen (Nery et al. 2014; Sun et al. 2017; Ehrlich et al. 2019), but KRTAPs (Khan et al. 2014) and other genes that might be expressed in baleen tissues merit further study, in particular using more complete genome assemblies for mysticetes. Given preliminary genetic evidence for an association between tooth and baleen development (Thewissen et al. 2017), it also will be of interest to test whether the same genes are responsible for both mineralization of the dentition and calcification of baleen, or whether new genes evolved with the elaboration of baleen that have yielded its unique biomechanical properties (Szewciw et al. 2010).

## Conclusion

Remarkably, basal relationships of mysticete whales differ for each phylogenetic study reviewed here, as do many homology decisions for key traits that are central to the feeding apparatus (Figs. 3, 5, 7-8). The challenge is that these same traits related to filter feeding in modern taxa provide critical character support at the most important nodes on the stem lineage of Mysticeti. Just a few discrepancies in how observations of specimens are encoded into characters or subtle differences in taxon sampling can rearrange the tree topologies and downstream interpretations of homology and character evolution. Even in some of the most recent comprehensive studies, surprising results are supported in parsimony analyses of morphological characters, including paraphyly of Mysticeti, paraphyly of Chaeomysticeti, and non-monophyly of Kinetomenta (Aetiocetidae + Chaeomysticeti) that imply extensive homoplasy in key feeding characters (Figs. 5b, 7-8; Appendix 1).

Overall however, comparative research on Mysticeti to this point generally has corroborated the close relationship of aetiocetids (Fig. 4) and eomysticetids (Fig. 6) to crown Mysticeti (Fig. 9a). Given anatomical evidence for the simultaneous occurrence of both teeth and baleen in these stem mysticetes, recent ontogenetic observations, and genomic data, we contend that the simplest historical reconstruction of the teeth-to-baleen transition is represented by the ‘transitional chimaeric feeder’ scenario (Fig. 9b). For phylogenetic hypotheses that support Kinetomenta, the ‘transitional chimaeric feeder’ scenario requires fewer inferred changes in anatomy, behavior, and function relative to scenarios that posit suction-feeding intermediates that lacked baleen yet already possessed several anatomical features related to filter-feeding (in modern baleen whales) that instead were presumably first used for alternative purposes.

Despite grand phylogenetic efforts (Figs. 3, 5, 7-8), many challenges still remain to be resolved in the coming years by the discovery and description of especially well-preserved fossil specimens (e.g., Fitzgerald 2006; Lambert et al. 2017; Geisler et al. 2017), more detailed analyses of particular character systems (e.g., Hocking et al 2017b; Deméré and Ekdale 2022), quantification of isotopes and enamel structure that can help reconstruct paleodiets (Clementz et al. 2014; Loch et al. 2020), as well as expanded ontogenetic (e.g., Thewissen et al. 2017), genomic, and proteomic work (e.g., Árnason et al. 2017; Solazzo et al. 2017; Ehrlich et al. 2019). Even given extensive new data, however, we predict that the evolutionary history of baleen will remain contentious among some specialists in the field. While it is clear that the discovery of direct fossil evidence for baleen plates or ‘proto-baleen’ bristles in long extinct stem mysticetes would be the most decisive data, evaluations of the relative support for competing hypotheses are based on currently available data. In the case of the teeth-to-baleen transition, the various lines of evidence presented in support of the ‘transitional chimaeric feeder’ scenario, although not definitive, represent the most parsimonious current solution.

## Acknowledgments

Funding was provided by NSF DEB-1457735 (to J.G. and M.S.). We thank Carl Buell, Alberto Gennari, Christian de Muizon, and Robert Boessenecker for artwork and Raul Ramos, Alex Aguilar, Ewan Fordyce, Maksim Plikus, and Robert Boessenecker for photographs.

## Declarations

**Funding:** Funding was provided by NSF DEB-1457735 to J.G. and M.S.

## Availability of data and material

All phylogenetic trees generated in this study are included in the published article.

## Code availability

not applicable

## Additional declarations for articles that report results of studies involving humans and/or animals

**not applicable**

## Appendix 1. Number of taxa, number of characters, minimum tree length, number of minimum length trees, and strict consensus of minimum length trees for 11 morphological character matrices for Mysticeti (2006-2019)

Fitzgerald (2006): 26 taxa, 266 characters (217 parsimony informative) Minimum tree length: 997

Number of minimum length trees: 1

Notes: Our parsimony analysis of the data matrix extracted from supplementary online materials did not yield three minimum length trees as in Fitzgerald (2006), perhaps due to analysis using a more recent version of PAUP* that does not accommodate missing and inapplicable character states using the ‘gapmode = newstate’ option with ordered characters. The single tree produced in our analysis is fully resolved but fully compatible with the strict consensus of three trees from the original publication. Given this discrepancy, we mapped feeding characters on the published strict consensus tree (Fitzgerald 2006).

single minimum length tree in Newick format: (Sus,Hippopotamus,(Georgiacetus,((Zygorhiza,(((Mammalodon,(Chonecetus,(Aetiocetus,((Eomysticetus,Micromysticetus),(Diorocetus,(Pelocetus,((Caperea,Eubalaena),(Balaenoptera,Eschrichtius)))))))),Janjucetus),(ChM_PV5720,ChM_PV4745))),((Archaeodelphis,Xenorophus),(Agorophi us,(Squalodon,(Platanista,((Mesoplodon,Physeter),Tursiops))))))));

Deméré et al. (2008): 36 taxa, 115 characters (113 parsimony informative)

Minimum tree length: 406

Number of minimum length trees: 17

Strict consensus in Newick format:

((((Physeter_macrocephalus,Ziphiidae),Squalodon_calvertensis),Agorophius),(((((((((((Megaptera _novaeangliae,((Balaenoptera_physalus,((Balaenoptera_borealis,(Balaenoptera_bonaerensis,Bala enoptera_acutorostrata)),Balaenoptera_edeni_brydei)),Balaenoptera_musculus)),Megaptera_huba chi),Megaptera_miocaena),((Eschrichtius_robustus,SDSNH_90517),Balaenoptera_gastaldii)),Par abalaenoptera_baulinensis),(Cetotherium_rathkii,Mixocetus_elysius)),(Caperea_marginata,(Balae na_mysticetus,Eubalaena)),Aglaocetus_patulus,Cophocetus_oregonensis,Diorocetus_hiatus,Isana cetus_laticephalus,Parietobalaena_palmeri,Pelocetus_calvertensis),Eomysticetus_whitmorei),(Aet iocetus_weltoni,Aetiocetus_cotylalveus,Aetiocetus_polydentatus,Chonecetus_goedertorum)),Ma mmalodon_colliveri),Janjucetus_hunderi),Zygorhiza_kochii);

Fitzgerald (2010): 22 taxa, 123 characters (120 parsimony informative)

Minimum tree length: 451

Number of minimum length trees: 1

single minimum length tree in Newick format: (Georgiacetus,((Basilosaurus,Dorudon),Zygorhiza),((((Llanocetus,(Mammalodon,Janjucetus)),((( Chonecetus_sookensis,Chonecetus_goedertorum),((Aetiocetus_cotylalveus,Aetiocetus_weltoni), Aetiocetus_polydentatus)),(Eomysticetus,(Micromysticetus,(Diorocetus,(Pelocetus,(Eubalaena,Balaenoptera))))))),(ChM_PV5720,ChM_PV4745)),(Simocetus,Waipatia)));

Marx (2011): 56 taxa, 150 characters (150 parsimony informative)

Minimum tree length: 375

Number of minimum length trees: 9 Strict consensus in Newick format:

(Zygorhiza_kochii,(Patriocetus_ehrlichii,Physeter_catodon),((((Aetiocetus_cotylalveus,Aetiocetu s_polydentatus,Aetiocetus_weltoni),Chonecetus_goedertorum),(Janjucetus_hunderi,Mammalodo n_colliveri)),((((((Aglaocetus_moreni,(((((((((((Archaebalaenoptera_castriarquati,(Megaptera_hu bachi,(Megaptera_miocaena,Megaptera_novaeangliae)),Protororqualus_cuvierii),Parabalaenopter a_baulinensis),(((((Balaenoptera_acutorostrata,Balaenoptera_bonaerensis),Balaenoptera_borealis) ,Balaenoptera_physalus),((Balaenoptera_musculus,Balaenoptera_omurai),Balaenoptera_siberi)), Balaenoptera_edeni)),(Eschrichtioides_gastaldii,Eschrichtius_robustus)),Caperea_marginata),((A ulocetus_latus,Cetotherium_megalophysum,((Cetotherium_rathkei,Herpetocetus_transatlanticus), Nannocetus_eremus),Metopocetus_durinasus,Metopocetus_vandelli,Piscobalaena_nana),Mixocet us_elysius),(Cephalotropis_nectus,Pinocetus_polonicus)),Titanocetus_sammarinensis),Isanacetus_laticephalus),Pelocetus_calvertensis),Cophocetus_oregonensis),Uranocetus_gramensis)),Aglaoc etus_patulus),Diorocetus_chichibuensis,Parietobalaena_palmeri,Parietobalaena_yamaokai),Dioro cetus_hiatus),((Balaena_ricei,Balaena_mysticetus),(Balaenella_brachyrhynus,(Balaenula_astensis ,Eubalaena_glacialis,Eubalaena_shinshuensis)))),(Eomysticetus_whitmorei,Mauicetus_lophoceph alus))));

Boessenecker and Fordyce (2015b): 74 taxa, 363 characters (358 parsimony informative)

Minimum tree length: 1407

Number of minimum length trees: 180

Strict consensus in Newick format:

(Zygorhiza_kochii,((Waipatia_maerewhenua,Simocetus_rayi),(((((Aetiocetus_cotylalveus,(Aetio cetus_polydentatus,Aetiocetus_weltoni)),((((((((((Aglaocetus_moreni,Uranocetus_gramensis),Co phocetus_oregonensis),((Aglaocetus_patulus,Diorocetus_chichibuensis),(Parietobalaena_palmeri, Parietobalaena_campiniana),Parietobalaena_yamaokai,Pelocetus_calvertensis),Diorocetus_hiatus),Isanacetus_laticephalus),Titanocetus_sammarinensis),((Archaebalaenoptera_castriarquati,((((((( (Balaenoptera_acutorostrata,Balaenoptera_bonaerensis,Balaenoptera_borealis,Balaenoptera_berta e),(Balaenoptera_edeni,((Balaenoptera_musculus,Balaenoptera_physalus),Balaenoptera_omurai))),((Balaenoptera_siberi,Plesiobalaenoptera_quarantellii),Diunatans_luctoretmurgo)),Parabalaenop tera_baulinensis),(Megaptera_miocaena,Megaptera_novaeangliae)),Balaenoptera_cortesi_var_po rtisi),Megaptera_hubachi),Protororqualus_cuvierii)),(Eschrichtioides_gastaldii,Eschrichtius_robu stus,Gricetoides_aurorae))),((((Cetotherium_rathkii,Cetotherium_riabinini),Kurdalogonus_mched lidzei,Vampalus_sayasanicus),Metopocetus_durinasus),((((Nannocetus_eremus,(Herpetocetus_tra nsatlanticus,(Herpetocetus_bramblei,(Herpetocetus_morrowi,Herpetocetus_sendaicus)))),Piscoba laena_nana),Herentalia_nigra),Brandtocetus_chongulek))),Joumocetus_shimizui),(((Balaena_rice i,Balaena_mysticetus,Balaenula_astensis,Eubalaena_glacialis,Eubalaena_shinshuensis),Balaenell a_brachyrhynus),(Caperea_marginata,Miocaperea_pulchra))),((’Tohoraata_spp.’,(Tokarahia_kaua eroa,Tokarahia_lophocephalus)),Yamatocetus_canaliculatus,Micromysticetus_rothauseni,Eomyst icetus_whitmorei))),Chonecetus_goedertorum),(Janjucetus_hunderi,Mammalodon_colliveri)),(Ch M_PV_4745,ChM_PV_5720))),(Dorudon_atrox,Basilosaurus_spp));

Marx and Fordyce (2015): 90 taxa, 272 characters (272 parsimony informative)

Minimum tree length: 1233

Number of minimum length trees: 38880

Strict consensus in Newick format:

(Zygorhiza_kochii,(Archaeodelphis_patrius,(Waipatia_maerewhenua,Physeter_catodon)),(((((((( Aetiocetus_cotylalveus,Aetiocetus_polydentatus,Aetiocetus_weltoni),(Chonecetus_sookensis,Mo rawanocetus_yabukii),OCPC_1178),(Chonecetus_goedertorum,UWBM_84024)),(Janjucetus_hun deri,(Mammalodon_colliveri,OU_22026))),OU_GS10897),((((((((Aglaocetus_moreni,Mauicetus_ parki,ZMT_67),((Aglaocetus_patulus,((((Archaebalaenoptera_castriarquati,Parabalaenoptera_bau linensis),(((((Balaenoptera_acutorostrata,Balaenoptera_bonaerensis),Balaenoptera_borealis,Balae noptera_musculus,Balaenoptera_omurai,Balaenoptera_physalus,Balaenoptera_siberi),NMNZ_M M001630),Megaptera_novaeangliae),(Diunatans_luctoretemergo,(Megaptera_hubachi,Plesiobala enoptera_quarantellii))),Balaenoptera_bertae,(Balaenoptera_portisi,((Eschrichtioides_gastaldii,Es chrichtius_robustus),Gricetoides_aurorae)),Balaenoptera_ryani,Megaptera_miocaena),Pelocetus_ calvertensis,Uranocetus_gramensis),(Diorocetus_hiatus,Thinocetus_arthritus))),((((Diorocetus_chichibuensis,Parietobalaena_campiniana,(Parietobalaena_palmeri,Pinocetus_polonicus),(’Parietoba laena_sp.’,Tiphyocetus_temblorensis),Parietobalaena_yamaokai),Isanacetus_laticephalus),OU_22 705),Diorocetus_shobarensis))),Joumocetus_shimizui),Titanocetus_sammarinensis),(((Brandtocet us_chongulek,Cetotherium_rathkii,Cetotherium_riabinini,Kurdalagonus_mchedlidzei),(((Caperea_marginata,Miocaperea_pulchra),((Herpetocetus_bramblei,Herpetocetus_morrowi,’Herpetocetus_ sp.’,Herpetocetus_transatlanticus),Nannocetus_eremus)),Cephalotropis_coronatus)),(Cetotherium_megalophysum,(Metopocetus_durinasus,Piscobalaena_nana)))),((((((Balaena_montalionis,Balae na_mysticetus,Balaena_ricei),Eubalaena_belgica,(Eubalaena_shinshuensis,’Eubalaena_spp.’)),Bal aenula_astensis),Balaenella_brachyrhynus),’Balaenula_sp.’),Morenocetus_parvus,Peripolocetus_v exillifer)),OU_22224),(((Eomysticetus_whitmorei,Yamatocetus_canaliculatus),OU_22044),Micr omysticetus_rothauseni))),Llanocetus_denticrenatus),ChM_PV4745));

Lambert et al. (2017): 38 taxa, 272 characters (265 parsimony informative)

Minimum tree length: 912

Number of minimum length trees: 2

Strict consensus in Newick format:

(Cynthiacetus_peruvianus,((Simocetus_rayi,Physeter_macrocephalus),Archaeodelphis_patrius),( Mystacodon_selenensis,(((Janjucetus_hunderi,Mammalodon_colliveri),(Chonecetus_goedertoru m,(Aetiocetus_polydentatus,Aetiocetus_cotylalveus,Aetiocetus_weltoni))),(ChM_PV4745,(Llano cetus_denticrenatus,((ChM_PV5720,((Morenocetus_parvus,(Balaena_mysticetus,’Eubalaena_spp. ‘)),(((((Caperea_marginata,Miocaperea_pulchra),(Herpetocetus_morrowi,(Herpetocetus_transatla nticus,Herpetocetus_bramblei))),Piscobalaena_nana),(Cetotherium_rathkii,Cetotherium_riabinini)),(Diorocetus_hiatus,((Aglaocetus_patulus,(Aglaocetus_moreni,(Parietobalaena_campiniana,Pari etobalaena_palmeri))),(Pelocetus_calvertensis,(Eschrichtius_robustus,(Megaptera_novaeangliae,( Balaenoptera_acutorostrata,Balaenoptera_physalus))))))))),(Micromysticetus_rothauseni,(Yamato cetus_canaliculatus,Eomysticetus_whitmorei))))))));

Geisler et al. (2017): 53 taxa, 206 characters (204 parsimony informative)

Minimum tree length: 941

Number of minimum length trees: 168 Strict consensus in Newick format:

((Basilosaurus,(Dorudon_atrox,(Zygorhiza_kochii,((Simocetus_rayi,(((Squalodon_calvertensis,(P hyseter_macrocephalus,Ziphiidae)),Waipatia_maerewhenua),Agorophius_pygmaeus)),((ChM_P V5720,Coronodon_havensteini),Metasqualodon_symmetricus,((Llanocetus_denticrenatus,ZMT_ 62),((Fucaia_buelli,((((Aetiocetus_weltoni,Aetiocetus_cotylalveus,Aetiocetus_polydentatus),Fuc aia_goedertorum),Chonecetus_sookensis),(Janjucetus_hunderi,Mammalodon_colliveri))),((Eomy sticetus_whitmorei,(’Tohoraata_spp.’,’Tokarahia_spp.’,Waharoa_ruwhenua),Micromysticetus,Ya matocetus_canaliculatus),(Diorocetus_hiatus,(Pelocetus_calvertensis,(Parietobalaena_palmeri,((A glaocetus_patulus,Isanacetus_laticephalus),((Caperea_marginata,(Balaena_mysticetus,Eubalaena)),(Cetotherium_rathkii,Mixocetus_elysius),Cophocetus_oregonensis,(((Eschrichtius_robustus,SD SNH_90517),Balaenoptera_gastaldii),Megaptera_miocaena,Parabalaenoptera_baulinen,(Megapte ra_hubachi,(Megaptera_novaeangliae,Balaenoptera_physalus,(Balaenoptera_musculus,(Balaenop tera_borealis,Balaenoptera_edeni_brydei,Balaenoptera_bonaerensis,Balaenoptera_acutorostrata)))))))))))))))))):0.5,Georgiacetus_vogtlensis:0.5);

Peredo et al. (2018): 86 taxa, 363 characters (359 parsimony informative)

Minimum tree length: 1587

Number of minimum length trees: 630

Notes: the same data matrix was used in Peredo and Pyenson (2018).

Strict consensus in Newick format:

(Zygorhiza_kochii,((Waipatia_maerewhenua,Simocetus_rayi),((((((((((((((Aglaocetus_moreni,Ur anocetus_gramensis),Cophocetus_oregonensis),((Aglaocetus_patulus,Diorocetus_chichibuensis),( Parietobalaena_palmeri,Parietobalaena_campiniana),Parietobalaena_yamaokai,Pelocetus_calvert ensis),Diorocetus_hiatus),Isanacetus_laticephalus),Titanocetus_sammarinensis),((Archaebalaeno ptera_castriarquat,((((((((Balaenoptera_acutorostrata,Balaenoptera_bonaerensis,Balaenoptera_bor ealis,Balaenoptera_bertae),(Balaenoptera_edeni,((Balaenoptera_musculus,Balaenoptera_physalus),Balaenoptera_omurai))),((Balaenoptera_siberi,Plesiobalaenoptera_quarantellii),Diunatans_lucto retmurgo)),Parabalaenoptera_baulinensis),(Megaptera_miocaena,Megaptera_novaeangliae)),Bala enoptera_cortesi_var_por),Megaptera_hubachi),Protororqualus_cuvierii)),(Eschrichtioides_gastal dii,Eschrichtius_robustus,Gricetoides_aurorae))),(((((Cetotherium_rathkii,Cetotherium_riabinini), Kurdalogonus_mchedlidzei,Vampalus_sayasanicus),Metopocetus_durinasus),((((Nannocetus_ere mus,(Herpetocetus_transatlanticus,(Herpetocetus_bramblei,(Herpetocetus_morrowi,Herpetocetus_sendaicus)))),Piscobalaena_nana),Herentalia_nigra),Brandtocetus_chongulek)),Joumocetus_shi mizui)),(((Balaena_ricei,Balaena_mysticetus,Balaenula_astensis,Eubalaena_glacialis,Eubalaena_ shinshuensis),Balaenella_brachyrhynus),(Caperea_marginata,Miocaperea_pulchra))),Horopeta_u marere),(((’Tohoraata_spp.’,(Tokarahia_kauaeroa,Tokarahia_lophocephalus),Waharoa_ruwhenua),Matapanui_waihao),Yamatocetus_canaliculatus,Micromysticetus_rothauseni,Eomysticetus_whit morei)),(Sitsqwayk_cornishorum,USNM_314627)),((Aetiocetus_cotylalveus,(Aetiocetus_polyde ntatus,Aetiocetus_weltoni)),((Fucaia_buelli,Fucaia_goedertorum),UWBM_82941),Salishacetus_ meadi,((Aetiocetus_tomitai,(Chonecetus_sookensis,Morawanocetus_yabukii)),UWBM_87135))), (Janjucetus_hunderi,Mammalodon_colliveri)),(ChM_PV_4745,ChM_PV_5720))),(Dorudon_atro x,’Basilosaurus_spp.’));

Fordyce and Marx (2018): 106 taxa, 275 characters (275 parsimony informative)

Minimum tree length: 1349

Number of minimum length trees: 34304

Notes: Parsimony analysis of the morphological characters did not support monophyly of Mysticeti. We therefore utilized the Bayesian consensus tree based on morphology and molecules from Fordyce and Marx (2018) for mapping feeding characters in Fig. 8. Both the parsimony strict consensus tree for morphology and the Bayesian tree for combined data are given below.

Parsimony strict consensus in Newick format: (Zygorhiza_kochii,(((Albertocetus_meffordorum,Archaeodelphis_patrius),Olympicetus_avitus,( Waipatia_maerewhenua,Physeter_macrocephalus)),(((((((((Aetiocetus_cotylalveus,Aetiocetus_po lydentatus),Aetiocetus_weltoni),OCPC_1178),(Fucaia_buelli,Fucaia_goedertorum)),((((Aglaocet us_moreni,Aglaocetus_patulus,(((((Archaebalaenoptera_castriarquati,((((Balaenoptera_acutorostr ata,Balaenoptera_bonaerensis),(Balaenoptera_borealis,((Balaenoptera_musculus,NMNZ_MM001 630),Balaenoptera_physalus))),Balaenoptera_omurai),Balaenoptera_siberi)),Megaptera_novaean gliae),Incakujira_anillodefuego),Parabalaenoptera_baulinensis),Balaenoptera_bertae,(Balaenopter a_portisi,((Eschrichtioides_gastaldii,Eschrichtius_robustus),Gricetoides_aurorae)),Balaenoptera_r yani,(Diunatans_luctoretemergo,Fragilicetus_velponi,Megaptera_hubachi,Plesiobalaenoptera_qu arantellii),Megaptera_miocaena),(((((Aulocetus_latus,(((Caperea_marginata,Miocaperea_pulchra),((((Herpetocetus_bramblei,Herpetocetus_morrowi),(’Herpetocetus_sp.’,NMNS_PV19540)),Herp etocetus_transatlanticus),Nannocetus_eremus)),Cephalotropis_coronatus)),((Brandtocetus_chong ulek,(Cetotherium_rathkii,Cetotherium_riabinini)),Kurdalagonus_mchedlidzei)),((Metopocetus_d urinasus,Metopocetus_hunteri),Piscobalaena_nana)),Herentalia_nigra),(Cetotherium_megalophys um,Metopocetus_vandelli)),((((((Balaena_montalionis,Balaena_mysticetus,Balaena_ricei),(Eubal aena_ianitrix,(Eubalaena_shinshuensis,’Eubalaena_spp.’))),Balaenula_astensis),Balaenella_brach yrhynus),’Balaenula_sp.’),Morenocetus_parvus,Peripolocetus_vexillifer),((Diorocetus_chichibuen sis,Parietobalaena_campiniana,(Parietobalaena_palmeri,Pinocetus_polonicus),(’Parietobalaena_sp.’,Tiphyocetus_temblorensis),Parietobalaena_yamaokai),Isanacetus_laticephalus),(Diorocetus_hiatus,Thinocetus_arthritus),Diorocetus_shobarensis,Joumocetus_shimizui,Mauicetus_parki,OU_22 705,Pelocetus_calvertensis,Titanocetus_sammarinensis,Tiucetus_rosae,Uranocetus_gramensis,Z MT_67),Horopeta_umarere),OU_22224,Whakakai_waipata),((Eomysticetus_whitmorei,Yamatoc etus_canaliculatus),(Matapanui_waihao,(Tohoraata_raekohao,Tokarahia_kauaeroa,Waharoa_ruw henua)),Micromysticetus_rothauseni))),(Chonecetus_sookensis,Morawanocetus_yabukii)),OU_G S10897),Coronodon_havensteini),(Janjucetus_hunderi,(Mammalodon_colliveri,Mammalodon_ha kataramea)))),(Llanocetus_denticrenatus,Mystacodon_selenensis));

Bayesian combined data tree in Newick format: (Zygorhiza_kochii:0.005936742,(((Albertocetus_meffordorum:0.004762457,Archaeodelphis_patr ius:0.003562445):0.007547998,(Waipatia_maerewhenua:0.005858093,Physeter_macrocephalus: 0.02865823):0.01802253):0.001979787,Olympicetus_avitus:0.003430435):0.008846279,((((((((((Aetiocetus_cotylalveus:0.003828212,Aetiocetus_polydentatus:0.003110149):0.001222006,Aetio cetus_weltoni:0.001881723):0.006130564,OCPC_1178:0.007467242):0.002697012,(Fucaia_buelli:0.002847145,Fucaia_goedertorum:0.001432189):0.007683302):0.005322719,Chonecetus_sook ensis:0.001176503):0.002593584,(((((((Aglaocetus_moreni:0.001855734,Mauicetus_parki:0.003 289156):0.001678701,ZMT_67:0.0062494):0.004657639,((((Aglaocetus_patulus:0.003226881,(((((Archaebalaenoptera_castriarquati:0.01403892,Parabalaenoptera_baulinensis:0.00636043):0.00 3914812,(((Balaenoptera_acutorostrata:0.01353481,Balaenoptera_bonaerensis:0.008492146):0.0 1062932,Balaenoptera_bertae:0.01355161):0.01129957,((((Balaenoptera_borealis:0.01661122,Ba laenoptera_omurai:0.02697021):0.003045662,(Balaenoptera_musculus:0.01548804,NMNZ_MM 001630:0.003525332):0.005831321):0.004344853,(((Balaenoptera_physalus:0.02115366,Megaptera_novaeangliae:0.01692724):0.003492104,Incakujira_anillodefuego:0.01111955):0.006540906,((Balaenoptera_portisi:0.003386709,((Eschrichtioides_gastaldii:0.005378056,Gricetoides_aurora e:0.007715093):0.003557611,Eschrichtius_robustus:0.004637599):0.01195338):0.008826684,((Diunatans_luctoretemergo:0.01184207,Fragilicetus_velponi:0.008429416):0.004523488,(Megapt era_hubachi:0.006483132,Plesiobalaenoptera_quarantellii:0.010134):0.004545186):0.004642979):0.003512613):0.002210281):0.002584608,Balaenoptera_siberi:0.005978731):0.002536365):0.00172476):0.006396772,(Balaenoptera_ryani:0.009675268,Megaptera_miocaena:0.01340319):0.0 02770652):0.005903242,(Pelocetus_calvertensis:0.005532136,Uranocetus_gramensis:0.0060263 85):0.004464163):0.001989963,(((((((Aulocetus_latus:0.004459229,(Cetotherium_megalophysu m:0.003155441,Metopocetus_vandelli:0.002499759):0.0036299):0.002358038,((Metopocetus_d urinasus:0.001442301,Metopocetus_hunteri:0.002683932):0.01405799,Piscobalaena_nana:0.005 847589):0.005851826):0.00405425,(((Brandtocetus_chongulek:0.0031466,(Cetotherium_riabinin i:0.001756915,Kurdalagonus_mchedlidzei:0.003135206):0.00278539):0.002233289,Cetotherium_rathkii:0.009283864):0.01139562,Herentalia_nigra:0.01027044):0.004691926):0.003956701,(((Caperea_marginata:0.007141826,Miocaperea_pulchra:0.005644843):0.02058076,((((Herpetocetu s_bramblei:0.001381608,Herpetocetus_morrowi:0.003351012):0.001702554,(’Herpetocetus_sp.’: 0.001111989,NMNS_PV19540:0.001203288):0.002254647):0.00189362,Herpetocetus_transatla nticus:0.001406395):0.00499799,Nannocetus_eremus:0.003970936):0.009059672):0.006637674,Cephalotropis_coronatus:0.0103581):0.003733926):0.005978666,Joumocetus_shimizui:0.005847 192):0.002527393,Tiucetus_rosae:0.005597087):0.006958546,(Diorocetus_hiatus:0.001240824, Thinocetus_arthritus:0.006326648):0.004735413):0.005369853):0.00328242):0.002775609,((((Diorocetus_chichibuensis:0.004048948,(((Parietobalaena_campiniana:0.004030215,(’Parietobalaen a_sp.’:0.003300922,Tiphyocetus_temblorensis:0.002731098):0.002119772):0.00176437,(Parietob alaena_palmeri:0.001448439,Pinocetus_polonicus:0.006640526):0.001069989):0.002188237,Par ietobalaena_yamaokai:0.003028849):0.002109548):0.005645147,Isanacetus_laticephalus:0.0018 04021):0.004228741,OU_22705:0.004947175):0.003311185,Diorocetus_shobarensis:0.004415173):0.004321945):0.004670765,((((((Balaena_montalionis:0.003329911,(Balaena_mysticetus:0.0 08603945,Balaena_ricei:0.001018764):0.001789729):0.002812391,(Eubalaena_ianitrix:0.002575 507,(Eubalaena_shinshuensis:0.005514485,’Eubalaena_spp.’:0.00261434):0.005033338):0.005139279):0.002910737,(Balaenella_brachyrhynus:0.003154082,Balaenula_astensis:0.002143649):0. 002049175):0.001901428,’Balaenula_sp.’:0.00570575):0.006461453,Peripolocetus_vexillifer:0.0 05437092):0.002836467,Morenocetus_parvus:0.005001642):0.01627574):0.005412766,Titanocetus_sammarinensis:0.01052158):0.01413724):0.01306845,Horopeta_umarere:0.003403357):0.01 382415,OU_22224:0.02076156):0.005868653,Whakakai_waipata:0.007164906):0.009337821,(((Eomysticetus_whitmorei:0.005234376,Yamatocetus_canaliculatus:0.002985905):0.002380472,M icromysticetus_rothauseni:0.005414977):0.002938926,(Matapanui_waihao:0.001793079,(Tohora ata_raekohao:7.576104E-4,(Tokarahia_kauaeroa:0.001436352,Waharoa_ruwhenua:0.00107129):0.001231495):0.0057419 05):0.004962339):0.01134334):0.01879592):0.003120532,Morawanocetus_yabukii:0.003896515):0.007183905,(Janjucetus_hunderi:0.003141369,(Mammalodon_colliveri:0.001371781,Mammal odon_hakataramea:0.00148955):0.006998726):0.01190211):0.003541884,((Llanocetus_denticren atus:0.006107906,Mystacodon_selenensis:0.006542345):0.007707808,OU_GS10897:0.0082167 74):0.005255141):0.004323793,Coronodon_havensteini:0.01597085):0.005969355);

de Muizon et al. (2019): 43 taxa, 282 characters (274 parsimony informative)

Minimum tree length: 970

Number of minimum length trees: 2

Strict consensus in Newick format:

(Zygorhiza_kochii,Cynthiacetus_peruvianus,((Archaeodelphis_patrius,(Simocetus_rayii,(Waipati a_maerewhenua,(Squalodon_spp,(Tursiops_truncatus,Physeter_macrocephalus))))),(Mystacodon_selenensis,(((Llanocetus_denticrenatus,((Fucaia_goedertorum,((Aetiocetus_cotylalveus,Aetiocet us_polydentatus),Aetiocetus_weltoni)),((Micromysticetus_rothauseni,(Yamatocetus_canaliculatu s,Eomysticetus_whitmorei)),((Morenocetus_parvus,(Balaena_mysticetus,’Eubalaena_spp.’)),(((Ca perea_marginata,Miocaperea_pulchra),(Cetotherium_rathkii,Cetotherium_riabinini),Piscobalaena_nana,(Herpetocetus_morrowi,Herpetocetus_transatlanticus,Herpetocetus_bramblei)),(Diorocetus_hiatus,((Aglaocetus_patulus,(Aglaocetus_moreni,(Parietobalaena_campiniana,Parietobalaena_pa lmeri))),(Pelocetus_calvertensis,(Eschrichtius_robustus,(Megaptera_novaeangliae,(Balaenoptera_ acutorostrata,Balaenoptera_physalus))))))))))),((Coronodon_havensteini,ChM_PV5720),ChM_P V4745)),(Janjucetus_hunderi,Mammalodon_colliveri)))));

## Appendix 2. Characters related to the filter-feeding apparatus of mysticetes from 11 studies that were mapped onto phylogenetic hypotheses. The wording of each character description was taken directly from each publication

### Mandibular symphysis not sutured

Fitzgerald (2006): character 41 (ordered).
Mandibular symphysis.
0 = fused
1 = sutured but unfused
2 = not sutured, shallow longitudinal fossa for fibrocartilaginous connection
3 = not sutured, well-developed longitudinal groove for fibrocartilaginous connection
Note: character mapping for cladogram in Fig. 3a merges states 0 and 1, as well as states 2 and 3.

Deméré et al. (2008): character 53. Mandibular symphysis.
0 = sutured
1 = not sutured (ligamentous attachment in extant mysticetes).

Fitzgerald (2010): character 26. Mandibular symphysis.
0 = sutured but unfused
1 = not sutured, well-developed longitudinal fossa or groove for fibrocartilaginous connection Note: coded this character as binary which is simplified relative to multistate ordered in Fitzgerald (2006). Miscodings of *Mammalodon* and *Janjucetus* are corrected in Fitzgerald (2010) with change from ‘not sutured’ to ‘missing/ambiguous’.

Marx (2011): character 106
Mandibular symphysis.
0 = sutured or fused
1 = not sutured

Boessenecker and Fordyce (2015b): character 255
Mandible, symphysis.
0 = sutured or fused
1 = not sutured

Marx and Fordyce (2015): character 217.
Mandibular symphysis.
0 = sutured or fused
1 = unfused

Lambert et al. (2017): character 217.
Mandibular symphysis.
0 = sutured or fused
1= unfused

Geisler et al. (2017): character 49.
Mandibular symphysis.
0 = sutured,
1 = not sutured (ligamentous attachment in extant mysticetes)

Peredo et al. (2018): character 255.
Mandible, symphysis.
0 = sutured or fused
1 = not sutured

Fordyce and Marx (2018): character 217.
Mandibular symphysis.
0 = sutured or fused
1 = unsutured

de Muizon et al. (2019): character 222.
Mandibular symphysis.
0 = sutured or fused
1 = unfused

### Symphyseal groove (mandible)

Fitzgerald (2006): see above character 41.
Deméré et al. (2008): see character 53 (above).
Fitzgerald (2010): see character 26 (above).
Marx (2011): not coded in matrix.

Boessenecker and Fordyce (2015b): character 278.
Mandible, symphyseal groove.
0 = absent in adults
1 = prominent in adults

Marx and Fordyce (2015): not coded in matrix.
Lambert et al. (2017): not coded in matrix
Geisler et al. (2017): see character 49 (above).

Peredo et al. (2018): character 278
Mandible, symphyseal groove.
0 = absent in adults
1 = prominent in adults

Fordyce and Marx (2018): not coded in matrix.
de Muizon et al. (2019): not coded in matrix.

### Lateral bowing of mandibles

Fitzgerald (2006): character 43.
Mandible.
0 = bowed medially
1 = straight
2 = slightly bowed laterally, a line drawn from the posteriormost to anteriormost points stays within the body of mandible
3 = strongly bowed outward, line from anterior to posterior points does not entirely lie within body of mandible
Note: character mapping for cladogram in Fig. 3a merges states 1, 2, and 3.

Deméré et al. (2008): character 57 (ordered).
Mandible, curvature of ramus, in dorsal aspect.
0 = Laterally concave
1 = Straight
2 = Laterally convex
Note: character mapping for cladogram in Fig. 3b merges states 1 and 2.

Fitzgerald (2010): character 27.
Mandible.
0 = bowed medially
1 = straight
2 = slightly bowed laterally, a line drawn from the posteriormost to anteriormost points stays within the body of mandible
3 = strongly bowed outward, line from anterior to posterior points does not entirely lie within body of mandible
Note: character mapping for cladogram in Fig. 3c merges states 1-3.

Marx (2011): character 109.
Horizontal ramus of mandible in dorsal view.
0 = bowed medially
1 = straight or slightly bowed laterally, a line drawn from the posteriormost to anteriormost points stays within the body of the mandible
2 = strongly bowed outward, line from anterior to posterior points does not entirely lie within the body of the mandible
Note: character mapping for cladogram in Fig. 3d merges states 1 and 2.

Boessenecker and Fordyce (2015b): character 257.
Mandible, curvature of horizontal ramus in dorsal aspect.
0 = medially bowed
1 = straight or slightly bowed laterally, line connecting anterior and posterior tips stays within body of mandible
2 = strongly bowed laterally, line connecting anterior and posterior tips medial to ramus Note: character mapping for cladogram in Fig. 5a merges states 1 and 2.

Marx and Fordyce (2015): character 219 (ordered).
Mandibular body in dorsal view.
0 = bowed medially
1 = straight
2 = bowed laterally, but with curvature mainly confined to anterior portion of mandible
3 = evenly bowed laterally
Note: character mapping for cladogram in Fig. 5b merges states 1-3.

Lambert et al. (2017): character 219 (ordered).
Mandibular_body in dorsal view.
0 = bowed medially 1 = straight
2 = bowed laterally but with curvature mainly confined to anterior portion of mandible
3 = evenly bowed laterally
Note: character mapping for cladogram in Fig. 7a merges states 1-3.

Geisler et al. (2017): character 53 (ordered).
Mandible, curvature of ramus, in dorsal aspect.
0 = Laterally concave
1 = Straight
2 = Laterally convex a line drawn from the extremes stays within body of mandible
3 = Laterally convex that line at least partially medial to body of mandible.
Note: character mapping for cladogram in Fig. 7b merges states 1-3.

Peredo et al. (2018): character 257.
Mandible, curvature of horizontal ramus in dorsal aspect.
0 = medially bowed
1 = straight or slightly bowed laterally, line connecting anterior and posterior tips stays within body of mandible
2 = strongly bowed laterally, line connecting anterior and posterior tips medial to ramus
Note: character mapping for cladogram in Fig. 7c merges states 1 and 2.

Fordyce and Marx (2018): character 219 (ordered).
Mandibular body in dorsal view.
0 = bowed medially
1 = straight
2 = bowed laterally, but with curvature mainly confined to anterior portion of mandible 3 = evenly bowed laterally
Note: character mapping for cladogram in Fig. 8a merges states 1-3.

de Muizon et al. (2019): character 224 (ordered).
Mandibular body in dorsal view.
0 = bowed medially
1 = straight
2 = bowed laterally, but with curvature mainly confined to anterior portion of mandible
3 = evenly bowed laterally
Note: character mapping for cladogram in Fig. 8b merges states 1-3.

### Baleen present

Fitzgerald (2006): character 1.
Baleen.
0 = absent
1 = present

Deméré et al. (2008): character 71.
Baleen.
0 = absent
1 = present

Fitzgerald (2010): character 1.
Baleen.
0 = absent
1 = present
Note: no extant outgroups were coded in the matrix, so this character is not parsimony-informative.

Marx (2011): character 138.
Baleen.
0 = absent
1 = present

Boessenecker and Fordyce (2015b): character 346.
Soft tissue, baleen.
0 = absent
1 = present

Marx and Fordyce (2015): not coded in matrix.
Lambert et al. (2017): not coded in matrix.

Geisler et al. (2017): character 67. Baleen.
0 = absent
1 = present

Peredo et al. (2018): character 346. Soft tissue, baleen.
0 = absent
1 = present

Fordyce and Marx (2018): not coded in matrix.
de Muizon et al. (2019): not coded in matrix.

### Lateral palatal foramina and sulci

Fitzgerald (2006): character 18.
Palatal surface of maxilla.
0 = bears few vascular foramina, those that are present are small
1 = bears many, large vascular foramina that open laterally and anterolaterally into long sulci
2 = bears numerous small vascular foramina that lack sulci
Note: in Fig. 3a, state 2 that was coded only for *Physeter macrocephalus* is lighter shade of green; *Eubalaena glacialis* was coded state 0 (gray), unlike other extant mysticetes that were coded state 1 (darker green).

Deméré et al. (2008): character 38.
Lateral nutrient foramina and sulci on palate.
0 = absent
1 = present

Fitzgerald (2010): character 14.
Lateral nutrient foramina opening into sulci on the palatal surface of maxilla.
0 = absent
1 = present
Notes: coding is changed relative to Fitzgerald (2006). Fitzgerald (2010) simplified to a binary character and coded aetiocetids as ‘present’ and homologous to the state in most toothless baleen whales, in contrast to the prior study. *Eubalaena glacialis*, however, was again as ‘absent’ for lateral nutrient foramina.

Marx (2011): character 20.
Palatal nutrient foramina and sulci.
0 = absent
1 = present

Boessenecker and Fordyce (2015b): character 24.
Maxilla, nutrient foramina and sulci.
0 = absent
1 = present

Marx and Fordyce (2015): character 19.
Palatal nutrient foramina and sulci.
0 = absent
1 = present

Lambert et al. (2017): character 19.
Palatal nutrient foramina and sulci.
0 = absent
1 = present

Geisler et al. (2017): character 35.
Lateral nutrient foramina and sulci on palate.
0 = absent
1 = present

Peredo et al. (2018): character 24.
Maxilla, nutrient foramina and sulci.
0 = absent
1 = present

Fordyce and Marx (2018): character 18.
Palatal nutrient foramina and sulci.
0 = absent
1 = present

de Muizon et al. (2019): character 22.
Palatal nutrient foramina and sulci.
0 = absent
1 = present

### Postnatal teeth absent or vestigial

Fitzgerald (2006): character 38 (ordered).
Number of teeth in lower jaw.
0 = none
1 = 0, 1 tooth
2 = 0, 2 teeth
3 = 8-9 teeth
4 = 11-12 teeth
5 = 13-14 teeth
6 = 20-23 teeth
7 = 24-27 teeth
8 = 28-34 teeth
9 = more than 40 teeth
Notes: character mapping for cladogram in Fig. 3a merges states 1-9, all of which indicate the presence of postnatal mineralized teeth. Fitzgerald (2006) also coded character 25 that describes variation in the number of upper teeth in maxilla.

Deméré et al. (2008): character 61.
Mineralized teeth in adults.
0 = present
1 = absent.

Fitzgerald (2010): character 25 (ordered).
Number of teeth in lower jaw.
0 = none
1 = 1 tooth
2 = 2 teeth
3 = 8-9 teeth
4 = 11 teeth
5 = 12 teeth
6 = 13-15 teeth
7 = 20-23 teeth
8 = 24-27 teeth
9 = more than 28 teeth
Notes: character mapping for cladogram in Fig. 3a merges states 1-9, all of which indicate the presence of postnatal mineralized teeth. Fitzgerald (2006) also coded character 17 that describes variation in the number of upper teeth in maxilla.

Marx (2011): character 25
Teeth in adult individuals.
0 = present
1 = absent

Boessenecker and Fordyce (2015b): character 281.
Dentition, teeth.
0 = present in adult
1 = absent in adult

Marx and Fordyce (2015): character 26.
Teeth in adult individuals.
0 = present
1 = absent or vestigial

Lambert et al. (2017): character 26.
Teeth in adult individuals.
0 = present
1 = absent or vestigial

Geisler et al. (2017): character 57.
Mineralized teeth in adults.
0 = present
1 = absent

Peredo et al. (2018): character 281.
Dentition, teeth.
0 = present in adult
1 = absent in adult

Fordyce and Marx (2018): character 25.
Teeth in adult individuals.
0 = present
1 = absent or vestigial

de Muizon et al. (2019): character 28.
Teeth in adult individuals
0 = present
1 = absent or vestigial

## References

Árnason Ú, Lammers F, Kumar V, Nilsson MA, Janke A (2018) Whole-genome sequencing of the blue whale and other rorquals finds signatures for introgressive gene flow. Sci Adv 4:eaap9873

Barnes LG, Kimura M, Furusawa H, Sawamura H (1994) Classification and distribution of Oligocene Aetiocetidae (Mammalia; Cetacea; Mysticeti) from western North America and Japan. The Island Arc 3:392–431. doi: 10.1111/j.1440-1738.1994.tb00122.x

Bartlett JD, Ganss B, Goldberg M, Moradian**-**Oldak J, Paine ML, Snead ML, Wen X, White SN, Zhou YL (2006) Protein–protein interactions of the developing enamel matrix. Curr Top Dev Biol 74:57–115

Bartlett JD, Simmer JP (2015) New perspectives on amelotin and amelogenesis. J Dent Res 94:642–644

Berta A, Ekdale EG, Zellmer NT, Deméré, TA, Kienle SS, Smallcomb M (2015) Eye, nose, hair, and throat: external anatomy of the head of a neonate gray whale (Cetacea, Mysticeti, Eschrichtiidae). Anat Rec 298:648–659

Berta A, Lanzetti A, Ekdale EG, Deméré TA (2016) From teeth to baleen and raptorial to bulk filter feeding in mysticete cetaceans: the role of paleontological, genetic, and geochemical data in feeding evolution and ecology. Integrative and Comparative Biology 56:1271–1284

Bisconti M (2012) Comparative osteology and phylogenetic relationships of *Miocaperea pulchra*, the first fossil pygmy right whale genus and species (Cetacea, Mysticeti, Neobalaenidae). Zool J Linn Soc 166:876–911. doi: 10.1111/j.1096-3642.2012.00862.x

Boessenecker RW, Fordyce RE (2015a) Anatomy, feeding ecology, and ontogeny of a transitional baleen whale: a new genus and species of Eomysticetidae (Mammalia: Cetacea) from the Oligocene of New Zealand. PeerJ 3:e1129. doi: 10.7717/peerj.1129

Boessenecker RW, Fordyce RE (2015b) A new genus and species of eomysticetid (Cetacea: Mysticeti) and a reinterpretation of ‘*Mauicetus lophocephalus*’ Marples, 1956: transitional baleen whales from the upper Oligocene of New Zealand. Zool. J Linn Soc 175:607–660. doi: 10.1111/zoj.12297

Boessenecker RW, Fordyce RE (2017) A new eomysticetid from the Oligocene Kokoamu Greensand of New Zealand and a review of the Eomysticetidae (Mammalia, Cetacea). J Syst Palaeontol 15:429–469. doi: 10.1080/14772019.2016.1191045

Bragulla HH, Homberger DG (2009) Structure and functions of keratin proteins in simple, keratinizing or cornifying epithelia. J Anat 214:516–559

Brodie PF (2001) Feeding mechanics of rorquals *Balaenoptera* sp. In: Mazin J, de Buffrenil V (eds) Secondary Adaptations of Tetrapods to Life in Water. Verlag, Munchen, pp. 345–352

Brower AVZ, Schawaroch V (1996) Three steps of homology assessment. Cladistics 12:265–272

Brusatte SL, O’Connor JK, Jarvis ED (2015) The origin and diversification of birds. Curr Biol 25:R888–R898

Cobourne MT, Sharpe PT (2010) Making up the numbers: the molecular control of mammalian dental formula. Semin Cell Dev Biol 21: 314–324

Clementz MT, Fordyce RE, Peek SL, Fox DL (2014) Ancient marine isoscapes and isotopic evidence of bulk-feeding by Oligocene cetaceans. Palaeogeogr Palaeoclimatol Palaeoecol 400:28–40

Darwin C (1872) The origin of species by means of natural selection, or the preservation of favoured races in the struggle for life, 6^th^ edition. John Murray, London

Deméré TA (2005) Palate vascularization in an Oligocene toothed mysticete (Cetacea: Mysticeti: Aetiocetidae); implications for the evolution of baleen. Cranbrook Inst Sci Misc Publ 1:21

Deméré TA, Berta A (2008) Skull anatomy of the Oligocene toothed mysticete *Aetiocetus weltoni* (Mammalia; Cetacea): implications for mysticete evolution and functional anatomy. Zool J Linn Soc 154:308–352. doi: 10.1111/j.1096-3642.2008.00414.x

Deméré TA, McGowen MR, Berta A, Gatesy J (2008) Morphological and molecular evidence for a stepwise evolutionary transition from teeth to baleen in mysticete whales. Syst Biol 57:15–37. doi: 10.1080/10635150701884632

de Muizon C, Bianucci G, Martínez-Cáceres M, Lambert O (2019) *Mystacodon selenensis*, the earliest known toothed mysticete (Cetacea, Mammalia) from the late Eocene of Peru: Anatomy, phylogeny, and feeding adaptations. Geodiversitas 41:401–499. doi:10.5252/geodiversitas2019v41a11. http://geodiversitas.com/41/11

de Pinna M (1991) Concepts and tests of homology in the cladistic paradigm. Cladistics 7:367– 394

de Queiroz A, Gatesy J (2007) The supermatrix approach to systematics. Trends Ecol Evol 22:34–41

Drake SE, Crish SD, George JC, Stimmelmayr R, Thewissen JG (2015) Sensory hairs in the bowhead whale, *Balaena mysticetus* (Cetacea, Mammalia). Anat Rec 298:1327–1335

Ehrlich F, Fischer H, Langbein L, Praetzel-Wunder S, Ebner B, Figlak K, Weissenbacher A, Sipos W, Tschachler E, Eckhart L (2019). Differential evolution of the epidermal keratin cytoskeleton in terrestrial and aquatic mammals Mol Biol Evol 36:328–340

Ekdale EG, Deméré TA (2022) Neurovascular evidence for a co-occurrence of teeth and baleen in an Oligocene mysticete and the transition to filter feeding in baleen whales. Zool J Linn Soc 194:395–415. doi: 10.1093/zoolinnean/zlab017

Ekdale EG, Deméré TA, Berta A (2015) Vascularization of the gray whale palate (Cetacea, Mysticeti, *Eschrichtius robustus*): soft tissue evidence for an alveolar source of blood to baleen. Anat Rec 298:691–702. doi: 10.1002/ar.23119

Espregueira Themudo G, Alves LQ, Machado AM, Lopes-Marques M, da Fonseca RR, Fonseca M, Ruivo R, Castro LFC (2020) Losing genes: the evolutionary remodeling of Cetacea skin. Front Mar Sci 7:592735. doi: 10.3389/fmars.2020.592375

Fink WL (1982) The conceptual relationship between ontogeny and phylogeny. Paleobiology 8:254–264

Fitzgerald EMG (2006) A bizarre new toothed mysticete (Cetacea) from Australia and the early evolution of baleen whales. Proc Royal Soc B 273:2955–2963

Fitzgerald EMG (2010) The morphology and systematics of *Mammalodon colliveri* (Cetacea: Mysticeti), a toothed mysticete from the Oligocene of Australia. Zool J Linn Soc-Lond 158: 367– 476

Fitzgerald EMG (2012) Archaeocete-like jaws in a baleen whale. Biol Lett 8:94–96. doi: 10.1098/rsbl.2011.0690

Fordyce RE (1984) Evolution and zoogeography of cetaceans in Australia. In: Archer M, Clayton G (eds) Vertebrate Zoogeography and Evolution in Australasia. Hesperian Press, Perth, pp. 929– 948

Fordyce RE, de Muizon C (2001) Evolutionary history of cetaceans: a review. In: Mazin J, de Buffrenil V (eds) Secondary Adaptations of Tetrapods to Life in Water. Verlag, Munchen, pp. 169–233

Fordyce RE, Marx FG (2018) Gigantism precedes filter feeding in baleen whale evolution. Curr Biol 28:1670–1676. doi: 10.1016/j.cub.2018.04.027

Fudge DS, Szewciw LJ, Schwalb AN (2009) Morphology and development of blue whale baleen: an annotated translation of Tycho Tullberg’s classic 1883 paper. Aquat Mamm 35:226–52

Gaskin DE (1982) The Ecology of Whales and Dolphins. Heinemann, Portsmouth, New Hampshire, USA

Gasse B, Liu X, Corre E, Sire J-Y (2015) Amelotin gene structure and expression during enamel formation in the opossum *Monodelphis domestica*. PLoS ONE 10:e0133314

Gatesy J, Geisler JH, Chang J, Buell C, Berta A, Meredith RW, Springer MS, McGowen MR (2013) A phylogenetic blueprint for a modern whale. Mol Phylogenet Evol 66:479–506

Geisler J.H, Sanders AE (2003) Morphological evidence for the phylogeny of Cetacea. J Mamm Evol 10:23–129

Geisler JH, Boessenecker RW, Brown M, Beatty B (2017) The origin of filter feeding in whales. Curr Biol 27:2036–2042. doi: 10.1016/j.cub.2017.06.003

Goldbogen JA, Pyenson, ND, Shadwick, RE (2007) Big gulps require high drag for fin whale lunge feeding. Mar Ecol Prog Ser 349:289–301

Goldbogen JA, Cade DE, Calambokidis J, Friedlaender AS, Potvin J, Segre PS Werth AJ (2017) How baleen whales feed: the biomechanics of engulfment and filtration. Annu Rev Mar Sci 9:367–386

Goldbogen JA, Madsen PT (2018) The evolution of foraging capacity and gigantism in cetaceans. J Exp Biol 221:jeb166033. doi: 10.1242/jeb.166033

Goloboff, PA (1993) Estimating character weights during tree search. Cladistics 9:83–91

Gould SJ (1970) Dollo on Dollo’s law: irreversibility and the status of evolutionary laws. J Hist Biol 3:189–212

Gould SJ, Vrba ES (1982) Exaptation – a missing term in the science of form. Paleobiology 8:4– 15

Greenwold MJ, Sawyer RH (2010) Genomic organization and molecular phylogenies of the beta (β) keratin multigene family in the chicken (*Gallus gallus*) and zebra finch (*Taeniopygia guttata*): implications for feather evolution. BMC Evol Biol 10:148

Hesse M, Zimek A, Weber K, Magin TM (2004) Comprehensive analysis of keratin gene clusters in humans and rodents. Eur J Cell Biol 83:19–26

Hocking DP, Marx FG, Park T, Fitzgerald EMG, Evans AR (2017a) A behavioural framework for the evolution of feeding in predatory aquatic mammals. Proc Royal Soc B 284. doi: 10.1098/rspb.2016.2750

Hocking DP, Marx FG, Fitzgerald EMG, Evans AR (2017b) Ancient whales did not filter feed with their teeth. Biol Lett 13:20170348. http://dx.doi.org/10.1098/rsbl.2017.0348

Hytönen MK, Arumilli M, Sarkiala E, Nieminen P, Lohi H (2019) Canine models of human amelogenesis imperfecta: identification of novel recessive *ENAM* and *ACP4* variants. Hum Genet 138:525–533

Ichishima H, Sawamura H, Ito H, Otani S, Ishikawa H (2008) Do the so-called nutrient foramina on the palate tell us the presence of baleen plates in toothed mysticetes? In: Abstracts of the Fifth Conference on Secondary Adaptation of Tetrapods to Life in Water, 9–13th June 2008. National Museum of Nature and Science, Tokyo, pp. 24–25

Ishikawa H, Amasaki H (1995) Development and physiological degradation of tooth buds and development of rudiment of baleen plate in southern minke whale, *Balaenoptera acutorostrata*. J Vet Med Sci 57:665–670. doi: 10.1248/cpb.37.3229

Jackson JA, Baker CS, Vant M, Steel DJ, Medrano-Gonzalez, L, Palumbi SR (2009) Big and slow: phylogenetic estimates of molecular evolution in baleen whales (suborder Mysticeti). Mol Biol Evol 26:2427–2440. doi:10.1093/molbev/msp169

Karlsen K (1962) Development of tooth germs and adjacent structures in the whalebone whale (*Balaenoptera physalus*). Hval skrif 45:5–56

Katoh, K, Misawa K, Kuma K, Miyata T (2002) MAFFT: a novel method for rapid multiple sequence alignment based on fast Fourier transform. Nucleic Acids Res 30:3059–3066

Kawasaki K, Hu JC-C, Simmer JP (2014) Evolution of *Klk4* and enamel maturation in eutherians. Biol Chem 395:1003–1013

Kawasaki K, Mikami M, Goto M, Shindo J, Amano M, Ishiyama M (2020) The evolution of unusually small amelogenin genes in cetaceans; pseudogenization, X–Y gene conversion, and feeding strategy. J Mol Evol 88:122–135. doi.org/10.1007/s00239-019-09917-0

Keane M, Semeiks J, Webb AE, Li YI, Quesada V, Craig T, Madsen LB, van Dam S, Brawand D, Marques PI, Michalak P, Kang L, Bhak J, Yim HS, Grishin NV, Nielsen NH, Heide-Jorgensen MP, Oziolor EM, Matson CW, Church GM, Stuart GW, Patton JC, George JC, Suydam R, Larsen K, Lopez-Otin C, O’Connell MJ, Bickham JW, Thomsen B, de Magalhaes JP (2015) Insights into the evolution of longevity from the bowhead whale genome. Cell Rep 10:112–120

Kearse M, Moir R, Wilson A, Stones-Havas S, Cheung M, Sturrock S, Buxton S, Cooper A, Markowitz S, Duran C, Thierer T, Ashton B, Mentjies P, Drummond A (2012) Geneious basic: an integrated and extendable desktop software platform for the organization and analysis of sequence data. Bioinformatics 28:1647–1649

Kellogg R (1965) A new whalebone whale from the Miocene Calvert Formation. Bull US Natl Mus 247:1–63

Khan I, Maldonado E, Vasconcelos V, O’Brien SJ, Johnson WE, Antunes (2014) Mammalian keratin associated proteins (KRTAPs) subgenomes: disentangling hair diversity and adaptation to terrestrial and aquatic environments. BMC Genomics 15:779

Kienle SS, Law CJ, Costa DP, Berta A, Mehta R (2017) Revisiting the behavioural framework of feeding in predatory aquatic mammals. Proc Royal Soc B 284: 20171035

Lambert O, Martínez-Cáceres M, Bianucci G, Di Celma C, RSalas-Gismondi R, Steurbaut E, Urbina M, de Muizon C (2017) Earliest mysticete from the Late Eocene of Peru sheds light on the origin of baleen whales. Curr Biol 27:1535–1541

Lambertsen RH (1983) Internal mechanism of rorqual feeding. J Mammal 64:76–88

Lambertsen R, Ulrich N, Straley J (1995) Frontomandibular stay of Balaenopteridae: a mechanism for momentum recapture during feeding. J Mammal 76:877–899. doi: 10.2307/1382758

Lambertsen R, Rasmussen K, Lancaster WC, Hintz R (2005) Functional morphology of the mouth of the bowhead whale and its implications for conservation. J Mammal 86:342–352

Lanzetti A (2019) Prenatal developmental sequence of the skull of minke whales and its implications for the evolution of mysticetes and the teeth-to-baleen transition. J Anat 235:725– 748

Lanzetti A, Berta A, Ekdale EG (2020) Prenatal development of the humpback whale: growth rate, tooth loss and skull shape changes in an evolutionary framework. Anat Rec 303:180–204. doi: 10.1002/ar.23990

Li YI, Kong LS, Ponting CP, Haerty W (2013) Rapid evolution of beta-keratin genes contribute to phenotypic differences that distinguish turtles and birds from other reptiles. Genome Biol Evol 5:923–933

Liang T, Hu Y, Kawasaki K, Zhang H, Zhang C, Saunders TL, Simmer JP, Hu JC-C (2021) Odontogenesis-associated phosphoprotein truncation blocks ameloblast transition into maturation in OdaphC41*/C41* mice. Sci Rep 11:1132

Ling JK (1977) Vibrissae of marine mammals. In: Harrison RJ (ed) Functional Anatomy of Marine Mammals. Academic Press, London, pp. 387–415

Loch C, Buono M, Kalthoff D, Mörs T, Fernández M (2020) Enamel microstructure in Eocene cetaceans from Antarctica (Archaeoceti and Mysticeti). J Mamm Evol 27:289–298. doi.org/10.1007/s10914-018-09456-3

Lowe CB, Clarke JA, Baker AJ, Haussler D, Edwards SV (2015) Feather development genes and associated regulatory innovation predate the origin of Dinosauria. Mol Biol Evol 32:23–38

Mabee PM (1993) Phylogenetic interpretation of ontogenetic change: sorting out the actual and artefactual in an empirical case study of centrarchid fishes. Zool J Linnean Soc 107:175–291

Marx FG (2011) The more the merrier? A large cladistic analysis of mysticetes, and comments on the transition from teeth to baleen. J Mammal Evol 18:77–100

Marx FG, Fordyce RE (2015) Baleen boom and bust: a synthesis of mysticete phylogeny, diversity and disparity. R Soc Open Sci 2:140434. doi: 10.1098/rsos.140434

Marx FG, Kohno N (2016) A new Miocene baleen whale from the Peruvian desert. R Soc Open Sci 3:160542. doi: 10.1098/rsos.160542

Marx FG, Tsai C-h, Fordyce RE (2015) A new Early Oligocene toothed ‘baleen’ whale (Mysticeti: Aetiocetidae) from western North America: one of the oldest and the smallest. R Soc Open Sci 2:150476. doi: 10.1098/rsos.150476

Marx FG, Hocking DP, Park T, Ziegler T, Evans AR, Fitzgerald EMG (2016) Suction feeding preceded filtering in baleen whale evolution. Mem Mus Vic 75:71–82

Marx FG, Collaretta A, Gioncada A, Post K, Lambert O, Bonaccorsi E, Urbina M, Bianucci G (2017) How whales used to filter: exceptionally preserved baleen in a Miocene cetotheriid. J Anat 231:212–220. doi: 10.1111/joa.12622

McGowen MR, Spaulding M, Gatesy J (2009) Divergence date estimation and a comprehensive molecular tree of extant cetaceans. Mol Phylogenet Evol 53:891–906. doi:10.1016/j.ympev.2009.08.018

McKnight DA, Fisher LW (2009) Molecular evolution of dentin phosphoprotein among toothed and toothless animals. BMC Evol Biol 9:299

Meredith RW, Gatesy J, Murphy WJ, Ryder OA, Springer MS (2009) Molecular decay of the tooth gene *Enamelin* (*ENAM*) mirrors the loss of enamel in the fossil record of placental mammals. PLoS Genet 5:e1000634

Meredith RW, Gatesy J, Cheng J, Springer MS (2011) Pseudogenization of the tooth gene enamelysin (*MMP20*) in the common ancestor of extant baleen whales. Proc Royal Soc B 278:993–1002

Meredith RW, Gatesy J, Springer MS (2013) Molecular decay of enamel matrix protein genes in turtles and other edentulous amniotes. BMC Evol Biol 13:20

Meredith RW, Zhang G, Gilbert MTP, Jarvis ED, Springer MS (2014) Evidence for a single loss of mineralized teeth in the common avian ancestor. Science 346:1254390–1254390

Miller GS (1929) The gums of the porpoise *Phocoenoides dalli* (True). Proc US Natl Mus 74:1–4

Mitchell ED (1989) A new cetacean from the Late Eocene La Meseta Formation, Seymour Island, Antarctic Peninsula. Can J Fish Aquat Sci 46:2219–2235

Mu Y, Huang X, Liu R, Gai Y, Liang N, Yin D, Shan L, Xu S, Yang G (2021) *ACPT* gene is inactivated in mammalian lineages that lack enamel or teeth. PeerJ 9:e10219

Mynett N, Mossman HL, Huettner T, Grant RA (2022). Diversity of vibrissal follicle anatomy in cetaceans. Anat Rec 305:609–621 https://doi.org/10.1002/ar.24714

Nery MF, Arroyo JI, Opazo JC (2014) Increased rate of hair keratin gene loss in the cetacean lineage. BMC Genomics 15:869

Nylander JAA, Ronquist F, Huelsenbeck JP, Nieves-Aldrey JL (2004) Bayesian phylogenetic analysis of combined data. Syst Biol 53:47–67

Oh JW, Chung O, Cho YS, MacGregor GR, Plikus MV (2015). Gene loss in keratinization programs accompanies adaptation of cetacean skin to aquatic lifestyle. Exp Dermatol 24:572–573

Okazaki Y (2012) A new mysticete from the upper Oligocene Ashiya Group, Kyushu, Japan, and its significance to mysticete evolution. Bulletin of the Kitakyushu Museum of Natural History and Human History Series A (Natural History) 10:129–152

Orton LS, Brodie PF (1987) Engulfing mechanics of fin whales. Can J Zool 65:2898–2907

Patterson C (1982) Morphological characters and homology. In: Joysey KA, Friday AE (eds) Problems of Phylogenetic Reconstruction. Academic Press, London. pp. 21–74

Patterson C (1988) Homology in classical and molecular biology. Mol Biol Evol 5:603–625

Peredo CM, Uhen MD (2016) A new basal chaeomysticete (Mammalia: Cetacea) from the Late Oligocene Pysht Formation of Washington, USA. Pap Palaeont 2:533–554. doi: 10.1002/spp2.1051

Peredo CM, Pyenson, ND (2018). *Salishicetus meadi*, a new aetiocetid from the late Oligocene of Washington State and implications for feeding transitions in early mysticete evolution. R Soc Open Sci 5:172336. doi: 10.1098/rsos.172336

Peredo CM, Pyenson ND, Boersma AT (2017) Decoupling tooth loss from the evolution of baleen in whales. Front Mar Sci 4:67. doi: 10.3389/fmars.2017.00067

Peredo CM, Pyenson ND, Marshall CD, Uhen MD (2018) Tooth loss precedes the origin of baleen in whales. Curr Biol 28:3992–4000. doi: 10.1016/j.cub.2018.10.047

Pilleri G (1989) *Balaenoptera siberi*, ein neuer spätmiozäner bartenwal aus der Pisco-formation Perus. In: Pilleri G (ed) Beiträge zur Paläontologie der Cetaceen Perus, Hirnanatomisches Institut Ostermundingen, Bern, pp. 65–84

Pivorunas A (1977) The fibrocartilage skeleton and related structures of the ventral pouch of balaenopterid whales. J Morphol 151:299–314

Pivorunas A (1979) The feeding mechanisms of baleen whales. Am Scient 67:432– 440

Pyenson ND, Goldbogen JA, Vogl AW, Szathmary G, Drake RL, Shadwick RE (2012) Discovery of a sensory organ that coordinates lunge feeding in rorqual whales. Nature 485:498–501. doi: 0.1038/nature11135

Ridewood WG (1923) Observations on the skull in foetal specimens of whales of the genera *Megaptera* and *Balaenoptera*. Phil Trans R Soc Lond B 211:209–272

Sanders AE, Barnes LG (2002) Paleontology of the Late Oligocene Ashley and Chandler Bridge Formations of South Carolina, 3: Eomysticetidae, a new family of primitive mysticetes (Mammalia, Cetacea). Smithsonian Contrib Paleobiol 93:313–356

Savoca MS, Czapanskiy MF, Kahane-Rapport SR, Gough WT, Fahlbusch JA, Bierlich KC, Segre PS, Di Clemente J, Penry GS, Wiley DN, Calambokidis J (2021) Baleen whale prey consumption based on high-resolution foraging measurements. Nature 599:85–90

Sawamura H (2008a) The origin of baleen whale–comparative morphology of the toothed mysticetes and the minke whale fetuses. J Fos Res 40:120–130

Sawamura H (2008b) Progress of the research on the toothed Mysticeti, AMP 14. Bull Ashoro Mus of Paleontol 5:23–40

Schweizer J, Bowden PE, Coulombe PA, Langbein L, Lane EB, Magin TM, Maltais L, Omary MB, Parry DAD, Rogers MA, Wright MW (2006) New consensus nomenclature for mammalian keratins. J Cell Biol 174:169–174

Shadwick RE, Potvin J Goldbogen JA (2019) Lunge feeding in rorqual whales. Physiology 4: 409–418

Sharma V, Hecker N, Roscito JG, Foerster L, Langer BE, Hiller M (2018) A genomics approach reveals insights into the importance of gene losses for mammalian adaptations. Nat Comm 9:1215

Slater GJ, Goldbogen JA, Pyenson ND (2017) Independent evolution of baleen whale gigantism linked to Plio-Pleistocene ocean dynamics. Proc Royal Soc B 284:20170546. doi.org/10.1098/rspb.2017.0546

Slijper EJ (1976) Whales and Dolphins. University of Michigan, Ann Arbor

Slijper EJ (1979) Whales, 2nd edition. Hutchinson, London

Smith KK (2001) Heterochrony revisited: the evolution of developmental sequences. Biol J Linn Soc 73:169–186

Solazzo C, Wadsley M, Dyer JM, Clerens S, Collins MJ, Plowman J (2013) Characterisation of novel α-keratin peptide markers for species identification in keratinous tissues using mass spectrometry. Rapid Commun Mass Spectrom 27:2685–98. doi: 10.1002/rcm.6730 PMID: 24591030

Solazzo C, Fitzhugh W, Kaplan S, Potter C, Dyer JM (2017) Molecular markers in keratins from Mysticeti whales for species identification of baleen in museum and archaeological collections. PLoS ONE 12(8): e0183053. doi:10.1371/journal.pone.0183053

Solis-Añorve A, González-Barba G, Hernández-Rivera R (2019) Description of a new toothed mysticete from the Late Oligocene of San Juan de La Costa, B.C.S., México. J South Am Earth Sci 89:337–346

Springer MS, Signore AV, Paijmans JLA, Vélez-Juarbe J, Domnin DP, Bauer CE, He K, Crerar L, Campos PF, Murphy WJ, Meredith RW, Gatesy J, Willerslev E, MacPhee RDE, Hofreiter M, Campbell KL (2015) Interordinal gene capture, the phylogenetic position of Steller’s sea cow based on molecular and morphological data, and the macroevolutionary history of Sirenia. Mol Phylogenet Evol 91:178–193

Springer MS, Starrett J, Morin PA, Lanzetti A, Hayashi C, Gatesy J (2016) Inactivation of *C4orf26* in toothless placental mammals. Mol Phylogenet Evol 95:34–45

Springer MS, Emerling CA, Gatesy J, Randall J, Collin MA, Hecker N, Hiller M, Delsuc F (2019) Odontogenic ameloblast-associated (*ODAM*) is inactivated in toothless/enamelless placental mammals and toothed whales. BMC Evol Biol 19:31

Springer MS, Guerrero-Juarez CF, Huelsmann M, Collin MA, Danil K, McGowen MR, Oh JW, Ramos R, Hiller M, Plikus MV, Gatesy J (2021) Genomic and anatomical comparisons of skin support independent adaptation to life in water by cetaceans and hippos. Curr. Biol. 31:2124– 2139. doi.org/10.1016/j.cub.2021.02.057

Steeman ME (2007) Cladistic analysis and a revised classification of fossil and recent mysticetes. Zool J Linn Soc 150:875–894

Steeman ME, Hebsgaard MB, Fordyce RE, Ho SYW, Rabosky DL, Nielsen R, Rahbek C, Glenner H, Sørensen MV, Willerslev E (2009) Radiation of extant cetaceans driven by restructuring of the oceans. Syst Biol 58:573–585

Sun X, Zhang Z, Sun Y, Li J, Xu S, Yang G (2017) Comparative genomics analyses of alpha-keratins reveal insights into evolutionary adaptation of marine mammals. Front Zool 14:4. doi: 10.1186/s12983-017-0225-x

Szewciw LJ, de Kerckhove DG, Grime GW, Fudge DS (2010) Calcification provides mechanical reinforcement to whale baleen α-keratin. Proc Royal Soc B 277:2597–605. doi: 10.1098/rspb.2010.0399 PMID: 20392736

Swofford DL (2002) PAUP*. Phylogenetic Analysis Using Parsimony (*and Other Methods). Sinauer Associates, Sunderland

Thesleff I (2006) The genetic basis of tooth development and dental defects. Am J Med Genet Part A 140A:2530–2535

Thewissen JGM, Hieronymus TL, George JC, Suydam R, Stimmelmayr R, McBurney D (2017) Evolutionary aspects of the development of teeth and baleen in the bowhead whale. J Anat 230:549–566. doi: 10.1111/joa.12579

Utrecht WLV (1965) On the growth of the baleen plate of the fin whale and the blue whale. Bijdr Dierk 35:3–38

Wenzel JW (1993) Behavioral homology and phylogeny. Ann Rev Ecol Syst 23:361–381

Werth AJ (2000) Feeding in marine mammals. In: Schwenk K (ed) Feeding. Academic, San Diego, pp. 487–526

Werth AJ, Potvin J, Shadwick RE, Jensen MM, Cade DE, Goldbogen, JA (2018) Filtration area scaling and evolution in mysticetes: trophic niche partitioning and the curious case of sei and pygmy right whales, Biol J Linn Soc 154:308–352. doi: 10.1111/j.1096-3642.2008.00414.x

Werth, AJ, Sformo TL, Lysiak NS, Rita D, George JC (2020) Baleen turnover and gut transit in mysticete whales and its environmental implications. Polar Biol 43:707–723. Doi.org/10.1007/s00300-020-02673-8

Wu DD, Irwin DM, Zhang YP (2008) Molecular evolution of the keratin associated protein gene family in mammals, role in the evolution of mammalian hair. BMC Evol Biol 8:241. doi:10.1186/1471-2148-8-241

Young S, Deméré TA, Ekdale EG, Berta A, Zellmer N (2015) Morphometrics and structure of complete baleen racks in gray whales (*Eschrichtius robustus*) from the Eastern North Pacific Ocean. Anat Rec 298: 703–719

